# Morphometric Identification of Parvalbumin-Positive Interneurons: A Data-Driven Approach

**DOI:** 10.1101/2024.12.27.630528

**Authors:** Maheshwar Panday, Leanne Monteiro, Ahad Daudi, Kathryn M. Murphy

**Affiliations:** McMaster Neuroscience Graduate Program, McMaster University, Hamilton, ON, L8S 4K1, Canada; Department of Psychology, Neuroscience & Behaviour, McMaster University, Hamilton, ON, L8S 4K1, Canada

**Author notes:** Correspondence should be addressed to Kathryn M. Murphy;, Kathryn Murphy, Department of Psychology, Neuroscience & Behavior, McMaster University, McMaster University, 1280 Main Street West, Hamilton, ON L8S 4K1, Canada.

## Abstract

Traditionally, anatomical studies of parvalbumin-positive (PV+) labelled interneurons describe them as a homogeneous population of neurons. In contrast, recent single-cell RNAseq studies have identified multiple transcriptomically distinct categories of PV+ cells. That difference between a single anatomical category of PV+ neurons and multiple transcriptomic categories presents a problem in understanding the role of these neurons in cortical function. One gap that might contribute to this discrepancy is that PV+ morphology is typically addressed using qualitative descriptions and simple quantifications, while single-cell RNAseq studies use big data and high dimensional analyses. PV+ neurons play critical roles in the experience-dependent development of the cortex and are often involved in disease-related changes associated with neurodegenerative and neuropsychiatric disorders. Here, we developed a modern data-driven analysis pipeline to quantify PV+ morphology. We quantified 97 morphometric features from 14274 PV+ neurons and applied unsupervised clustering that identified 13 different PV+ morphologies. We extended the analysis to compare PV+ dendritic arbour patterns and cell body morphologies. Finally, we compared the morphologies of PV+ neurons with the cell body morphologies of neurons expressing various genes associated with PV+ transcriptomic cell types. This approach identified a range of PV+ morphologies similar to the number of transcriptomic categories. It also found that the PV+ morphologies have cortical area, laminar, and transcriptomic biases that might contribute to cortical function.

## Introduction

Parvalbumin-positive (PV+) interneurons are the brain’s most abundant subclass of GABAergic interneurons. Recent single-cell RNAseq studies have identified multiple sub-categories of molecularly distinct PV+ neurons in the cortex. In contrast, PV+ interneurons are often categorized anatomically as a homogeneous group of neurons based on their immunoreactivity to the PV protein (Celio and Heizmann, 1981; Brederode et al., 1990; Rio et al., 1994) despite their unique and extensive dendritic and axonal arbour morphology differences (i.e., basket and chandelier cells) (Lewis and Lund, 1990; Wang et al., 2002; Jiang et al., 2015; Galarreta and Hestrin, 2002). This gap between the single-cell RNA and morphological definitions of this important inhibitory interneuron class is partly due to the qualitative and semi-quantitative approaches used to study the morphology and anatomical location of PV+ neurons. PV+ neurons are intimately involved in critical period closure, spike-time-dependent plasticity, and regulation of the excitatory-inhibitory balance (Levelt and Hübener, 2012). Furthermore, changes to PV+ neurons are linked to a variety of neurological and psychiatric disorders, including epilepsy, schizophrenia, autism as well as neurodegenerative disorders (Wöhr et al., 2015; Janickova et al., 2020; Ruden et al., 2021; Leitch, 2024). Here, we present a new approach to studying PV+ morphology that uses multiple measurements of morphological features and high-dimensional analyses to quantitatively categorize the organization of these neurons in the mouse somatosensory and visual cortex.

PV+ interneurons are implicated in a broad range of complex processes. As fast-spiking interneurons, they deliver and maintain highly temporally resolved inhibitory tone to pyramidal cells. In this way, PV+ interneurons act as cellular timekeepers, gating the flow of information through cortical circuits (Cardin et al., 2009). Forming elaborate networks of synaptic connections, PV+ interneurons also control the synchronous firing of excitatory neurons and are necessary for setting and maintaining gamma oscillations across larger cortical domains (Sohal et al., 2009; Levelt and Hübener, 2012). This establishes a centrally important role for PV+ cells in modulating sensory cortical circuitry in response to experience. Structurally, PV+ interneurons influence plasticity through structural interactions through both myelin and associations with the extracellular matrix. Myelin surrounding cortical PV+ interneurons forms shorter, “patchy” internodes that are more dynamic in response to sensory experience and remodel readily in response to abnormal experience (Micheva et al., 2018; Yang et al., 2020). PV+ interneurons are also associated with elaborate extracellular matrix structures, perineuronal nets, that act as a structural brake to plasticity by restricting the lateral mobility of receptors and anchoring synaptic connections (Levelt and Hübener, 2012; Sorg et al., 2016).

Cell morphology has been the classic approach to reconciling cellular diversity in the brain. From Cajal’s foundational work in the nineteenth century, a range of extensive shapes have been described in elaborate but qualitative terms. (Cajal, 1995). This has persisted in classic anatomical investigations of PV+ cells, where their morphologies were defined in terms of descriptions of their appearance together with their laminar locations in the cortex (Mikkonen et al., 1997; Zhu et al., 2018; Song et al., 2023). This gap in knowledge stems from a widening technical gap surrounding how to analyze single cells in anatomical techniques. Emerging approaches to quantifying and studying cell morphologies have identified subtle changes in cell shape that are imperceptible in conventional image analyses, carry salient, biologically relevant information about cell states in disease and quantitatively define disease progression at a single-cell level (Alizadeh et al., 2016; Hagemann et al., 2021). As single-cell transcriptomics has demonstrated specific transcriptomic types of PV+ interneurons are uniquely implicated in neurodegenerative disorders (Gabitto et al., 2024), it remains unknown whether there are specific PV+ interneuron morphologies that are indicative of an increased susceptibility to disease. To obtain these insights, comparable approaches to data acquisition and analysis are needed to align classic neuroanatomical data with single-cell transcriptomic approaches (Gliko et al., 2024).

In this study, we developed a sensitive, data-driven approach to define and characterize the cell morphologies of PV+ interneurons. We also applied an unsupervised, standardized approach to mapping the cortical layers. We robustly interrogate the morphology of the cell body and explore the patterns and relationships that may be present with dendritic structure using an integrated dataset of ISH-labelled and whole-filled cells. Finally, we explore the relationship between cell morphologies and transcriptomics by applying our pipeline to a collection of genes associated with PV+ transcriptomic types.

## Methods

### Animals

All anatomical data were retrieved from the Allen Brain Institute mouse ISH database (https://mouse.brain-map.org/) (Lein et al., 2007). The images in this database were from male C57BL6 mice at postnatal day 56 (P56). For the current study, PV+ data was collected from 6 mice. However, in the quality control step (described below), the images from one animal did not meet the criteria for inclusion, so the animal’s data were not included in subsequent analyses.

### Assessing Label Quality

For the dataset of PV+ cells, we assessed the label quality among the 6 animals. A single sample colorimetric ISH from S1 and V1 were processed and analyzed in FIJI. Images were first converted to grayscale, followed by a 50-pixel rolling ball background subtraction, before being binarized. In these processed images, the integrated density was obtained for all cells in the image. Estimation statistics were performed comparing the unpaired median differences for integrated density values among the six animals (using the dabestr package - version 0.3.0 in R) (Ho et al., 2019).

### Datasets

A summary of the composition of each dataset is shown in Table 1.

**Table 1.** Dataset composition summary.

| Dataset | Number of Animals | S1 cell count | V1 cell count | PV | Ndnf | Cux2 | Rorb | Deptor | Foxp2 | Ctgf | Cacna2d3 | Calb1 | Bdnf | Col25a1 | Etv1 | Fxyd6 | Kit | Krt12 | Nfib | Snca | Tac1 | Thsd7a | Tpbg |
| --- | --- | --- | --- | --- | --- | --- | --- | --- | --- | --- | --- | --- | --- | --- | --- | --- | --- | --- | --- | --- | --- | --- | --- |
| PV+ cells | 5 | 8823 | 5451 | ✓ |  |  |  |  |  |  |  |  |  |  |  |  |  |  |  |  |  |  |  |
| Laminar Marker Genes | 6 | 1383 | 645 |  | ✓ | ✓ | ✓ | ✓ | ✓ | ✓ |  |  |  |  |  |  |  |  |  |  |  |  |  |
| Whole, Filled Cells | 3 | 0 | 36 | ✓ |  |  |  |  |  |  |  |  |  |  |  |  |  |  |  |  |  |  |  |
| PV+ Cells and Associate d Genes | 13 | 2980 | 2162 | ✓ |  |  |  |  |  |  | ✓ | ✓ | ✓ | ✓ | ✓ | ✓ | ✓ | ✓ | ✓ | ✓ | ✓ | ✓ | ✓ |

Four datasets were constructed.

1. A dataset of ISH-labeled PV+ cells in S1 and V1.
2. A dataset of ISH-labeled cells for 6 laminar marker genes in S1 and V1.
3. A dataset of whole-cell filled PV+ cells in V1.
4. A dataset of ISH-labeled cells for 13 genes associated with PV+ transcriptomic types.

For the first three datasets, the colorimetric ISH and processed expression images were retrieved at full resolution (1 pixel = 1 μm). The Allen Brain Institute preprocessed the expression images to remove the background (black) and colour-code the cells by their integrated optical density, reflecting the density of the ISH labelling (blue = minimum integrated density, red = maximum integrated).

The first dataset of PV+ cells used 11 images from sagittal sections containing S1 and V1 for each of the 5 animals. The reference atlas (https://atlas.brain-map.org/) was used to identify the middle of S1 & V1, and 5 images medial and lateral to that image were added for a total of 11 images. Each sagittal image contained both S1 and V1. Sampling boxes from the pial surface to the white matter (567μm X 1525μm) for the colorimetric ISH and expression images were extracted from S1 and V1 using Adobe Photoshop 2024. Thus, S1 and V1 had 11 ISH and expression image sampling regions extracted for analysis.

The second dataset used images from 6 laminar marker genes associated with a specific cortical layer (L1 - Ndnf, L2/3/4 - Cux2, L4 - Rorb, L5 - Deptor, L6A - Foxp2, L6B - Ctgf). One ISH and the corresponding expression image were taken from the middle of S1 and V1 for each gene, and a sampling box was extracted from each cortical area.

The third dataset used images from S1 and V1 for 13 genes (Bdnf, Calb1, Cacna2d3, Col25a1, Etv1, Fxyd6, Kit, Krt12, Nfib, Snca, Tac1, Thsd7a, Tpbg) associated with PV+ T-types (Tasic et al., 2016). Among the marker gene list for PV+ T-types in Tasic et al., 2016, these 13 genes had good quality labelling in the ISH database. Furthermore, these genes also represent the multimodal PV+ transcriptomic types defined in Gouwens et al., 2020. Again, one ISH and the corresponding expression image from the middle of S1 and V1 were used for each of the 13 genes.

The fourth dataset used 36 whole-cell filled PV+ cells from the Allen Cell Types Database (https://celltypes.brain-map.org/data) (Hodge et al., 2019). The cells used for analysis were from the 5 PV+ transgenic lines (Htr3a-Cre_NO152|Pvalb-T2A-Dre, Pvalb-IRES2-Cre-Neo, Pvalb-IRES-CRE, Pvalb-T2A-CreERT2, PVALB-T2A-FlPO|Vipr2-IRES2-CRE) and were categorized as aspiny neurons with complete reconstructions. Cells were sampled from V1 (VISp) in layers ⅔ to 6A (no cells were filled from layers 1 or 6B). Mice from both sexes aged P56 ± 3 days were in the database; however, the metadata did not specify the sex of individual mice (Hodge et al., 2019).

Since the whole-cell-filled dataset did not include expression images, an image of each soma had to be extracted for analysis. Images of the 36 cells were downloaded from the database and opened in Adobe Photoshop 2024. The magic wand tool (default tolerance parameters) was used to identify and copy the cell body into a “simulated cortex”. Two of the 36 cells had a dark halo around the cell body, so the levels tool was used to increase the contrast, and the soma was extracted with the magic wand. The simulated cortex contained 36 cells at their laminar location, which was identified in the Allen Cell Types Database.

### Image Processing and Analysis

A custom analysis pipeline was created in CellProfiler (Carpenter et al., 2006; Stirling et al., 2021) to quantify the morphology of the cells from the PV+, PV+ associated genes and whole-cell filled datasets. The first step converted the RGB expression images to grayscale using the ColorToGray Module, where the red, blue, and green channels were given equal weights of 1. The grayscale images were binarized using the Threshold module and the global threshold strategy because the expression images had uniform backgrounds. Otsu’s thresholding method and two-class thresholding were used to distinguish labelled cells and particles from the background. The thresholding was not smoothed and was not corrected. Next, the watershed algorithm was applied to segment the cells. The default watershed parameters were used (segmentation generated from a distance between adjacent objects, footprint of 8, and no downsampling). Finally, measurements of the cell morphologies were obtained using the MeasureObjectSizeShape module. This produced a data matrix where each row corresponds to a cell, and columns contain cell coordinates, metadata, and morphometric features. Also, an image of all the analyzed cells was generated. The unprocessed data file and CellProfiler-generated images were exported using the ExportToSpreadsheet and SaveImages modules.

### Cleaning and Extracting Cell Morphology Data

A two-step data-cleaning process was implemented in RStudio to prepare the unprocessed data file for analysis. First, the laminar location of all cells was converted into a standardized cortical thickness. Second, the data were filtered to remove particles that were either too small or irregularly shaped to be categorized as a neuron (Kooijmans et al., 2020).

#### Converting to Standard Cortical Thickness

In mice, cortical thickness is relatively uniform; however, for the subsequent analyses, it was important to ensure that the laminar location of the cells was in standardized units as the relative depth from the pial surface to the white matter. The distance from the pial surface to the start of the white matter was measured at ten uniformly spaced intervals across the width of the image, and the y coordinate for each cell was transformed such that 0 was the pial surface and 1 was the bottom of layer 6B. Any cells with y coordinates outside the thickness of the cortex, either above the pial surface or below layer 6B, were removed.

#### Removing Non-Cell Shaped Particles

The data file was filtered in RStudio to include cells within the known size (50-1500μm^2^) and circularity for neurons (0.2-1.0) (Kooijmans et al., 2020). This step removed irregularly shaped particles and small or large artifacts in the expression images. The final cell counts for each of our datasets are shown in Table 2.

**Table 2.** Cell Counts for Cell Morphology Datasets Prepared in this Study.

| **Dataset** | **S1 cell count** | **V1 Cell Count** |
| --- | --- | --- |
| **PV+ Cell Dataset (14274 cells)** |  |  |
| PV | 8823 | 5451 |
| **Laminar Marker Gene Dataset (2028 cells)** |  |  |
| L1 : Ndnf | 47 | 10 |
| L2/3/4 : Cux2 | 581 | 401 |
| L4 : Rorb | 515 | 71 |
| L5 : Deptor | 51 | 28 |
| L6A: Foxp2 | 82 | 95 |
| L6B: Ctgf | 107 | 40 |
| PV+ Genes Dataset (19416 cells) |  |  |
| PV | 8823 | 5451 |
| Bdnf | 210 | 177 |
| Cacna2d3 | 495 | 336 |
| Calb1 | 409 | 310 |
| Col25a1 | 547 | 264 |
| Etv1 | 157 | 155 |
| Fxyd6 | 326 | 258 |
| Kit | 48 | 50 |
| Krt12 | 62 | 62 |
| Nfib | 71 | 58 |
| Snca | 494 | 307 |
| Tac1 | 20 | 28 |
| Thsd7a | 55 | 64 |
| Tpbg | 86 | 93 |
| PV+ ISH and Whole-Cell Fill Dataset (5487 Cells) |  |  |
| PV+ ISH Cells (V1 Only) | - | 5451 |
| PV+ Filled Cells (V1 Only) | - | 36 |

### Identifying Cortical Layers

The dataset of laminar marker genes was used to define cortical layer boundaries in standard coordinates for S1 and V1. Laminar density profiles were calculated for each laminar marker gene in both S1 and V1 using the density() function in R. The density profiles for laminarly adjacent pairs of genes (e.g., L4 and L5) were used to calculate the intersections of the two profiles. A function for the difference between the two density profiles was obtained by linear interpolation using the approx() function before solving it to find the intersections between the two linear density profiles. The intersections were calculated using the uniroot.all() function from the rootSolve package in R (Soetaert and Herman, 2009) and serve as putative layer boundaries for S1 and V1.

### Cluster Analysis of Cell Morphologies

#### Preparing the Data Frames of Features and Cell Identifiers

We developed a sensitive, data-driven approach to analyze the cell morphology data and characterize morphology clusters. The first step was to separate the cell metadata (i.e., cortical area, animal ID, gene symbols) and X, and Y coordinates from the morphology measurements, creating 2 datafiles. Next, the redundant features (Central Moment 0_0, Spatial Moment 0_0, Inertia Tensor 0_1, Inertia Tensor 1_0) that duplicate cell area were removed. The remaining 97 size and shape morphometric features are listed in Table 2. The data frame used for the cluster analysis had a row for each cell and 97 columns of morphometric features.

Before clustering the morphometric data, a logarithmic transformation was applied to the seven Hu Moment Features (Hu Moments 0,1,2,3,4,5,6) to normalize the distribution. Then, all the data were z-scored to ensure that each feature had the potential to contribute equally to the cluster analysis.

#### Selecting the number of clusters

The elbow method was used to select the number of clusters by calculating the within-cluster sum of squares (WCSS) for a range of different numbers of clusters (k-values). The optimal k is identified as the point of maximal curvature, or “elbow”, on the WCSS graph. The elbow was calculated using three different R packages and functions: the fviz_nbclust() function from the factoextra package (Kassambara and Mundt, 2020), the find_curve_elbow() function from the pathviewr package (Baliga et al., 2023), and the elbow function from the elbow package (Casajus, 2020) and compared among the three results. Agreement among two or more elbow plot methods confirmed the optimal k-value for cluster analysis.

#### Robust-Sparse K-Means Clustering

We applied the Robust-Sparse K-Means Clustering (RSKC) algorithm to cluster the morphometric data. This adaptive, iterative clustering algorithm is robust to outliers and does not create clusters with one or comparatively few members (Kondo et al., 2016). In addition, it provides feature weights of the relative contributions of each morphological feature to the clustering. The greater the contribution of a feature to the clustering, the higher the weight assigned from RSKC. The top morphological features were defined as those carrying at least 50% of the maximum weight given to a single feature. We performed RSKC cluster analysis using the rskc function from the RSKC package in R.

#### Dimensionality Reduction and Visualization

To reduce the dimensionality and visualize the clusters in the high-dimensional cell morphology space, we applied the density-preserving t-SNE (denSNE) algorithm. This is an optimized implementation of the t-SNE algorithm (Maaten and Hinton, 2008) that better preserves the density of data points around patterns in local and global structures in the high-dimensional space (Narayan et al., 2021). The RSKC clustering was analyzed by multiplying the z-scored morphometric feature data by the corresponding feature weights (Balsor et al., 2021). Then, the cluster content was inspected by colour-coding the denSNE plots using the z-scored morphological values for the top-weighted features.

#### Characterizing Morphological Clusters

To quantify and characterize the representative morphological features in each cluster, the median z-scored values for the top-weighted morphology features were plotted in a heatmap to create a morphology phenotype matrix. The matrix had a column for each cluster and rows for the top-weighted morphology features. The resulting heatmap was organized by unsupervised hierarchical clustering (Ward.D2 method) to group similar clusters and features. The morphology phenotype matrix was then used to identify which size and shape morphological features drove the clustering.

The median z-scored values for the top-weighted size and shape features were used to identify morphological motifs. The number of size and shape morphology motifs were determined by applying unsupervised hierarchical clustering (Ward.D2) and the elbow method as described previously.

### Cluster Analysis of the PV+ Cell Dataset

The morphological analysis pipeline, as described above, was applied to our dataset of 14274 PV+ cells labelled by ISH. In this analysis, cells from V1 and S1 were analyzed together. After obtaining morphological clusters for PV+ cells, we assessed whether any morphology clusters were specific to either V1 or S1. We compared the proportions of cells from each cortical area for each morphology cluster to the global proportion of cells in this dataset (S1:V1 = 62%38%). We assessed these proportions using the chi-square test for goodness-of-fit.

### Laminar Analysis of PV+ Morphology Clusters

The laminar location of the cells in each PV+ cluster was visualized by mapping the location of each cell in a cluster using their standardized XY coordinates. This provided a visualization of the laminar location of all the cells that could be used to define the laminar profile for each morphological cluster using data from S1 and V1 for all animals. Then, laminar density profiles for each cluster in S1 and V1 were calculated using the ggMarginal() function from the ggExtra package in R (Attali and Baker, 2023). Laminar profiles were calculated separately for morphology clusters in S1 and V1.

#### Identifying Laminar Patterns

To determine if PV+ morphology clusters had different laminar arrangements, the 130 laminar density profiles (13 morphology clusters x 2 cortical areas x 5 animals) clustered. Since the laminar boundaries were at different depths in S1 and V1 we performed a cubic piecewise spline fit to transform the Y coordinate of V1 cells into the S1 cortical space. This allows us to compare the laminar profiles in V1 and S1, accounting for the difference in cortical thickness among the two cortical areas. To perform the interpolation, we first resampled the S1 cells, so there were the same number of cells from S1 and V1. Then, we performed the spline fit on the V1 cells. A single set of layer boundaries can be used by interpolating data from the two cortical areas into a common cortical space.

Unsupervised hierarchical clustering (Ward.D2 method) was used to identify laminar patterns. We started by constructing a dataframe of linear densities for each profile using the density() function in R. Next, we z-scored the linear density values so each profile had equal potential to impact the clustering. After determining the optimal number of clusters with the elbow method, we applied the Ward.D2 unsupervised hierarchical clustering to the z-scored linear density profiles for each morphology cluster from each cortical area for each animal. The average laminar profile for each cluster was calculated using all the laminar profiles within a cluster and visualizing those with a 95% confidence interval. We then assessed the area bias of each of these laminar patterns by comparing the proportion of laminar profiles from each pattern from each cortical area to the expected proportions (S1:V1 = 50%:50%). We assessed these proportions using the chi-square test for goodness of fit. We calculated the Shannon Entropy for each laminar pattern to quantify the morphological diversity in each laminar pattern.

### Cluster Analysis of the Whole-Cell Filled Dataset

#### Clustering Cell Body Morphologies

As described above, the cluster analysis pipeline was applied to the whole-cell filled dataset by combining it with the PV+ data from V1. Morphological data were prepared as described previously, and clustering was performed on morphometric z-scores. We determined the optimal number of clusters using the elbow method, partitioned cells into clusters using the RSKC algorithm, and visualized the clustering using the denSNE algorithm. For this dataset, implementing the denSNE serves the additional purpose of identifying whether whole-cell filled cells integrate into the clusters formed with V1 PV+ cells. We constructed morphology phenotypes for each cluster, paying particular attention to the clusters that contained whole-cell filled cells.

#### Clustering Dendritic Morphologies

In addition to the 97 cell body morphology measurements for the 36 whole-cell filled cells, 20 dendritic morphology features were available to quantify the dendritic arbour morphologies. The features are listed in Table 4. We determined the optimal k using the elbow methods before applying the RSKC algorithm to z-scored measures of dendritic morphology and obtained weights for the 20 features. As there are fewer whole-cell filled cells, we used the .swc neuronal reconstructions for these neurons to visualize each cell in a cluster. Reconstructions were plotted using the Allen Institute’s Cell Types Explorer Jupyter Notebook (unmodified original available from : https://alleninstitute.github.io/AllenSDK/cell_types.html), which we modified to automate and pseudo-color the reconstruction plots for improved visualization. We then visualized the correspondence between dendritic and cell body morphology clusters using the ggalluvial() package in R (Brunson, 2023).

**Table 3.** 97 Morphometric Features for Cluster Analysis.

| Morphometrics included in cluster analysis |  |  |
| --- | --- | --- |
| MaximumRadius | SpatialMoment_1_2 | SpatialMoment_2_3 |
| MeanRadius | SpatialMoment_2_1 | Zernike_3_3 |
| MinorAxisLength | HuMoment_2 | Zernike_7_5 |
| EquivalentDiameter | Zernike_5_1 | Zernike_5_5 |
| InertiaTensorEigenvalues_1 | HuMoment_0 | NormalizedMoment_1_1 |
| MinFeretDiameter | Zernike_1_1 | Zernike_5_3 |
| Area | Zernike_8_4 | NormalizedMoment_2_0 |
| MedianRadius | Zernike_3_1 | Zernike_7_3 |
| ConvexArea | SpatialMoment_0_3 | CentralMoment_1_3 |
| Perimeter | Zernike_8_6 | Zernike_7_1 |
| MaxFeretDiameter | InertiaTensor_1_0 | NormalizedMoment_3_1 |
| Eccentricity | CentralMoment_2_2 | AspectRatio |
| MajorAxisLength | Zernike_8_0 | Orientation |
| BoundingBoxArea | CentralMoment_1_1 | NormalizedMoment_2_2 |
| Zernike_0_0 | Compactness | HuMoment_4 |
| HuMoment_1 | Zernike_8_8 | NormalizedMoment_1_3 |
| SpatialMoment_0_1 | SpatialMoment_2_2 | HuMoment_5 |
| SpatialMoment_1_0 | Zernike_6_0 | NormalizedMoment_0_2 |
| InertiaTensorEigenvalues_0 | Solidity | NormalizedMoment_3_3 |
| HuMoment_3 | Extent | CentralMoment_0_3 |
| InertiaTensor_1_1 | SpatialMoment_1_3 | CentralMoment_2_1 |
| Zernike_4_0 | Zernike_4_4 | NormalizedMoment_2_1 |
| Zernike_2_2 | Zernike_9_7 | CentralMoment_0_1 |
| SpatialMoment_1_1 | Zernike_6_6 | NormalizedMoment_1_2 |
| CentralMoment_2_0 | Zernike_9_9 | CentralMoment_1_2 |
| InertiaTensor_0_0 | Zernike_9_5 | HuMoment_6 |
| CentralMoment_0_2 | Zernike_4_2 | NormalizedMoment_0_3 |
| SpatialMoment_2_0 | Zernike_6_4 | NormalizedMoment_3_0 |
| Zernike_2_0 | Zernike_7_7 | CentralMoment_2_3 |
| SpatialMoment_0_2 | Zernike_6_2 | CentralMoment_1_0 |
| FormFactor | Zernike_9_3 | NormalizedMoment_2_3 |
| Zernike_8_2 | Zernike_9_1 | NormalizedMoment_3_2 |
|  |  | EulerNumber |

**Table 4.** Dendritic Morphology Features for Whole-Cell Filled Cells.

| Morphometric |  |
| --- | --- |
| number_nodes | overall_height |
| total_length | average_fragmentation |
| average_contraction | average_parent_daughter_ratio |
| average_diameter | overall_depth |
| total_surface | number_stems |
| number_tips | max_euclidean_distance |
| number_branches | max_path_distance |
| number_bifurcations | soma_surface |
| total_volume | average_bifurcation_angle_local |
| overall_width | max_branch_order |

### Cluster Analysis of PV+ Associated Genes Dataset

The cluster analysis pipeline was applied to the dataset of 13 genes associated with PV+ T-types. In this analysis, we treated S1 and V1 separately.

For each cortical area, RSKC was used to identify morphological clusters, denSNE was used to visualize the clustering and morphometric organization of the cell morphology space and morphology phenotypes for each cluster. We assessed the proportions of genes present in each morphology cluster to interrogate the relationship between cell morphologies and single-cell transcriptomics. We compared the within-cluster proportions to the global proportion of genes across the datasets for S1 and V1. We used the chi-square test for goodness of fit to determine if any clusters over- or under-represented the PV+ associated genes.

To explore the laminar distributions of PV+ associated genes, we calculated the laminar profiles of individual clusters for each gene. We calculated the profiles for clusters that had at least 10 cells from a gene and z-scored them. This revealed the morphological heterogeneity among groups of cells labelled for a single gene.

### Statistical Analyses

Estimation statistics were used to compare morphological differences among clusters (dabestr package version 0.3.0) (Ho et al., 2019).

## Results

### Assessing ISH Label Quality Among Animals

The first step was to assess the quality of the ISH images among the animals to ensure comparable labelling so differences in label intensity did not impact the analyses. This quality control step was done by visual inspection and quantitative comparison of the distribution of integrated densities for the PV+ ISH-labelled cells from each animal.

Visually, the ISH labelling of animal 2293 in both S1 and V1 appeared lighter than the other five animals (Fig. 2A i-ii). Next, estimation statistics were used to quantitatively compare the integrated density of labelled cells among the animals (Fig. 2B). That analysis showed that the labelling intensity of cells from animal 2293 was less than the other animals and that the distributions from the other animals overlapped. Because the labelling intensity of animal 2293 was different, it was excluded from the subsequent analyses.

**Figure 2.**
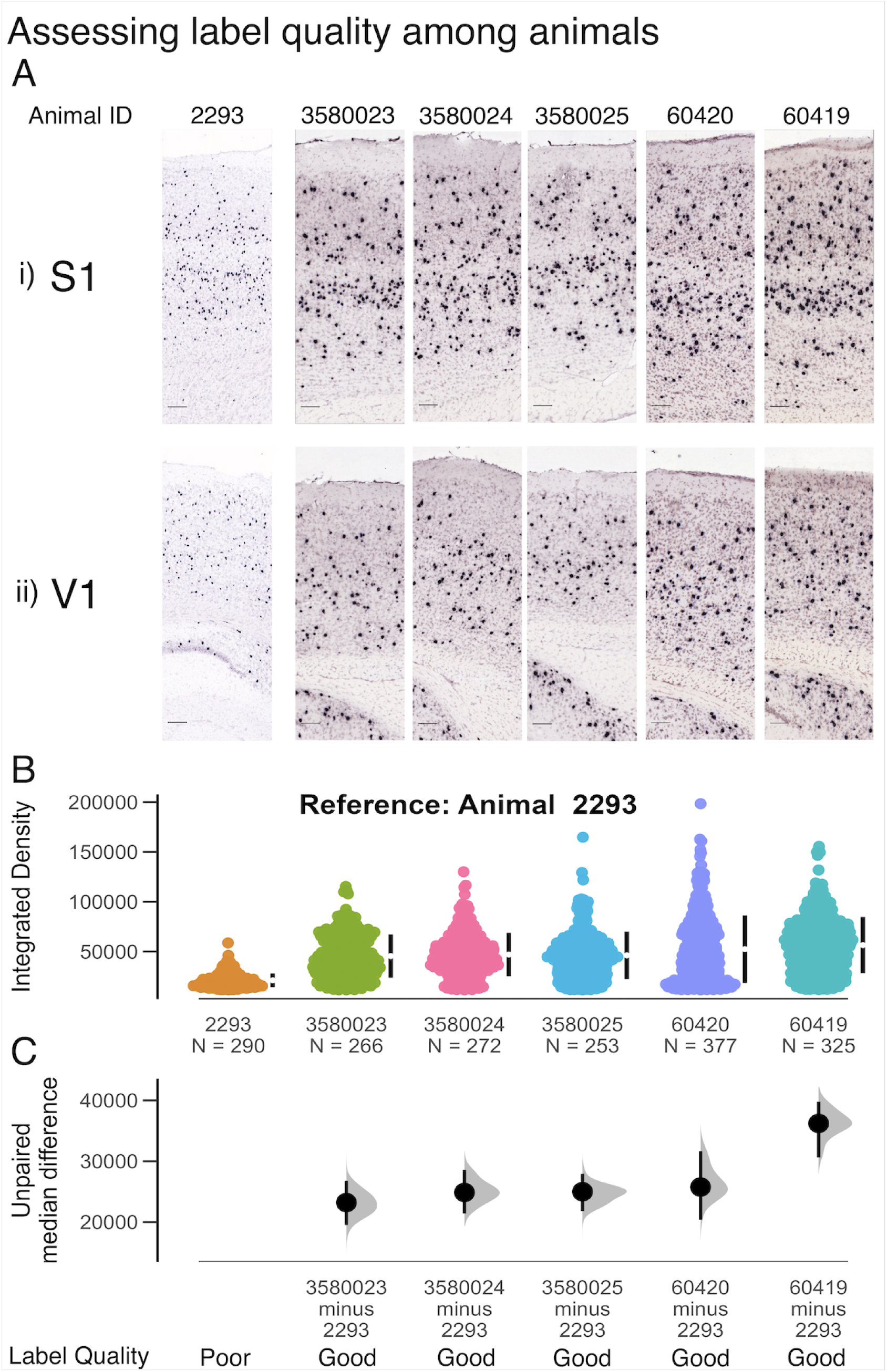
Data quality assessment among animals. (A) Samples of In Situ Hybridization taken from (i) S1, and (ii) V1 labelling the PV gene. In all images, scale bar = 100μm (B) Distributions of integrated density measurements (product of cell area and mean gray value) for all cells in each sample from each animal. Single points are single cells, and the distribution is constructed from both S1 and V1. The median integrated density value is shown as a white dot beside each bee swarm, and the 95% confidence interval is shown as the black vertical bar beside each bee swarm. (C) Distributions of unpaired median difference for integrated density values among animals, using data from animal 2293 as a reference. The unpaired median difference is shown as a black dot, and the 95% confidence interval is shown as the vertical bar.

### Defining cortical layers

To provide a high throughput analysis of PV+ somata in an anatomical context, we needed to define and map cortical layers an in a standardized and reproducible way. We used the laminarly restricted patterns for a panel of 6 laminar marker genes (L1 - Ndnf, L2/3/4 - Cux2, L4 - Rorb, L5 - Deptor, L6A - Foxp2, L6B - Ctgf) to construct a quantitative pipeline that would identify cortical layer boundaries in both V1 and S1.

We used the XY coordinates for each labelled cell in the ISH images shown in Figure 3 to identify layer boundaries. The coordinates were standardized among samples into a standard coordinate system, where the cortical depth ranged from 0 (pial surface) to 1 (bottom of L6B). We also used this standard coordinate system to identify subcortical cells (with a standard depth greater than 1) and excluded them from subsequent analyses.

**Figure 3.**
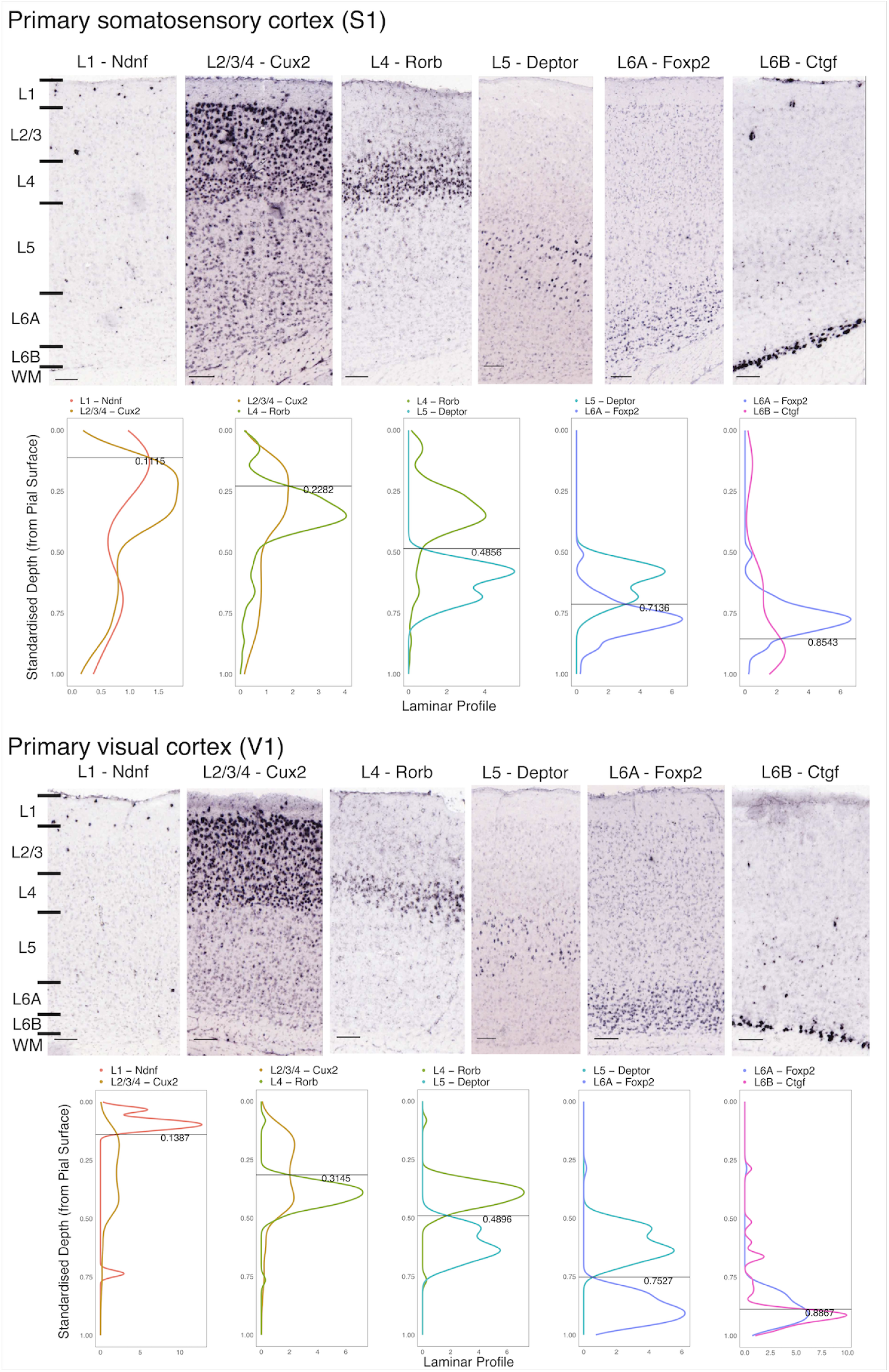
Defining cortical layer boundaries in S1 and V1. (A) upper panel : ISH for laminar marker genes extracted from S1, lower panel: laminar profiles for pairs of laminar marker genes that label adjacent layers. Laminar profiles are obtained in standard coordinates, where the depth ranges from 0 (pial surface) to 1 (bottom of L6B). The calculated intersections of the curves are taken as putative layer boundaries. (B): laminar boundaries calculated for V1. Upper panel : ISH for laminar marker genes. In all images, scale bar = 100μm. Lower panel: calculated layer boundaries from laminar profiles in standard coordinates.

The laminar profiles for each of the selected laminar marker genes were used to determine the layer boundaries in the standard cortical space. As expected, each laminar marker gene had a peak cell density in their respective layers. We determined the depth for layer boundaries in S1 (Fig.3A) and V1 (Fig. 3B) using the intersections of the laminar profiles for markers labelling pairs of adjacent cortical layers. These steps provided an unsupervised approach to identify the laminar boundaries in all animals.

### Laminar Cell Size Analysis of PV+ Cells

We began by applying a common approach to quantifying labelled cells using cell size and assigning cells to cortical layers. The goal was to test if that approach identified laminar differences in PV+ cells. Globally, between S1 and V1, the distributions of cell sizes overlap as shown in Figure 4A (S1: Median = 203μm^2^, Mean = 200.97 μm^2^, 95% CI = [199.06μm^2^-202.91μm^2^], V1: Median =184 μm^2^, Mean =184.39μm^2^, 95% CI = [182.18μm^2^ - 186.59μm^2^]). PV+ cell sizes overlapped in layers ⅔ to 6A. PV+ cells from Layer 1 in V1 (Median = 90 μm^2^, Mean = 114.27μm^2^, 95% CI = [102.83μm^2^ - 125.71μm^2^]) and S1 Layer 1 were smaller (Median = 149 μm^2^, Mean = 164.51 μm^2^, 95% CI = [154.55μm^2^ - 174.47μm^2^]) compared to cells from other cortical layers. Cells in Layer 6B are smaller than cells from layers ⅔-6A, but unlike Layer 1, the distributions overlap when comparing S1 and V1 (Fig. 4B). The distributions of cell sizes for S1 and V1 are consistently overlapping for L2/3-L6A (Fig. 5) The summary statistics for PV+ cell size by layer for each cortical area are in Table 5. Following this typical approach to analyzing PV+ cells in the cortex, we found that PV+ cell sizes overlap across Layers ⅔-6A, suggesting similar morphological organizations in those layers. However, this analysis only used cell size, whereas CellProfiler measured 97 size and shape parameters.

**Figure 4.**
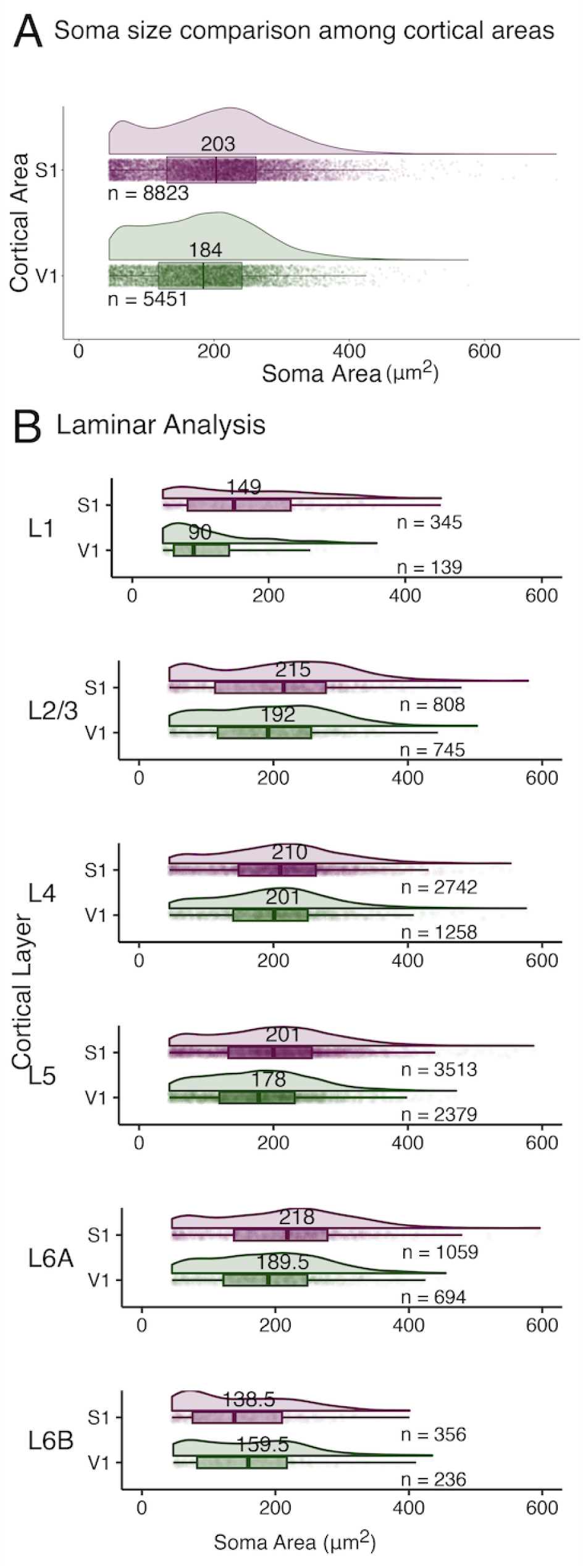
Size Distributions for PV+ interneurons in S1 and V1. (A) Raincloud plot comparing the global distributions of soma sizes of PV+ cells among S1 (purple) and V1(green). The global distribution is shown for each cortical area, above a horizontal boxplot. The median is shown above the boxplot at the median size. Cell size values are also plotted beneath the distribution curve, with single points as single cells. (S1: Median = 203μm^2^, Mean = 200.97 μm^2^, 95% CI = [199.06μm^2^-202.91μm^2^], V1: Median =184 μm^2^, Mean =184.39μm^2^, 95% CI = [182.18μm^2^ - 186.59μm^2^]). (B) Raincloud plots comparing the distributions of PV+ soma size among cortical areas for each cortical layer separately. Summary statistics comparing between cortical areas for each cortical layer are shown in Table 5.

**Figure 5.**
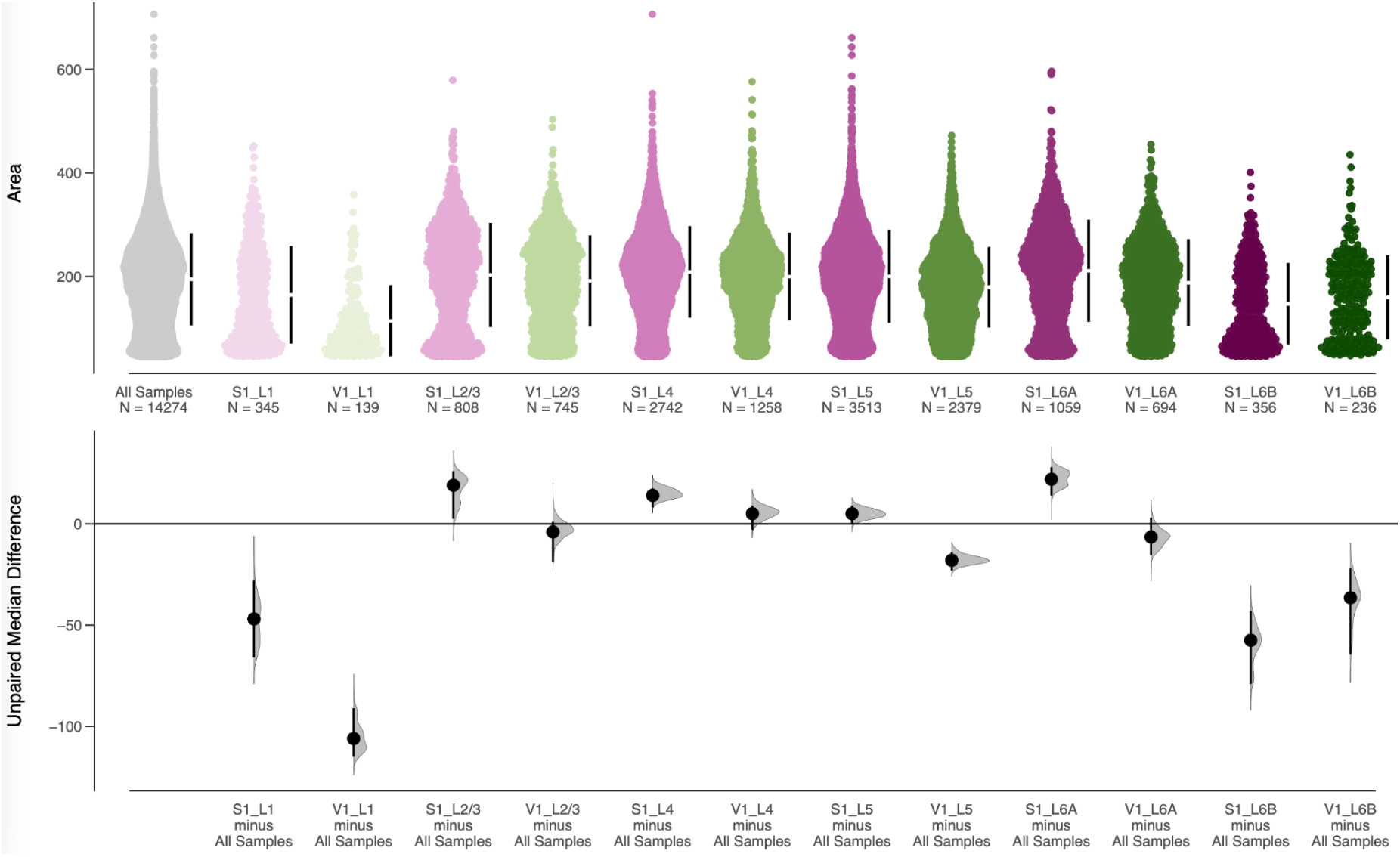
Estimation statistics comparing variation in PV+ soma size among cortical layers and areas. The unpaired median difference for 12 comparisons against a shared control are shown in the Cumming estimation plot. The reference distribution in this plot is all the data - representing the global distribution of cell size for the entire dataset. Cell size values are plotted on the upper axes as bee swarm plots. Purple bee swarms are from S1 and green swarms are from V1. In the bee swarm plots, single points are single cells. Beside each bee swarm, the median is shown as a white dot and the 95% confidence interval is plotted as a vertical black line. On the lower axes, the unpaired median differences are plotted as bootstrapped sampling distributions. The unpaired median difference is shown as a black dot and the 95% confidence interval is indicated by the vertical black line.

**Table 5.** Summary Statistics for PV+ Cell Size by Cortical Layer for S1 and V1.

| Cortical Area | Cortical Layer | Number of Cells | Median Cell Area ($\mu\text{m}^2$) | Mean Cell Area ($\mu\text{m}^2$) | 95% CI (Lower Bound) | 95% CI (Upper Bound) |
| --- | --- | --- | --- | --- | --- | --- |
| S1 | L1 | 345 | 149 | 164.51 | 154.55 | 174.46 |
| V1 | L1 | 139 | 90 | 114.27 | 102.83 | 125.71 |
| S1 | L2/3 | 808 | 215 | 203.12 | 196.19 | 210.04 |
| V1 | L2/3 | 745 | 192 | 191.63 | 185.32 | 197.94 |
| S1 | L4 | 2742 | 210 | 208.81 | 205.50 | 212.11 |
| V1 | L4 | 1258 | 201 | 199.94 | 195.25 | 204.64 |
| S1 | L5 | 3513 | 201 | 200.34 | 197.38 | 203.31 |
| V1 | L5 | 2379 | 178 | 179.31 | 176.18 | 182.43 |
| S1 | L6A | 1059 | 218 | 211.16 | 205.22 | 217.10 |
| V1 | L6A | 694 | 189.5 | 188.19 | 181.95 | 194.44 |
| S1 | L6B | 356 | 138.5 | 147.36 | 139.17 | 155.54 |
| V1 | L6B | 236 | 159.5 | 159.90 | 149.53 | 170.27 |

### Morphological Clustering of PV+ interneuron cell bodies

To determine if using a rich set of cell size and shape measurements would identify morphological differences among PV+ cells, we constructed a large data matrix of 97 morphology features (columns) for each of the 14274 PV+ cells (rows). This produces high-dimensional cell morphology data for each PV+ cell in our dataset. The optimal number of clusters (k = 13) was found using the elbow method (Fig. 6A), and the data were clustered using the Robust, Sparse K-Means Clustering (RSKC) algorithm. The RSKC algorithm was selected because it is an iterative clustering algorithm that is robust to outliers. Additionally, RSKC provides feature weights which inform us of the relative contribution of each morphology feature to the overall clustering (Kondo et al., 2016)).

**Figure 6.**
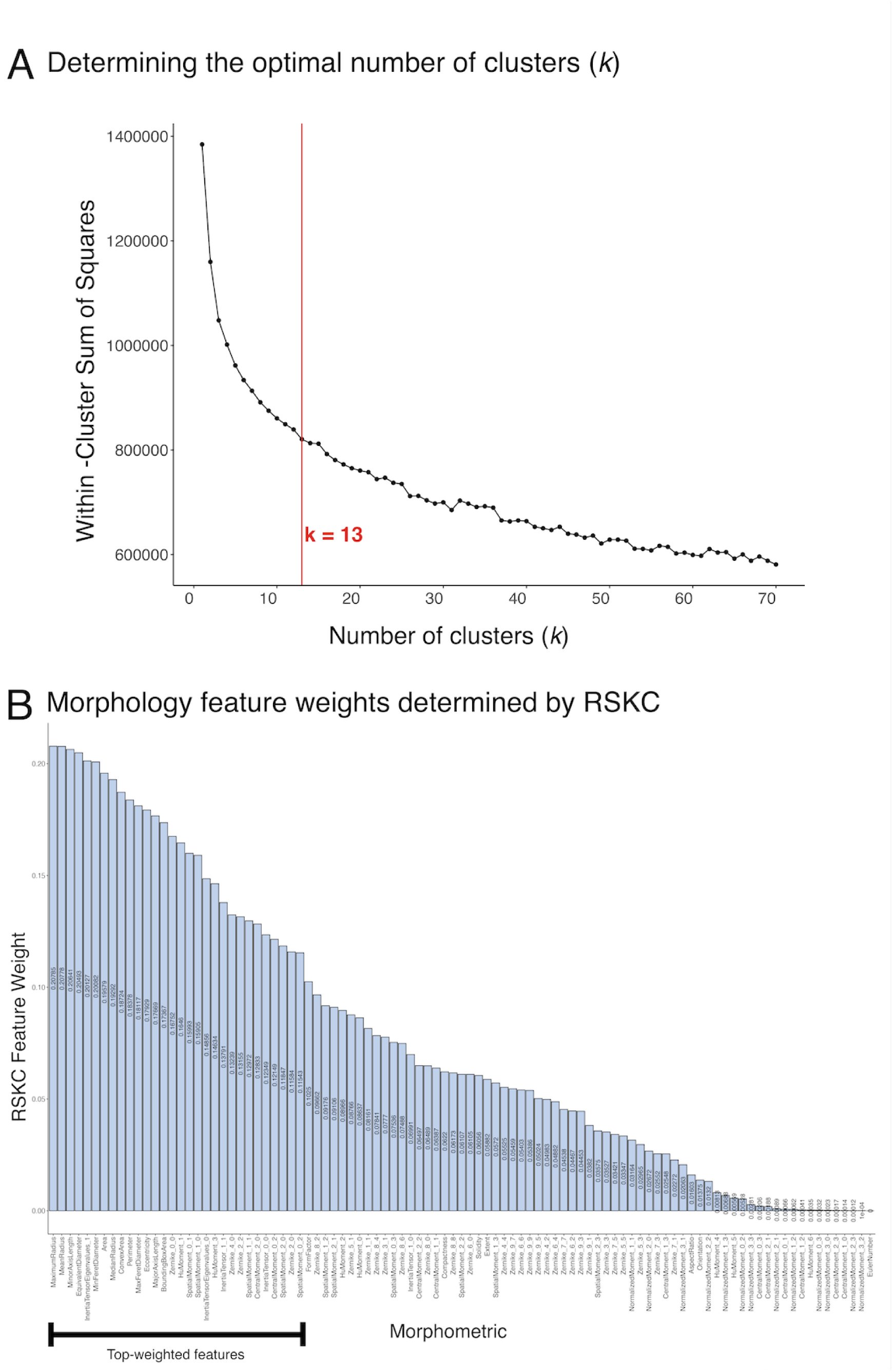
Unsupervised clustering on PV+ Interneuron morphologies. (A) Plot of the within-cluster sum of squares against the number of clusters, k. The optimal k-value (k = 13) was determined by the elbow method and shown by the red line. (B) Weights for all 97 morphology features as determined by the robust, sparse K-means clustering (RSKC) algorithm. Feature weights are plotted in decreasing order from left to right. Top-weighted features (>50% of maximum weight attributed to a single morphology feature), are identified by the black bar.

The feature weights from RSKC for the 13 morphology clusters of PV+ cells are shown in Figure 6B. Top-weighted features (>50% max feature weight) include a combination of size and shape parameters. Top-weighted size parameters include cell area, perimeter, diameters, axis lengths and radii. Among top-weighted shape parameters, both size-varying parameters that are proportional to the size of the cell (e.g., Spatial and Central Moments- which would also carry size information) and size-invariant parameters (e.g., Eccentricity, Zernike moments 0_0, 2_0, 2_2, ad 4_0 and Hu Moments 1 and 3) that describe the circularity, elongation and symmetry of the cell body. Morphology features carrying low weight include higher order Zernike, spatial and central moments that quantify more subtle differences in shape related to the symmetry of processes arranged around the cell body, and the Euler number, which describes the topology of the cell (i.e., the presence of holes or distensions of the cell body). Collectively, RSKC identified the morphology parameters that represent the largest variance among the cells. The distribution of RSKC weights describes morphology differences among cells in terms of their size, circularity, elongation, and symmetry of the cell body. Taken together, the feature weights from RSKC indicate that despite a consistent topological structure, there are aspects of cell size and shape that are sufficiently different to drive the separation of cells into distinct morphology clusters.

### Visualizing PV+ Morphology Clusters

To visualize the 13 morphology clusters, we applied the density-preserving t-SNE (denSNE) algorithm. For this analysis, the denSNE perplexity was set to 10, and the random number seed was 13579. The z-scored value for each morphometric feature was multiplied by the feature weight obtained from RSKC, and the denSNE algorithm was applied to these transformed values. This allows each feature to contribute to the dimensionality reduction in a manner consistent with their contribution to the clustering (Balsor et al. 2021). The denSNE plot was colour-coded for the 13 PV+ cell clusters and showed a mosaic of distinct morphological clusters (Fig.7A).

**Figure 7.**
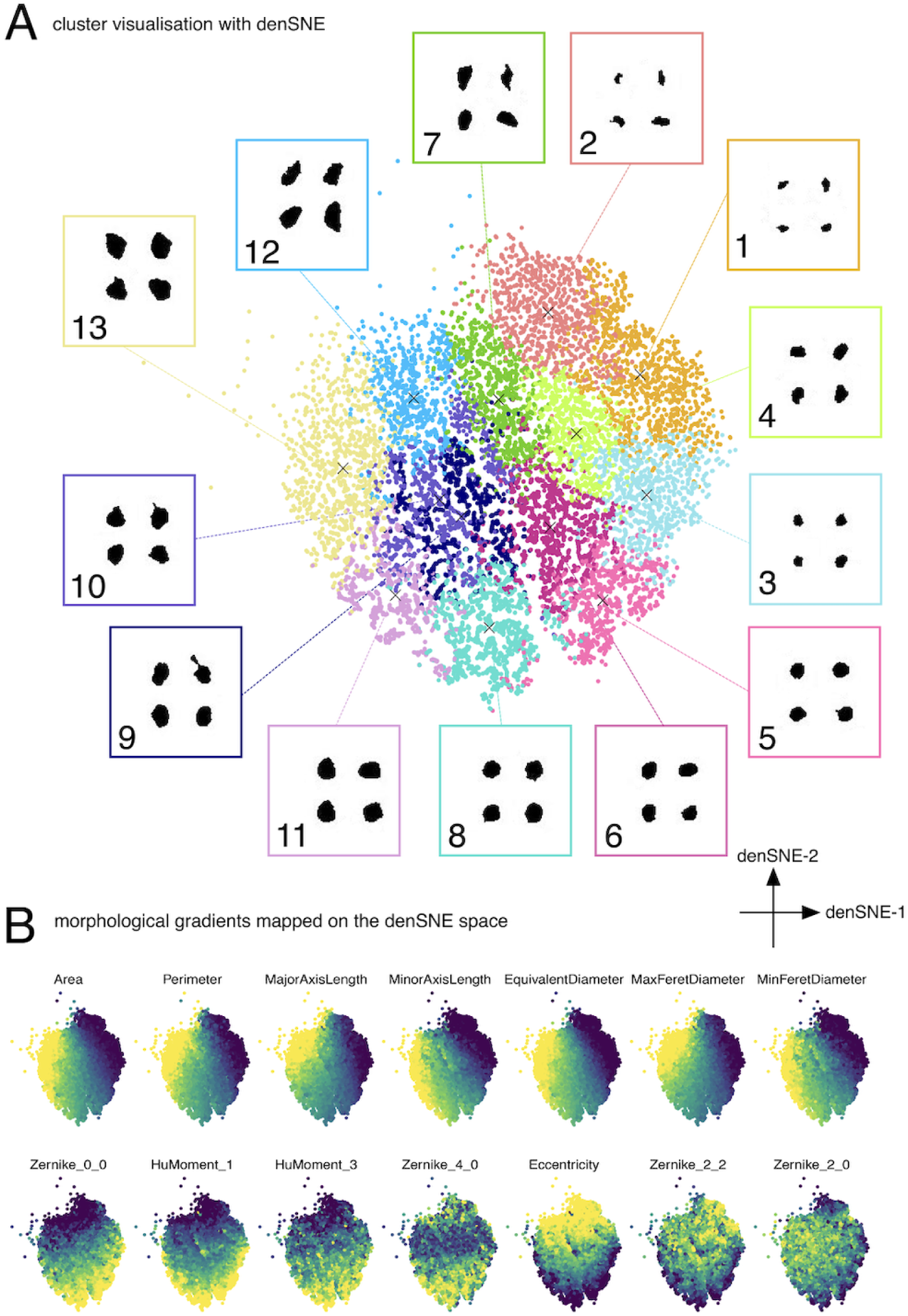
Visualizing RSKC morphology clusters for PV+ Cells: (A) Morphology space for PV+ cells from S1 and V1 visualized with density-preserving t-SNE (denSNE) (perplexity = 10, seed = 13579). The denSNE algorithm was applied to a transformed dataset where feature weights from RSKC were multiplied to the z-scored morphology feature measurements. Single points are single cells and the denSNE plot is coloured by cluster labels, with cluster centroids marked with a cross. Cluster numbers reflect the progression of median cell sizes (cluster 1 = lowest median size, and Cluster 13 = highest median cell size). 4 cells were sampled randomly from each morphology cluster and are shown in the inset boxes surrounding the denSNE plot. (B) Gradients of selected top-weighted morphology features mapped onto morphology space, visualized using the denSNE algorithm.The colour progression from navy to yellow indicates increasing feature values for each morphology feature. Size features are shown in the top row and shape features are shown in the bottom row.

We interrogated the distribution of features in the denSNE plot by colour-coding the points using the 14 top-weighted size and shape features (Fig.7B). These visualizations showed organized, gradual shifts across the desSNE. For example, the Area of the cells shifted from right to left going from small (dark blue) to progressively larger (yellow) cells. Similar gradient patterns were found for the other morphological features; however, the orientation of the gradient was different for the shape features.

The cell shape gradients in the denSNE varied from top to bottom. They were roughly orthogonal to gradients for the size features (Fig.7B). Importantly, the intersection of the size and shape gradients helped to identify morphological differences among the clusters. For example, clusters 1, 2 and 3 were small cells on the right side of the denSNE but were identified as different shapes and spread out across the vertical dimension of the denSNE plot.

We mapped the gradients of shape information and found measures of circularity increase from top to bottom, but measures of elongation increase from bottom to top in the denSNE plot (Fig.7B bottom row). Taken together, morphology clusters reflected the combination of size and shape features, which progressed orthogonally across the cell morphology space. Furthermore, the intersection of these gradients would determine how the morphology space is partitioned into clusters.

### Characterizing PV+ Cluster Morphologies

The denSNE plots (Fig. 7) indicate that the size and shape features grouped the cells into morphological clusters. However, a quantitative description of each cluster’s morphology features is needed to describe the characteristic PV+ soma phenotype for each cluster. We used the median z-scored of the top-weighted features for each of the 13 clusters to create a cluster-by-feature heatmap (Fig.8). The order of the morphology clusters (columns) and features (rows) was determined by hierarchical clustering. The size features ordered the 13 morphology clusters from left to right, with the clusters containing the smallest cells on the left and the largest on the right. The shape features further ordered the morphology clusters into interdigitated columns where clusters of similar-sized cells were separated based on the shape features. For example, morphology clusters 3 and 4 had a similar pattern of blue and green colour-coded cells for the size features but a different pattern for the shape features.

**Figure 8.**
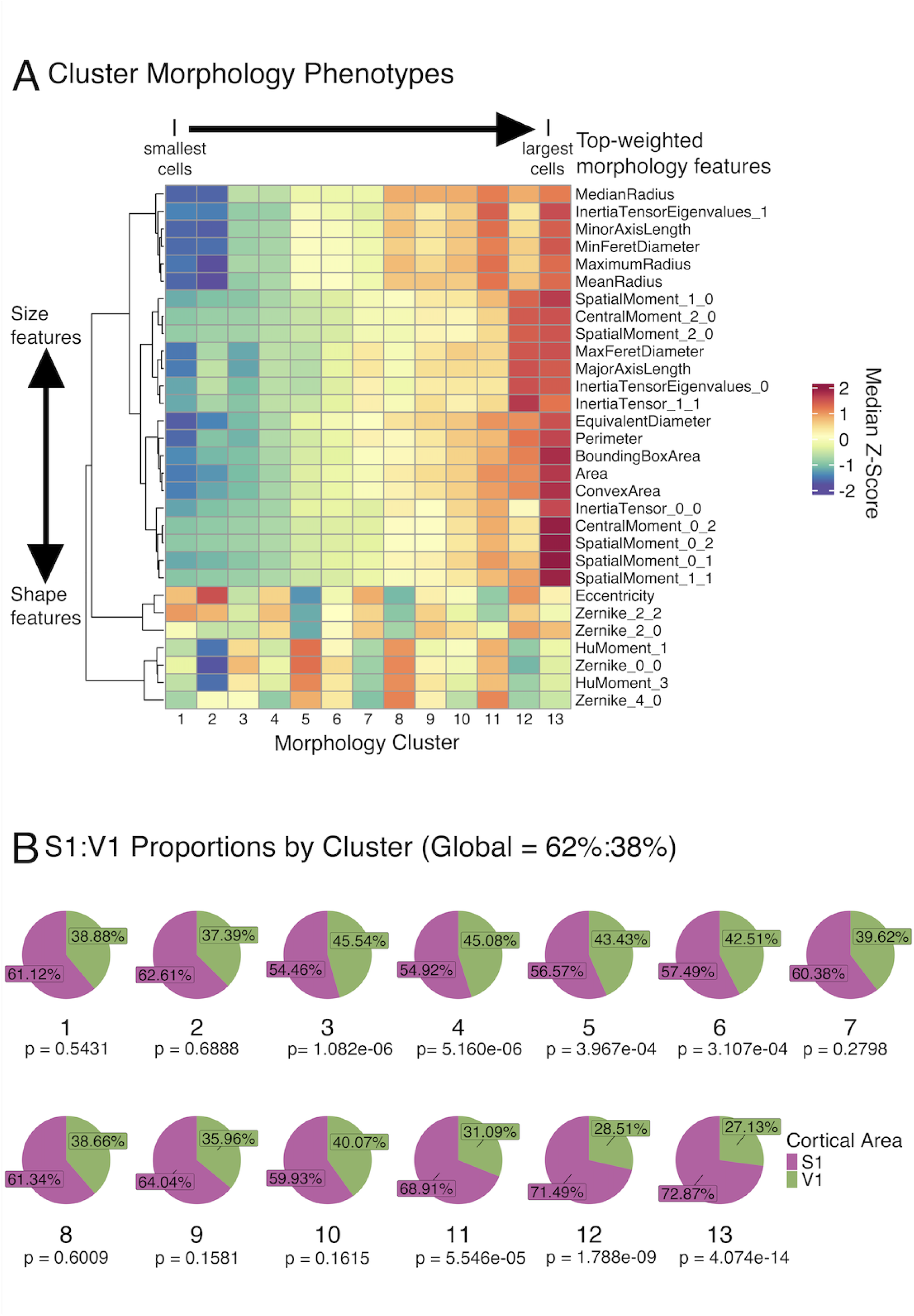
Characterizing RSKC morphology clusters. (A) cluster morphology phenotypes describing each cluster’s median size and shape characteristics. Phenotypes were constructed using the top-weighted morphology features, as shown in Figure 5B. Clusters are ordered and named according to the progression of median sizes (Cluster 1 = lowest median size, Cluster 13 = highest median size). The heatmap rows were organized using a row-wise dendrogram, using the Ward.D2 algorithm for hierarchical clustering. The first branch point on the row-wise dendrogram indicates the separation of top-weighted morphology features into size (towards the top of the heatmap) or shape features (towards the bottom of the heatmap). (B) Proportions of cells from each cortical area (S1 = purple, V1 = green) for each morphology cluster. P-values beneath each pie chart are used for the chi-square test, which compares the areal proportions within each cluster to the global areal proportion in the dataset. The overall proportion of cells from each cortical area in this dataset is S1 = 62% and V1 = 38%

We applied hierarchical clustering to the size and shape features separately within the phenotypes to determine how many different size divisions and shape motifs are represented in the 13 morphology clusters. The dendrograms for the size and shape clusters are shown in Figure 9. The clusters are separated coarsely into four major size divisions that we annotate as small, medium, large, and extra large. The clusters also separate into three prominent shape motifs that reflect fusiform, pyriform, and round cell shapes. We present here the characteristics of each shape motif.

**Figure 9.**
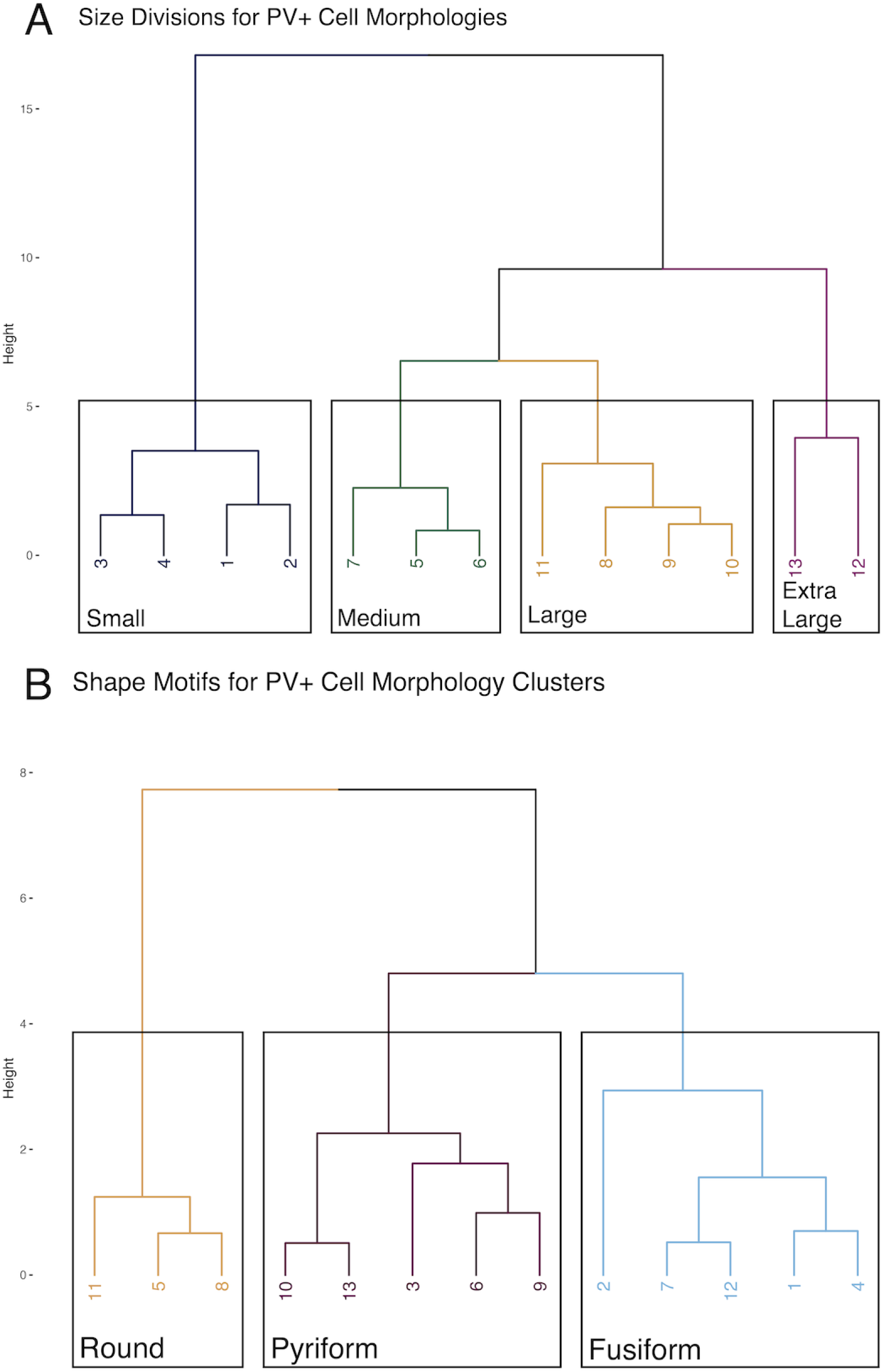
Characterizing size divisions and shape motifs among clusters. The median z-scores for top-weighted morphology features for each cluster were analyzed by unsupervised hierarchical clustering (Ward.D2) method, to identify the groups of clusters that share common size and shape characteristics.(A) Dendrogram for hierarchical clustering grouping individual morphology clusters into size divisions, based on the median values of top-weighted size features. (B) Dendrogram for hierarchical clustering grouping morphology clusters into shape motifs using the median z-scored values of top-weighted shape features. Cells fall into one of three motifs : fusiform, pyriform or round. The number of size divisions and shape motifs were determined using the elbow method.

#### Shape Motif 1 - fusiform cell bodies: clusters 1, 2, 4, 7, 12

These cells are elongated, as evidenced by their higher median eccentricity and second-order Zernike Moments. With increasing cell size, cells move from being more obliquely elongated to being vertically elongated-indicated by the progressive increase in median Zernike_2_0 (vertical astigmatism) and decrease in median Zernike_2_2 (oblique astigmatism). Cluster 2 was defined by a more extreme version of this shape motif - with more prominent elongation, resulting in sharply lower median measures of circularity (Fig.7A). These clusters occupy adjacent regions of the denSNE visualization of the cell morphology space (Fig.7). These clusters occupy a region of the morphology space defined by extreme shape characteristics, but still are organized along the size gradient. These clusters had higher median measures of elongation relative to clusters in other shape motifs (Figs. 11-13)

**Figure 10.**
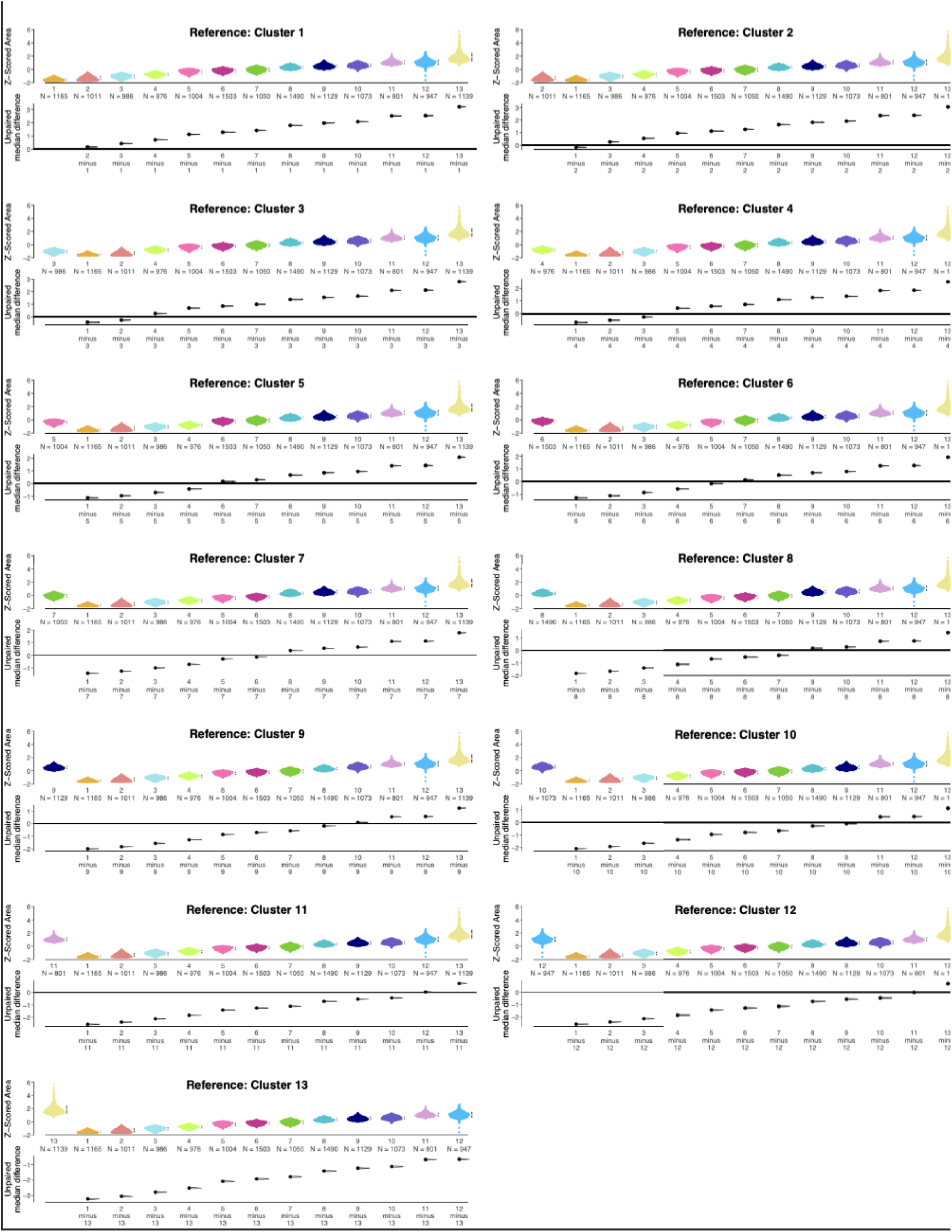
Soma Area for Each Morphology Cluster. Estimation statistics plots showing the distributions of z-scored cell body area for PV+ interneurons grouped by morphology cluster. The collection of plots shows comparisons among the clusters using each morphology cluster as a reference. Beeswarm plots of the z-scored cell body area are plotted on the upper axes of each plot. Beside each beeswarm the median z-scored cell size for each cluster is indicated as a white dot, and the 95% confidence interval about the median is shown by the black bar. The lower axes of each plot show the unpaired median difference plotted as a bootstrapped sampling distribution. The 95% confidence interval is shown as a black bar and the median is shown with a black dot.

**Figure 11.**
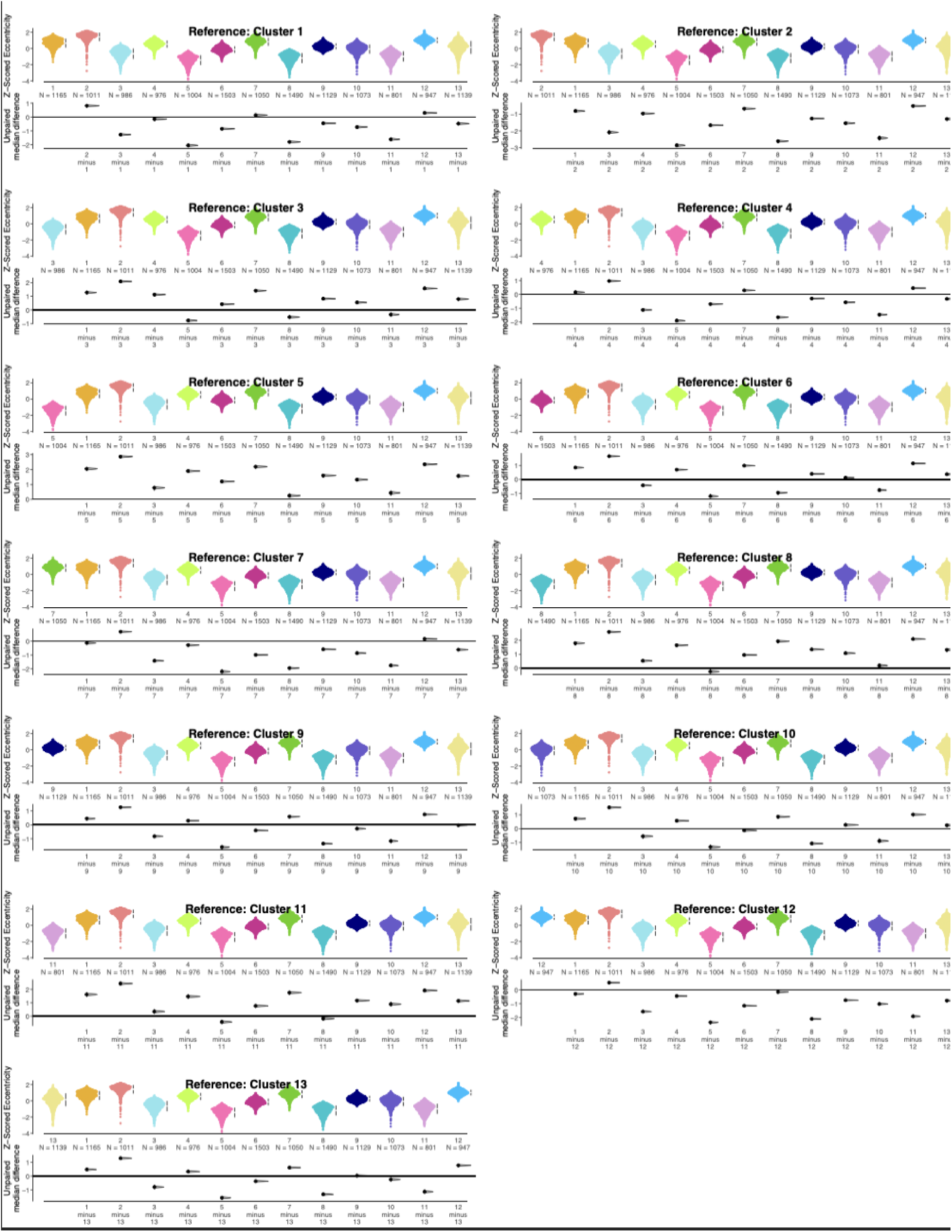
Cell eccentricity among morphology clusters of PV+ interneurons. Estimation statistics plots showing the distributions of z-scored cell body eccentricity for PV+ interneurons grouped by morphology cluster. The collection of plots shows comparisons among the clusters using each morphology cluster as a reference. Beeswarm plots of the z-scored cell eccentricity are plotted on the upper axes of each plot. Beside each beeswarm the median z-scored eccentricity for each cluster is indicated as a white dot, and the 95% confidence interval about the median is shown by the black bar. The lower axes of each plot show the unpaired median difference plotted as a bootstrapped sampling distribution. The 95% confidence interval is shown as a black bar and the median is shown with a black dot.

**Figure 12.**
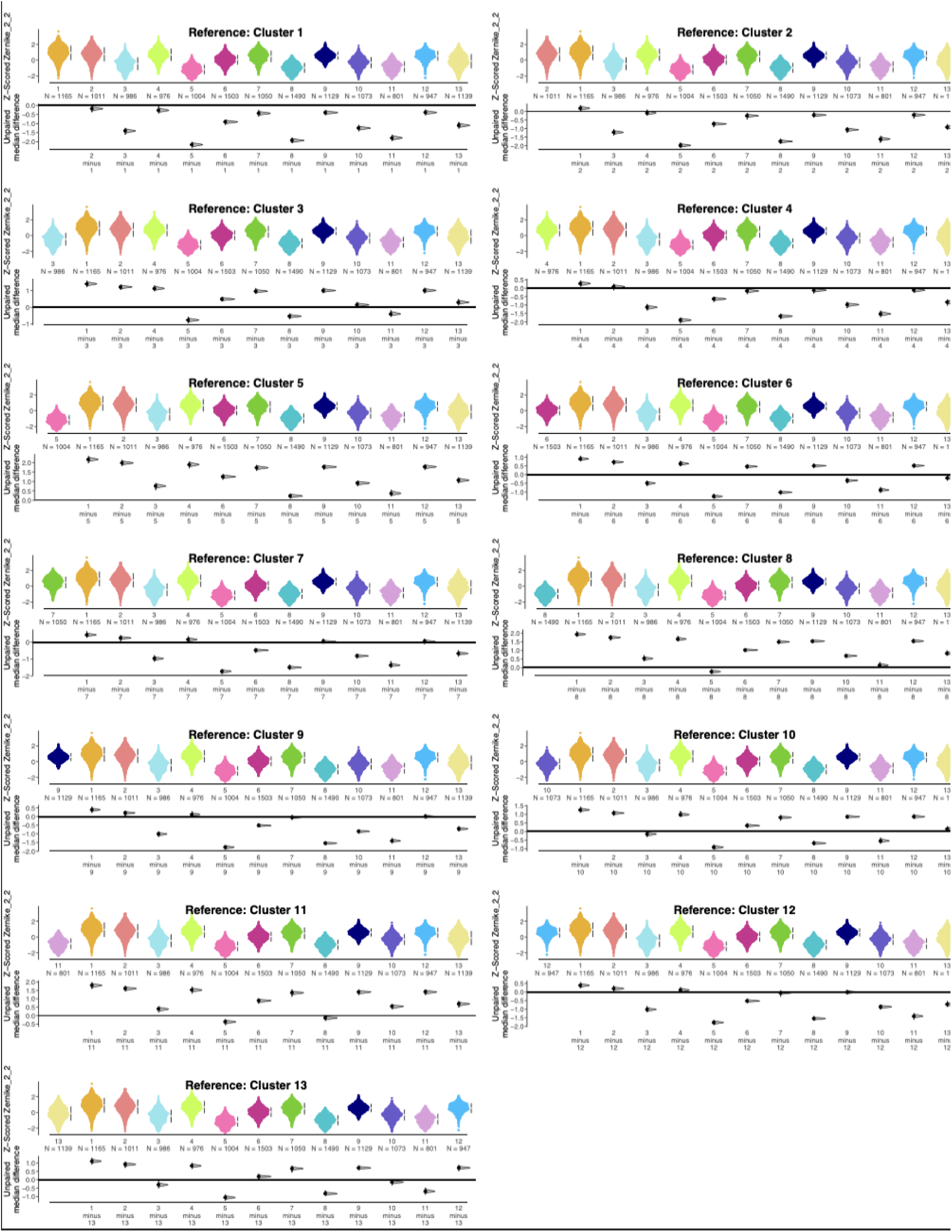
Zernike Moment 2_2 values among morphology clusters of PV+ interneurons. Estimation statistics plots showing the distributions of z-scored Zernike 2_2 moment values for single PV+ interneurons grouped by morphology cluster. The collection of plots shows comparisons among the clusters using each morphology cluster as a reference. Beeswarm plots of the z-scored cell Zernike 2_2 are plotted on the upper axes of each plot. Beside each beeswarm the median z-scored Zernike 2_2 moments for each cluster is indicated as a white dot, and the 95% confidence interval about the median is shown by the black bar. The lower axes of each plot show the unpaired median difference plotted as a bootstrapped sampling distribution. The 95% confidence interval is shown as a black bar and the median is shown with a black dot.

**Figure 13.**
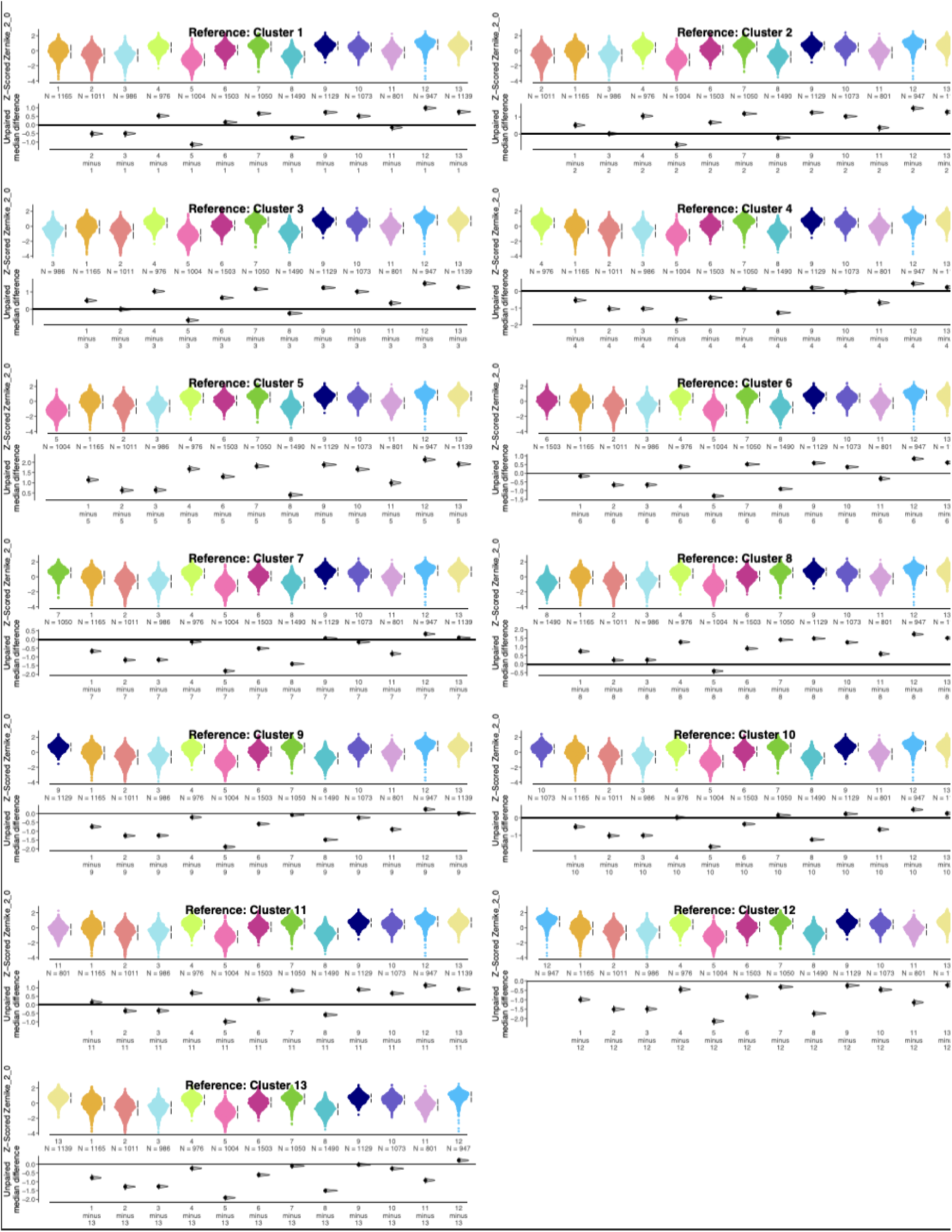
Zernike Moment 2_0 values among morphology clusters of PV+ interneurons. Estimation statistics plots showing the distributions of z-scored Zernike 2_0 moment values for single PV+ interneurons grouped by morphology cluster. The collection of plots shows comparisons among the clusters using each morphology cluster as a reference. Beeswarm plots of the z-scored cell Zernike 2_0 are plotted on the upper axes of each plot. Beside each beeswarm the median z-scored Zernike 2_0 values for each cluster is indicated as a white dot, and the 95% confidence interval about the median is shown by the black bar. The lower axes of each plot show the unpaired median difference plotted as a bootstrapped sampling distribution. The 95% confidence interval is shown as a black bar and the median is shown with a black dot.

#### Shape Motif 2 - pyriform cell bodies: clusters 3, 6, 9, 10, 13

These cells are defined by intermediate median measures of cell circularity and elongation (Fig.8A, Figs. 11-17). These cells have intermediate shape characteristics - they are not as elongated as cells in shape motif 1 and not as circular as cells in shape motif 3. Cells in shape motif 2 were round on one side and taper to a point. Their median z-scored shape measurement values were closer to 0 than those in the other shape motifs. These clusters were found in the middle of the morphology space, in between regions occupied by clusters with the other two shape motifs (Fig. 6). Relative to the other shape motifs, pyriform cells had lower measures of elongation than fusiform cells (Figs.11-13), and lower measures of circularity than round cells (Figs.14-17).

**Figure 14.**
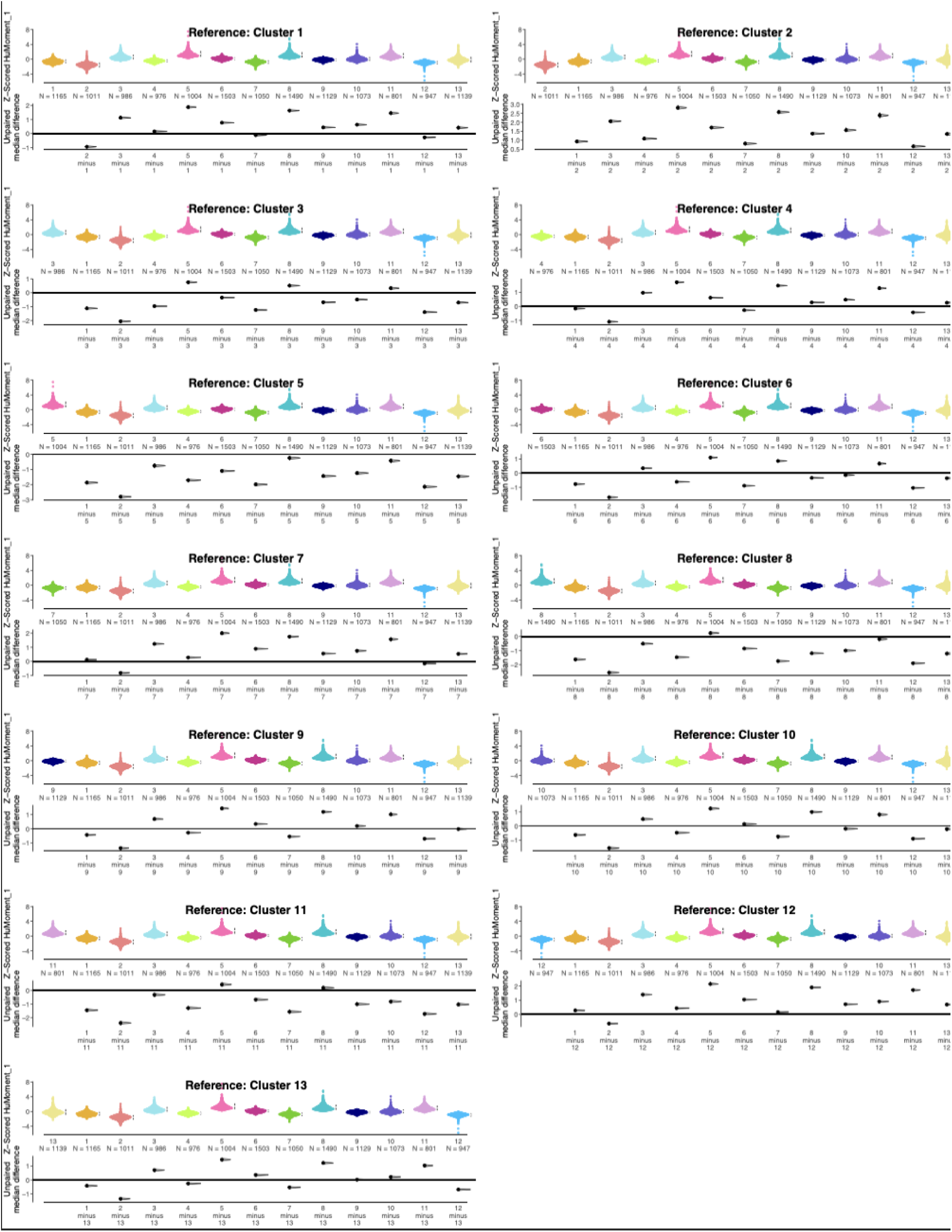
Hu Moment 1 values among morphology clusters of PV+ interneurons. Estimation statistics plots showing the distributions of z-scored Hu moment 1 values for single PV+ interneurons grouped by morphology cluster. The collection of plots shows comparisons among the clusters using each morphology cluster as a reference. Beeswarm plots of the z-scored cell Hu Moment 1 values are plotted on the upper axes of each plot. Beside each beeswarm the median z-scored Hu Moment 1 values for each cluster is indicated as a white dot, and the 95% confidence interval about the median is shown by the black bar. The lower axes of each plot show the unpaired median difference plotted as a bootstrapped sampling distribution. The 95% confidence interval is shown as a black bar and the median is shown with a black dot.

**Figure 15.**
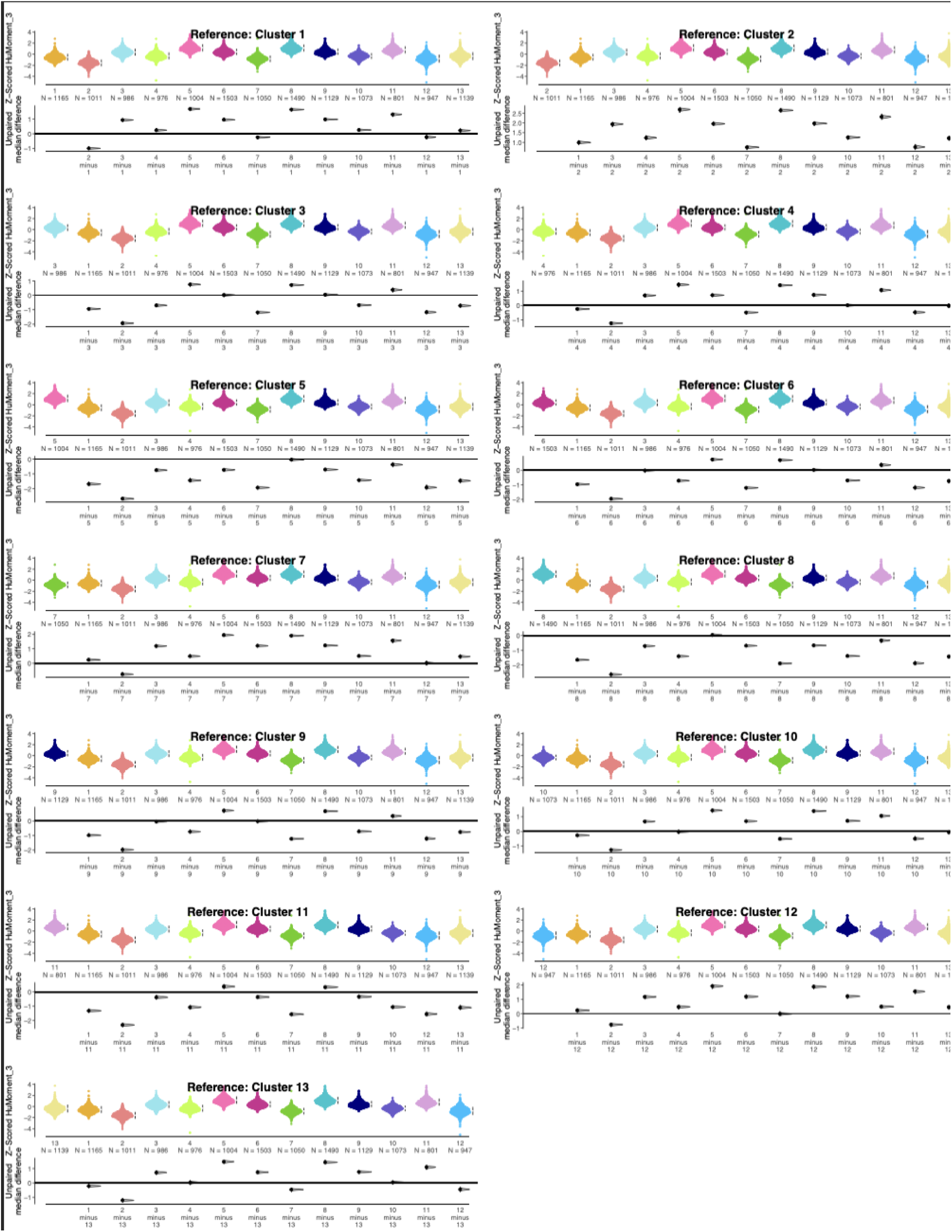
Hu Moment 3 values among morphology clusters of PV+ interneurons. Estimation statistics plots showing the distributions of z-scored Hu moment 3 values for single PV+ interneurons grouped by morphology cluster. The collection of plots shows comparisons among the clusters using each morphology cluster as a reference. Beeswarm plots of the z-scored cell Hu Moment 3 values are plotted on the upper axes of each plot. Beside each beeswarm the median z-scored Hu Moment 3 values for each cluster is indicated as a white dot, and the 95% confidence interval about the median is shown by the black bar. The lower axes of each plot show the unpaired median difference plotted as a bootstrapped sampling distribution. The 95% confidence interval is shown as a black bar and the median is shown with a black dot.

**Figure 16.**
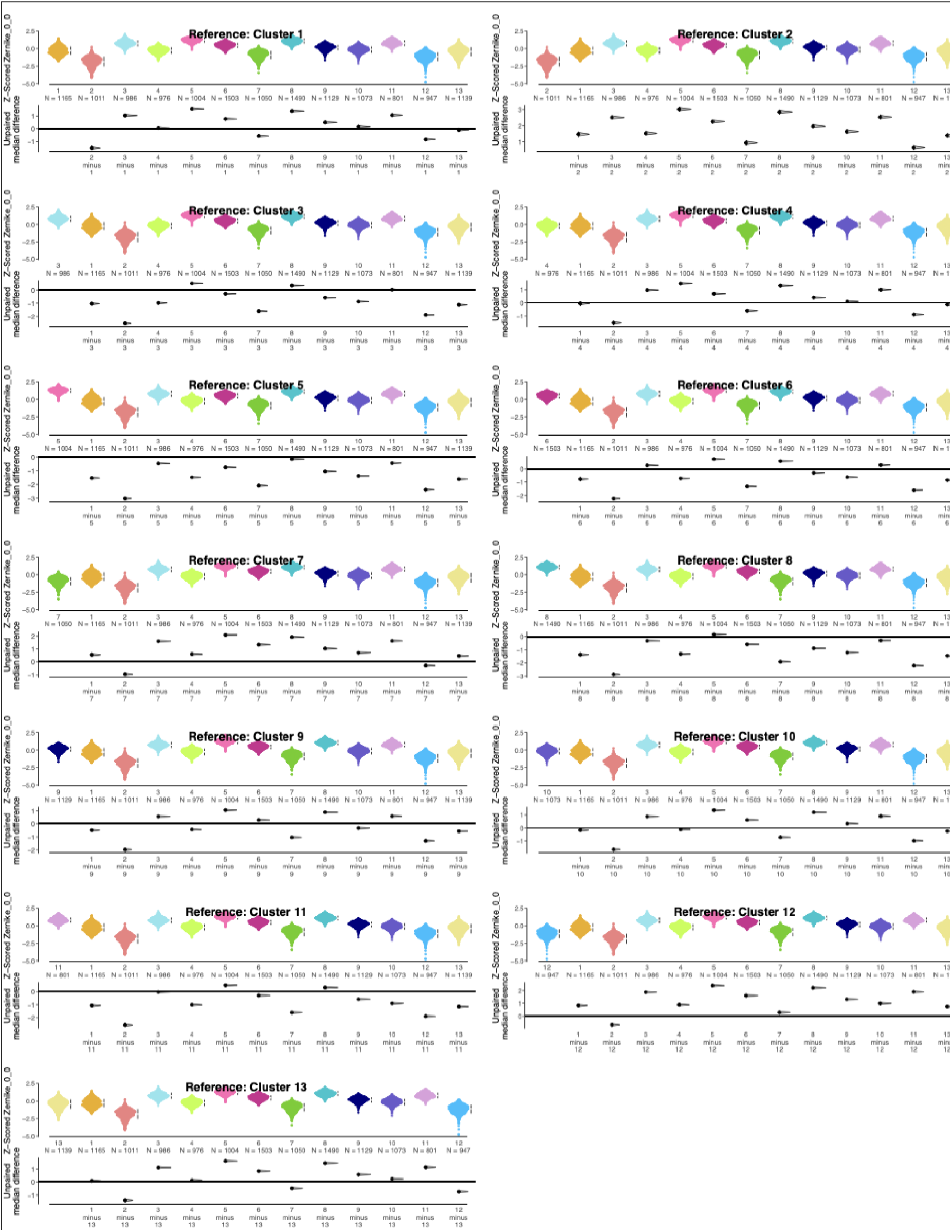
Zernike Moment 0_0 values among morphology clusters of PV+ interneurons. Estimation statistics plots showing the distributions of z-scored Zernike Moment 0_0 values for single PV+ interneurons grouped by morphology cluster. The collection of plots shows comparisons among the clusters using each morphology cluster as a reference. Beeswarm plots of the z-scored cell Zernike Moment 0_0 values are plotted on the upper axes of each plot. Beside each beeswarm the median z-scored Zernike moment 0_0 values for each cluster is indicated as a white dot, and the 95% confidence interval about the median is shown by the black bar. The lower axes of each plot show the unpaired median difference plotted as a bootstrapped sampling distribution. The 95% confidence interval is shown as a black bar and the median is shown with a black dot.

**Figure 17.**
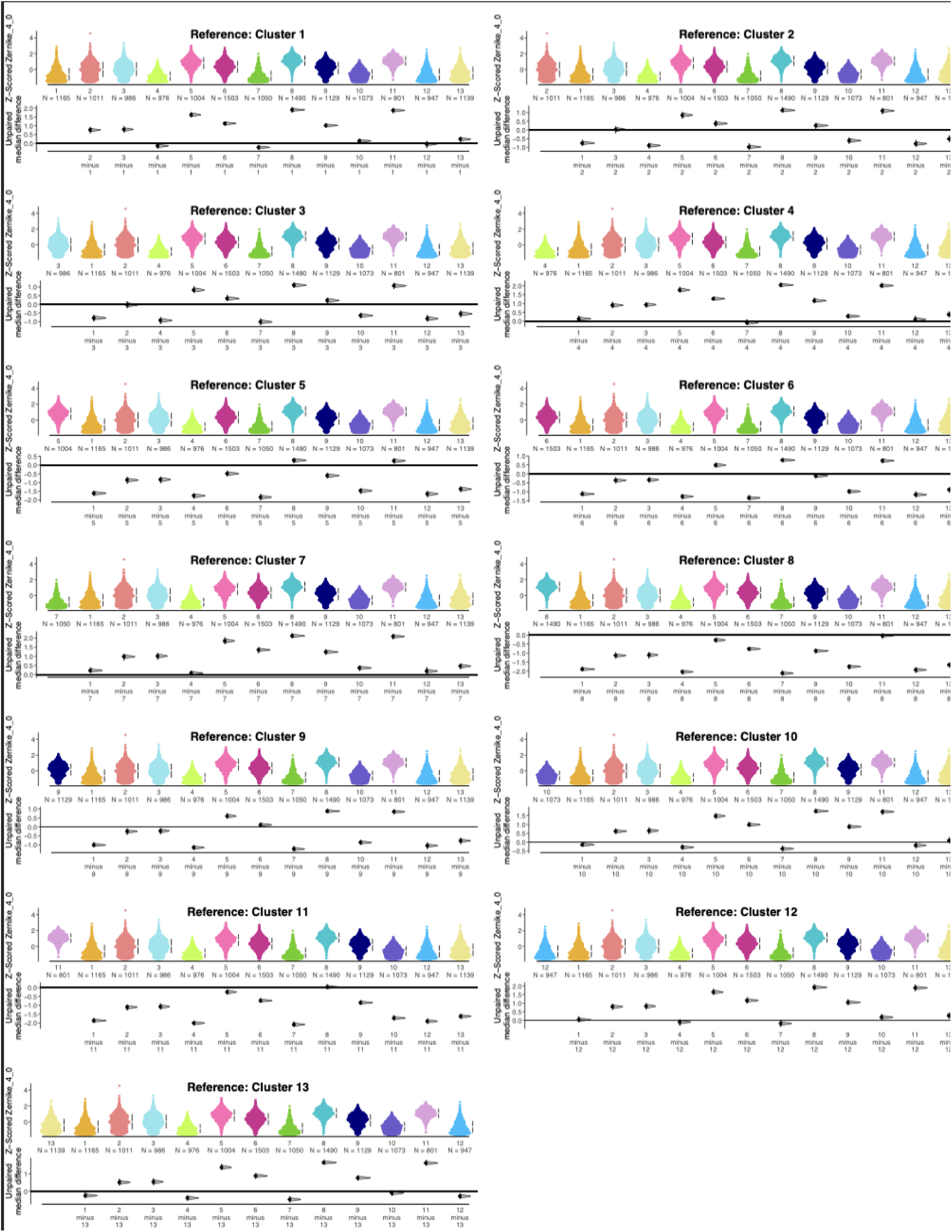
Zernike Moment 4_0 values among morphology clusters of PV+ interneurons. Estimation statistics plots showing the distributions of z-scored Zernike Moment 4_0 values for single PV+ interneurons grouped by morphology cluster. The collection of plots shows comparisons among the clusters using each morphology cluster as a reference. Beeswarm plots of the z-scored cell Zernike Moment 4_0 values are plotted on the upper axes of each plot. Beside each beeswarm the median z-scored Zernike moment 4_0 values for each cluster is indicated as a white dot, and the 95% confidence interval about the median is shown by the black bar. The lower axes of each plot show the unpaired median difference plotted as a bootstrapped sampling distribution. The 95% confidence interval is shown as a black bar and the median is shown with a black dot.

#### Shape Motif 3 - round cell bodies: clusters 5, 8, 11

Cells in these clusters are more circular with high median measures of circularity (Zernike_0_0, Hu Moments 1 and 3, Zernike_4_0) and lower measures of elongation. The largest cells, cluster 11, had radially symmetrical bumps around the perimeter of the soma that probably reflect processes emerging from the soma. The median Zernike_4_0 was high in cluster 11 and measured circularity and fourfold radial symmetry. As processes around the perimeter of the soma will tend to display greater symmetry in more circular somata. These clusters were visualized across the bottom of the morphology (Fig.7). Estimation statistics performed on measures of circularity identify clusters 5, 8, and 11 as having the highest median circularities of all clusters (Figs. 14-17).

### Cortical Area Distribution of PV+ Morphologies

We analyzed if any morphologies were more commonly found in S1 versus V1 by quantifying the proportion of cells from each area in a cluster (Fig. 7B) and assessed these differences with the chi-square test for goodness of fit. Among clusters 11-13, which have the largest median cell sizes across all clusters, more cells come from S1 than expected (global proportion S1:V1 = 62%:38%. Cluster 11: p = 5.546e-06, Cluster 12: p = 1.789e-09, Cluster 13 : p = 4.074e-14). Among clusters 3-6, which represent small and medium median cell sizes, more cells come from V1 than expected (global proportion S1:V1 = 62%:38%, Cluster 3 : p = 1.082e-06, Cluster 4 : p = 5.160e-06, Cluster 5 : p = 3.967e-04, Cluster 6 : p = 3.107e-04).

In contrast to measuring only the area of the cell body, the analysis of PV+ cells applied an unsupervised, data-driven approach to a high-dimensional representation of PV+ cell morphologies. The results of using our analysis pipeline indicate that measurements of both size and shape parameters are necessary to fully characterize the morphology of PV+ neurons. For example, we showed multiple shapes within each size division. Furthermore, the approach highlights the importance of using a large number of cells and morphology features to quantify the complexity of PV+ cell morphologies and gain insight into the distribution of PV+ cell morphology.

### Laminar Distribution of Morphology Clusters in S1 and V1

The spatial location of the cells in each cluster was mapped onto the standardized cortical space for S1 and V1. The laminar density profile was calculated for each cluster in each cortical area (Fig. 18). Visual inspection of these maps showed that none of the PV+ morphology clusters was laminarly restricted. Still, all of the clusters had a laminar location with peak PV+ cell density. Comparing the cortical areas, the same morphology cluster appeared to have different laminar patterns. For example, clusters 2 and 12 have both a supragranular and infragranular peak in S1. Cluster 2 has a more broad, diffuse distribution across the layers in V1, whereas cluster 12 is mainly found in supragranular layers.

**Figure 18.**
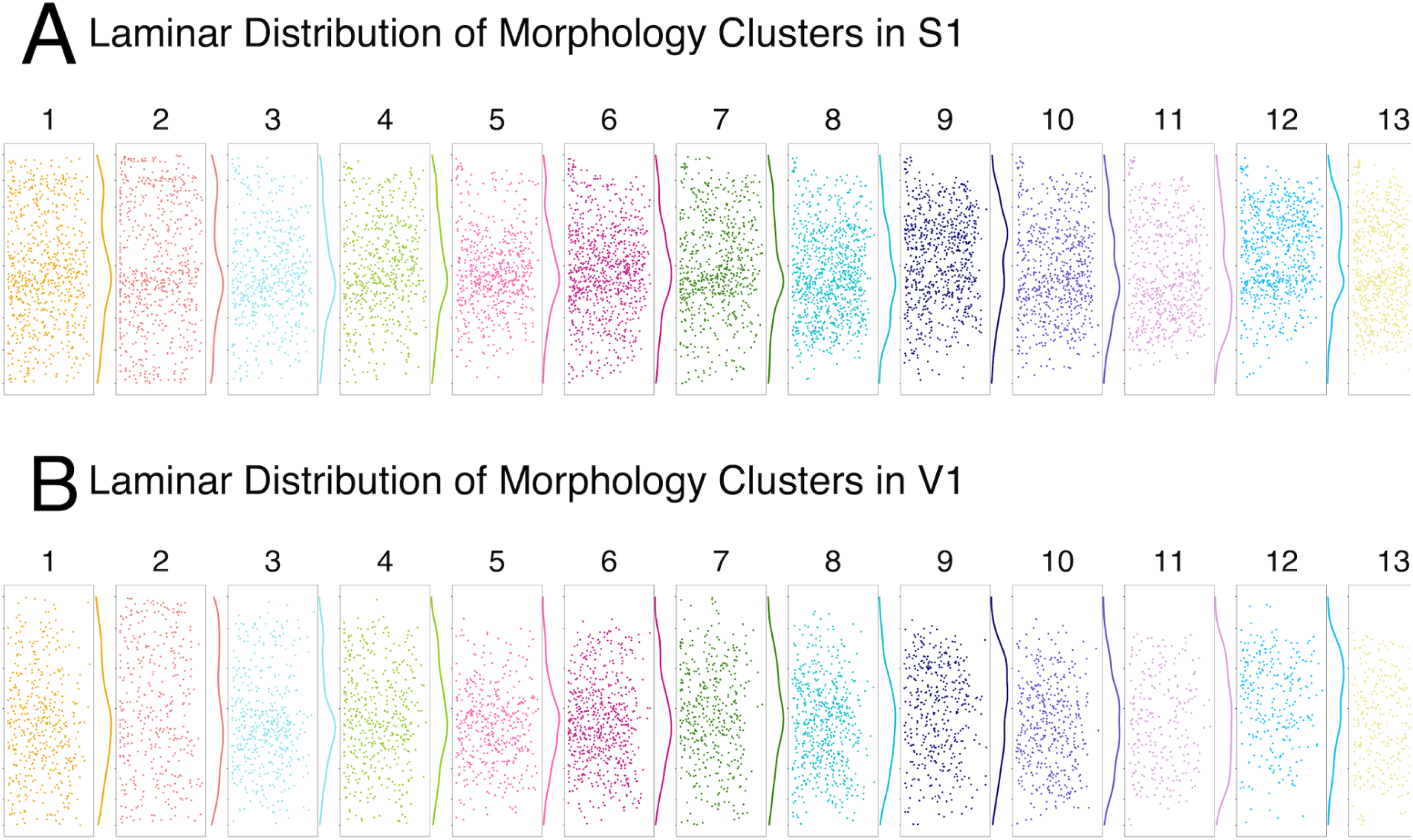
Laminar distributions of PV+ Morphology Clusters in S1 and V1. - For each cluster, cells (single points) are shown in standard coordinates in each cortical area. In a single plot, data from all 5 animals are shown. The layer boundaries are calculated from the analysis in Figure 3. Each cluster cell plot shows data pooled from all animals within a single cortical area. Beside each plot is the laminar profile for each morphology cluster, showing their distribution across the cortical layers for S1 and V1.

The laminar distributions of each morphology cluster in S1 and V1 are shown in Figure 19. We used chi-square analysis to test whether any clusters had more PV+ cells in a layer than expected. Except for clusters 4 and 10 in S1, and clusters 4, 7, and 10 in V1, all clusters had different proportions of cells by layer than expected. Notably, we found that in S1 and V1, cluster 2 had more cells from layers 1 and 6B, and cluster 12 had more cells from layers 4 and 5 than expected (S1 cluster 2: p= 3.112e-37, 12: p = 1.683e-16; V1 clusters 2: p = 3.396e-50, 12: p =3.238e-11).

**Figure 19.**
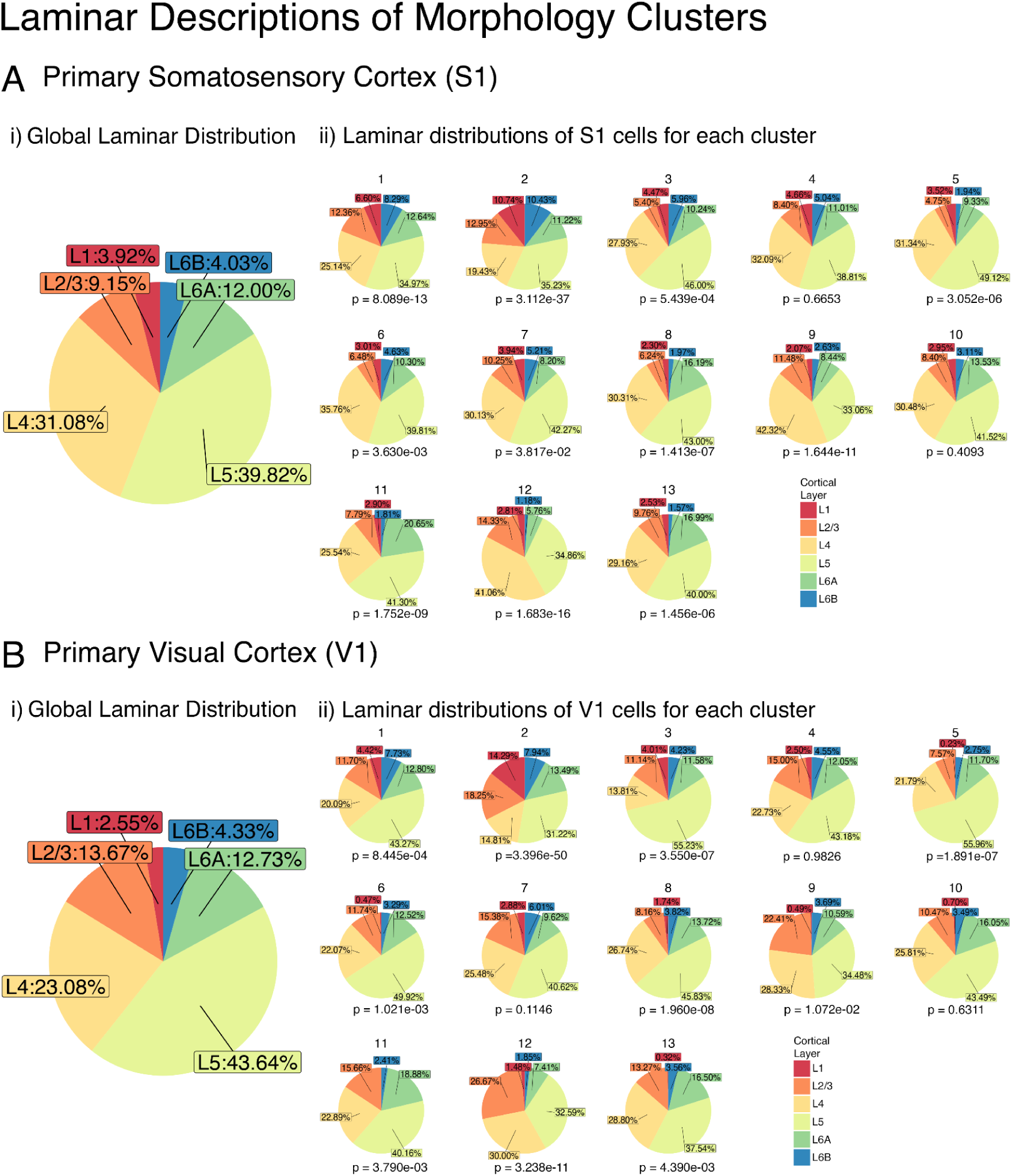
Proportions of PV+ cells by cortical layer for each morphology cluster. For S1 (A), and V1(B) the global proportions of cells were determined (i). For each cluster, in each cortical area, the proportions of cells by cortical layer were determined (ii). P-values are from the chi-square test for goodness of fit, comparing the laminar proportions in each morphology cluster to the overall laminar proportions in each cortical area.

### Unsupervised Analysis of PV+ Cell Laminar Patterns

To quantify the laminar patterns of each PV+ cell morphology cluster, we applied unsupervised hierarchical clustering to a dataset of 130 laminar density profiles (13 morphology clusters x 2 cortical areas x 5 animals). The clustering identified 6 different laminar patterns (Fig. 20A). We interrogated each laminar cluster to determine the proportion of profiles from S1 vs V1 and the 3 cell shape motifs. Finally, we used a river plot to connect the cells in each of the 13 PV+ morphology clusters with their laminar profile cluster (Fig. 20B).

**Figure 20.**
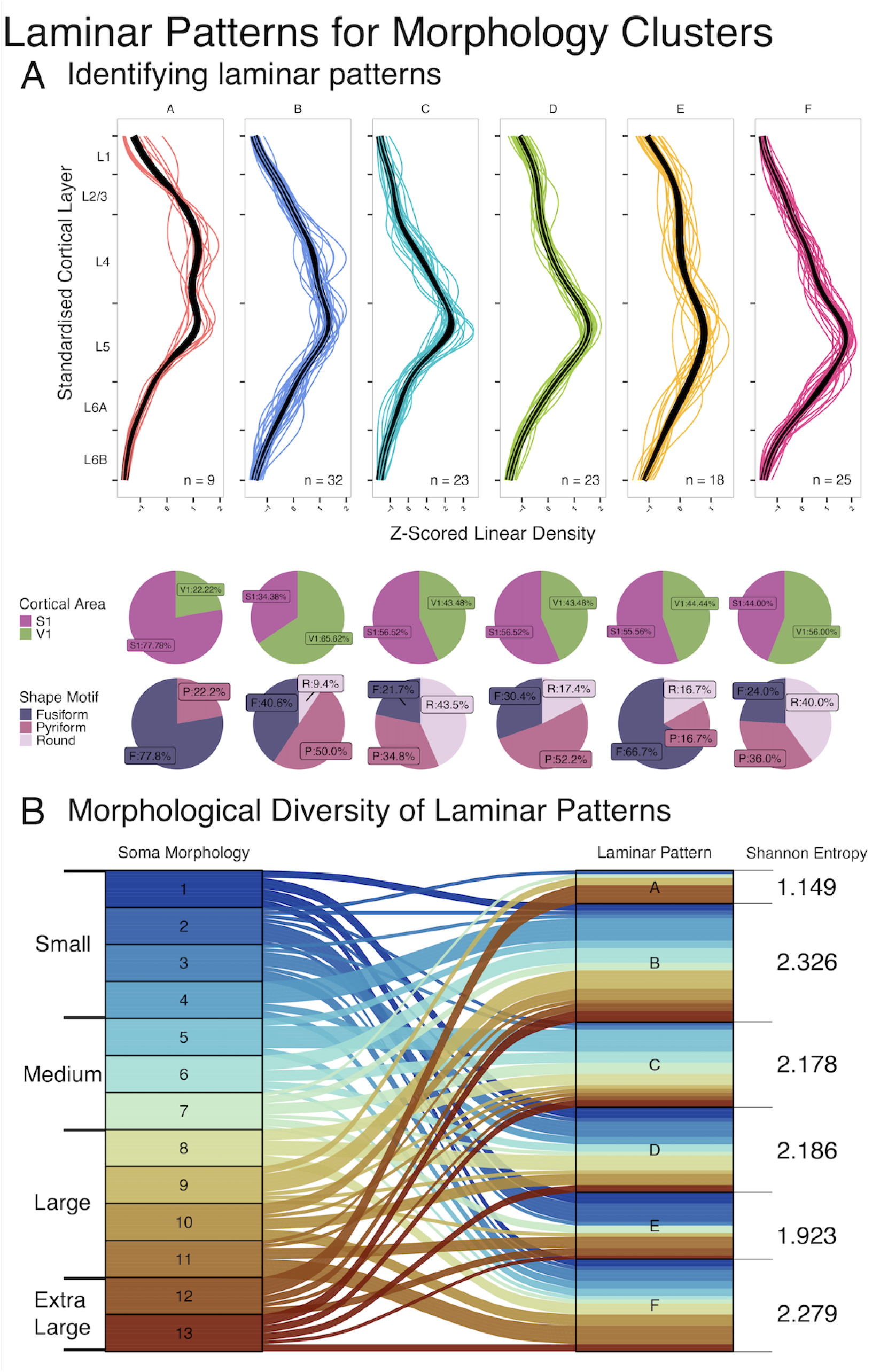
Characterizing the laminar distributions of PV+ cell morphologies. (A) 130 laminar profiles (5 animals x 2 cortical areas x 13 morphology clusters) were clustered into 6 patterns using unsupervised hierarchical clustering (ward.D2 algorithm). Cells from V1 were interpolated into S1 cortical space using a cubic piecewise spline fit so a single-layer definition could be used. Single-coloured profiles represent the laminar profile for a single morphology cluster in one cortical area from one animal. The average curve for each laminar pattern is plotted as a thick black line. Top row: pie charts summarizing the distribution of laminar profiles from each cortical area in each laminar pattern (S1: purple, V1: green). The proportions are compared to the overall distribution of profiles in the dataset (S1: 50%, V1:50%). Lower row: pie charts summarizing the distribution of shape motifs in each laminar pattern (F = Fusiform, P = Pyriform, R = Round). (B) River plot showing the correspondence between the laminar patterns (left column) and the morphology clusters (right column). Size divisions for the morphology clusters are shown to the left of the river plot. Shannon entropy values were calculated for each laminar pattern and are shown to the right of the river plot.

#### Laminar Pattern A

This laminar pattern describes a sharp increase in cell density across layers 2/3, before decreasing across layer 4 and increasing again in layer 5. This pattern is found slightly more in S1 compared to V1 (S1:V1 = 77.78%:22.22%, p = 9.588e-02). This pattern consists primarily of extra-large fusiform cells (Cluster 12). This pattern is morphologically restricted and biased towards a single morphology (p = 4.62e-02), as it has a Shannon entropy score of 1.149.

#### Laminar Pattern B

This laminar pattern increases more gradually across layer ⅔ and continues to increase across layer 4 to peak cell density in layer 5. This pattern is found slightly more in V1(S1:V1 = 34.38%:65.62%, p = 7.710e-02). Compared to S1, where large, fusiform PV+ cells populate supragranular layers (Fig. 8A), in V1, medium-sized pyriform cells populate these layers(Fig. 8B). Unlike pattern A, this laminar pattern is morphologically broad, with less morphological bias within the pattern (Shannon entropy = 2.326).

#### Laminar Pattern C

This laminar pattern describes low cell density across the supragranular layers - layers 1 and 2/3. Cell density increases across layers 4 and upper layer 5 before decreasing across the deeper sublaminae of layer 5. This pattern is found slightly more in S1 than V1, but does not differ strongly from expected (S1: V1 = 56.52%, 43.48%), p = 5.316e-01. This pattern is the only one to consist mostly of round cells (43.5% round cells, p = 5.236e-02). Morphology clusters with medium-sized cells (Clusters 5, 6, and 7) are in this pattern. This pattern is also morphologically diverse with less morphology bias (Shannon Entropy = 2.178).

#### Laminar Pattern D

Compared to pattern C, which also has a peak in L5, pattern D has increased cell density across the supragranular layers, including layer 1. Cell density increases progressively along the depth of layer 4, with a peak towards the middle of layer 5. This pattern and the latter two show increased cell densities into L6A, which characterize the deep infragranular biases seen across some of the morphology clusters shown previously. This pattern consists primarily of pyriform cells, with slightly more than expected (52.2% pyriform cells, p = 4.000e-01). This pattern also consists of morphology clusters with mainly medium to large sizes.

#### Laminar Pattern E

This pattern, like pattern A, increases cell density across layers ⅔, and 5. However, the increase in cell density is more gradual across the supragranular layers than in pattern A. In pattern E, cell density is sustained from layer ⅔ across layer 4 before increasing again across layer 5. This pattern does not show a sharp decrease in cell density across layer 4, as seen in pattern A. This pattern is found slightly more in S1(S1:V1 = 55.56%, 44.44%, p = 6.374e-01) and consists mainly of small, fusiform cells (66.7% fusiform, p = 4.356e-02). With the second-lowest Shannon entropy of all the laminar patterns, this pattern is also morphologically biased and seen across fewer morphology clusters. As this pattern is dominated by small fusiform cells, cells from clusters 1 and 2 are more represented in this pattern than in others.

#### Laminar Pattern F

Like patterns D and E, pattern F gradually increases cell density across layers 2/3, 4 and 5 before decreasing in L6A. This pattern has the deepest peak cell density in Layer 5 of all the laminar patterns. Found slightly more in V1, this pattern has a similar proportion of fusiform, round and pyriform cells as the distribution of shape motifs in pattern C, with the greatest share of cells being round (40% round cells, p = 1.054e-01). This pattern has a similar composition of sizes compared to pattern C, which also consists mainly of medium cell sizes with a combination of all three shape motifs. However, this pattern consists of larger cells than pattern C’s composition. This laminar pattern is also morphologically broad with comparably low morphological bias (Shannon entropy = 2.279).

We found weak laminar differences when we applied the typical method that uses only cell size to assess laminar differences (Fig. 4). In contrast, we identified robust differences in the laminar patterns of PV+ cell morphologies using density profiles and unsupervised clustering. Layers with the greatest PV+ cell density were found to have comparable, overlapping size distributions, indicating high similarity. The approach, however, identifies variations in shape among cells of similar sizes and leverages the variations in shape in addition to size differences to characterize morphologies of single PV+ cells. Although strong areal biases were only seen for extra-large cells with these more comprehensive morphology clusters, the laminar analysis of PV+ morphologies identified laminar patterns biased towards either V1 or S1. Our data-driven approach considers both the size and shape information of single PV+ cells. The spatial distributions of these morphology clusters follow characteristic patterns that are more representative of S1 and V1.

### Comparing Cell Body Morphology and Dendritic Morphology

We analyzed the soma and dendritic morphology of whole-cell filled PV+ cells using data from mouse V1 from the Allen Brain Institute (Hodge et al., 2019). The first step in integrating these data with the PV+ ISH dataset was to use estimation statistics to compare 3 size (Cell Body Area, Bounding Box Area, and Perimeter) and shape (Circularity, Aspect Ratio and Eccentricity) parameters from the whole-cell filled cells and ensure that they were not different from the V1 PV+ dataset (Fig. 21). None of those measured differed between the PV+ whole-cell filled and ISH-labelled cells so we integrate the morphometric data from those cells. Combining data from two modalities, we used this dataset to assess the relationship between dendritic and soma morphologies in V1. This dataset comprises 5451 cells labelled by ISH and 36 filled cells from mouse V1. For the 36 filled cells, 20 measurements quantifying the dendritic morphologies were obtained from the Allen database. We used RSKC to cluster the cells separately based on their dendritic and soma morphologies.

**Figure 21.**
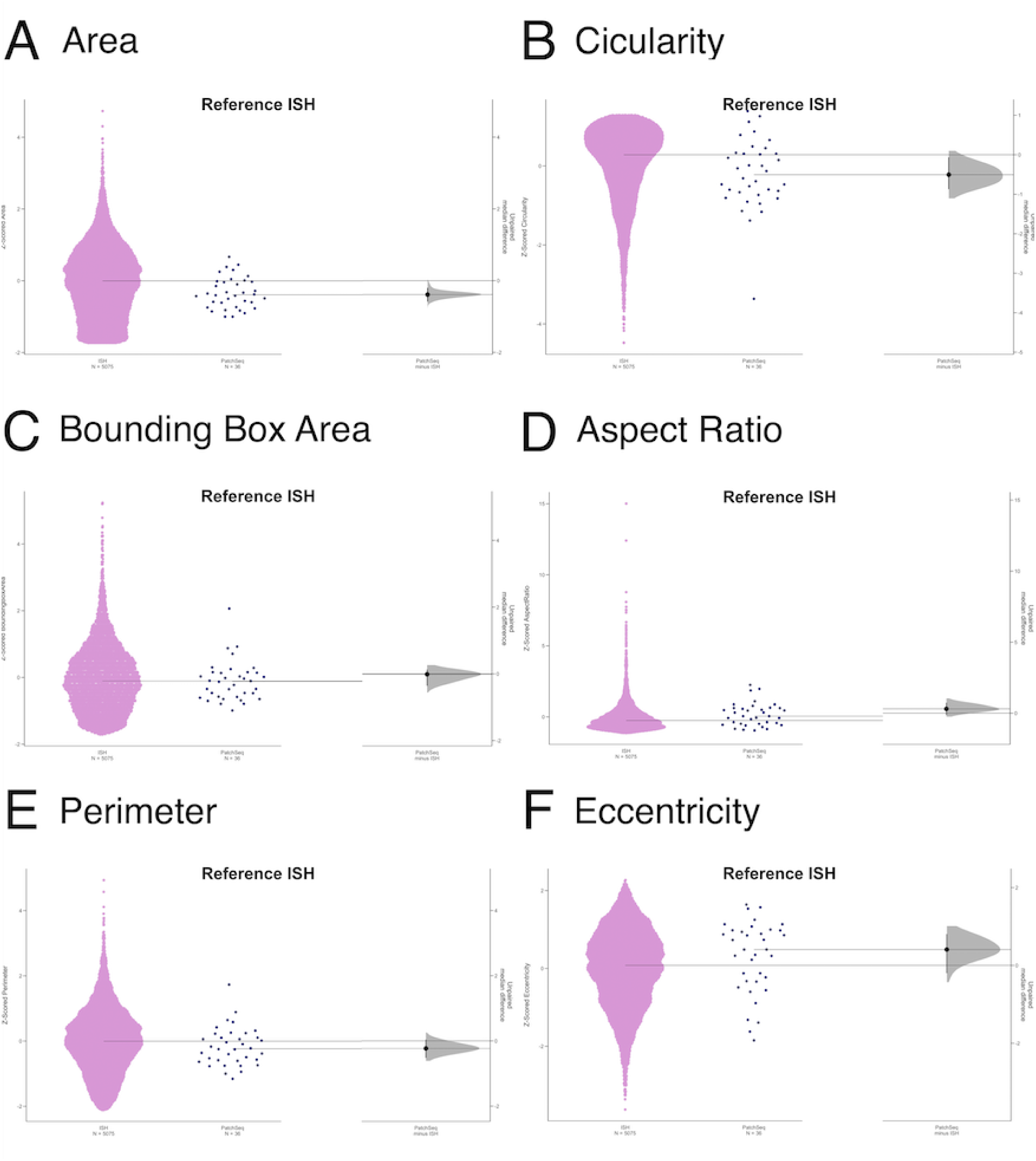
Filled cells have comparable size and shape parameters to ISH-labelled cells. Gardner-Altman estimation plots showing the unpaired median difference between ISH labelled cells and filled cells. ISH-labelled cells are being used as a reference. Both groups are plotted on the left axes, and the unpaired median difference is plotted on the right axes as a bootstrapped sampling distribution. The median difference is plotted as a black dot, and the 95% confidence interval is indicated by the vertical black line (A) Area, (B) Circularity, (C) Bounding Box Area, (D) Aspect Ratio, (E) Perimeter, (F) Eccentricity

#### Cluster Analysis of Dendritic Morphologies

The 20 dendrite morphology measures were used to cluster the 36 filled cells. The elbow method determined that the optimal k=3 and RSKC was used to cluster the cells based on the dendritic morphology features (Fig. 22A). The top-weighted dendritic morphology features included the number of nodes, the total length of the dendritic arbour, the average contraction, average diameter, total size of the dendritic field, and the number of tips (Fig. 22B). These features describe the size of the dendritic field, together with large-scale shape properties related to the arrangements of dendrites within the field. The lowest-weighted features include the soma size, average bifurcation angle, and branch order. Although information about the cell body was present in this cluster analysis, the only parameter provided was the size. The clusters obtained here using dendritic morphology features (Figs. 23 and 24) are consistent with previously published data (Jiang et al., 2015).

**Figure 22.**
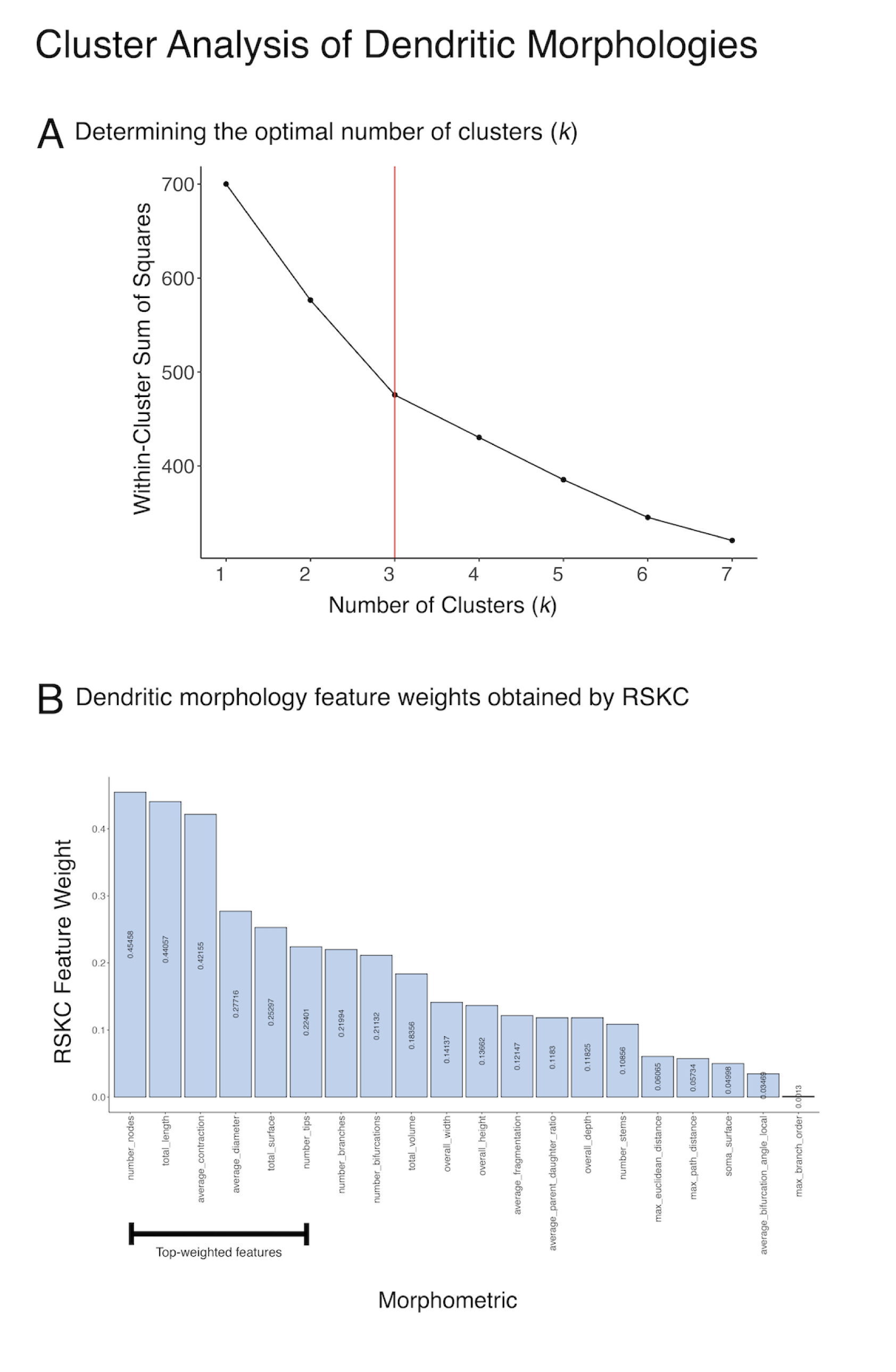
Clustering dendritic morphology features for PV+ filled cells. (A) Elbow plot showing the within-cluster sum of squares plotted as a function of the number of clusters (k). The optimal number of clusters (k= 3) is shown in red. (B) Feature weights as determined by RSKC for the 20 dendritic morphology features. Features quantify dendritic arbour morphologies, and top weighted (>50% maximum weight assigned to a single feature) features are indicated with the black bar.

**Figure 23.**
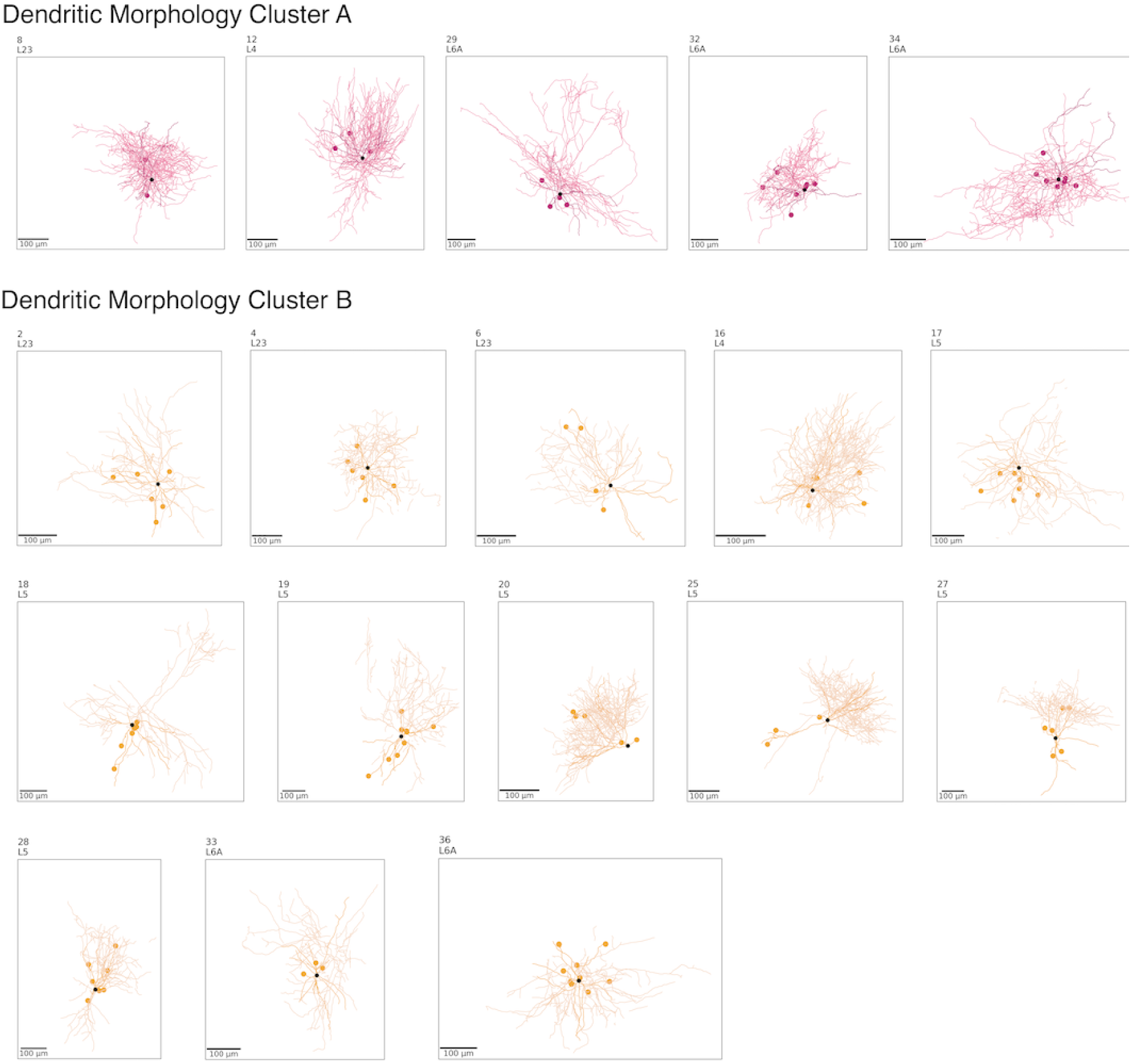
Cell Reconstructions of Filled cells (Dendritic Morphologies A and B). Whole cell reconstructions for each filled cell for the first two of three dendritic morphology clusters. Pale segments are the dendritic arbour, and dark segments are the axonal arbour. Truncated portions of arbours are indicated with a coloured dot. The black dot indicates the soma. Cells are numbered from 1-36 and are ordered by their cortical layer. Scale bars in all reconstructions = 100 microns. The different colors indicate the different dendritic morphology clusters (A = pink, B = gold).

**Figure 24.**
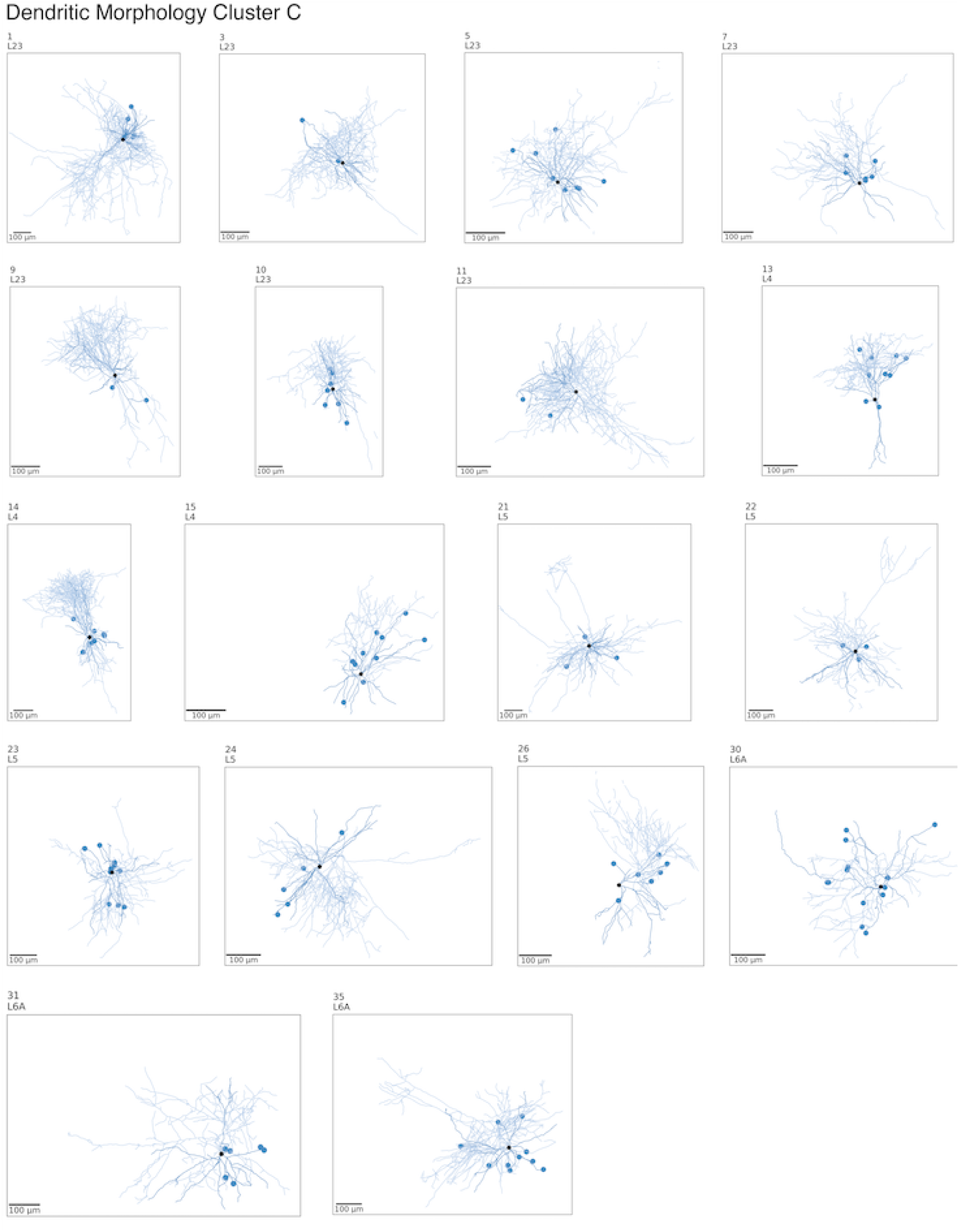
Cell Reconstructions of Filled cells (Dendritic Morphology C). Whole cell reconstructions for each filled cell for the last of three dendritic morphology clusters. Pale segments are the dendritic arbour, and dark segments are the axonal arbour. Truncated portions of arbours are indicated with a coloured dot. The black dot indicates the soma. Cell numbers indicate their position in the cortical layers, lower numbers indicating upper layers and higher numbers indicating deeper layers. Scale bar for all images = 100 microns.

Next, cell body morphology measurements of the PV+ whole-cell filled and ISH cells from V1 and RSKC were applied to cluster those data. Eleven cell body morphology clusters were identified, and the PV+ whole-cell filled cells were found in 7 of those clusters (Figs. 25 & 26). The top-weighted features were similar to those found when clustering only the PV+ ISH-labeled cells and included a combination of size and shape parameters (Fig. 25B). Among the top-weighted shape features were measures of elongation (Eccentricity, Zernike Moments 2_0 and 2_2, and the aspect ratio) and circularity (Zernike Moments 0_0 and 4_0, together with Hu Moments 1 and 3).

**Figure 25.**
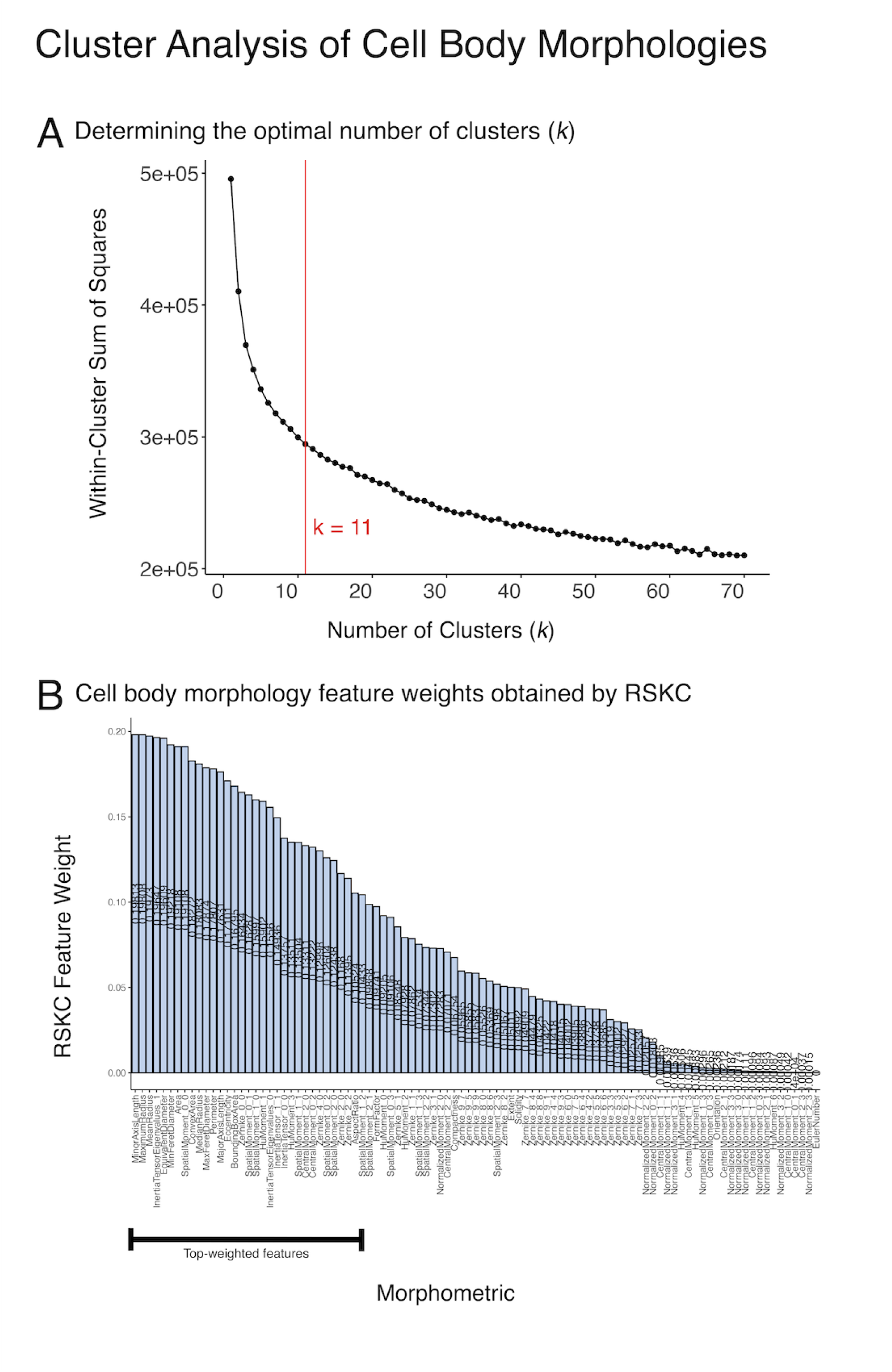
Clustering cell body morphologies of PV+ filled and ISH-labelled cells. (A) - Elbow plot showing the optimal number of clusters (k =11), (B) Feature weights as determined by RSKC performed on 97 morphology features for each cell soma. Top-weighted features (>50% of the maximum weight assigned to a single feature) are indicated with the black bar.

**Figure 26.**
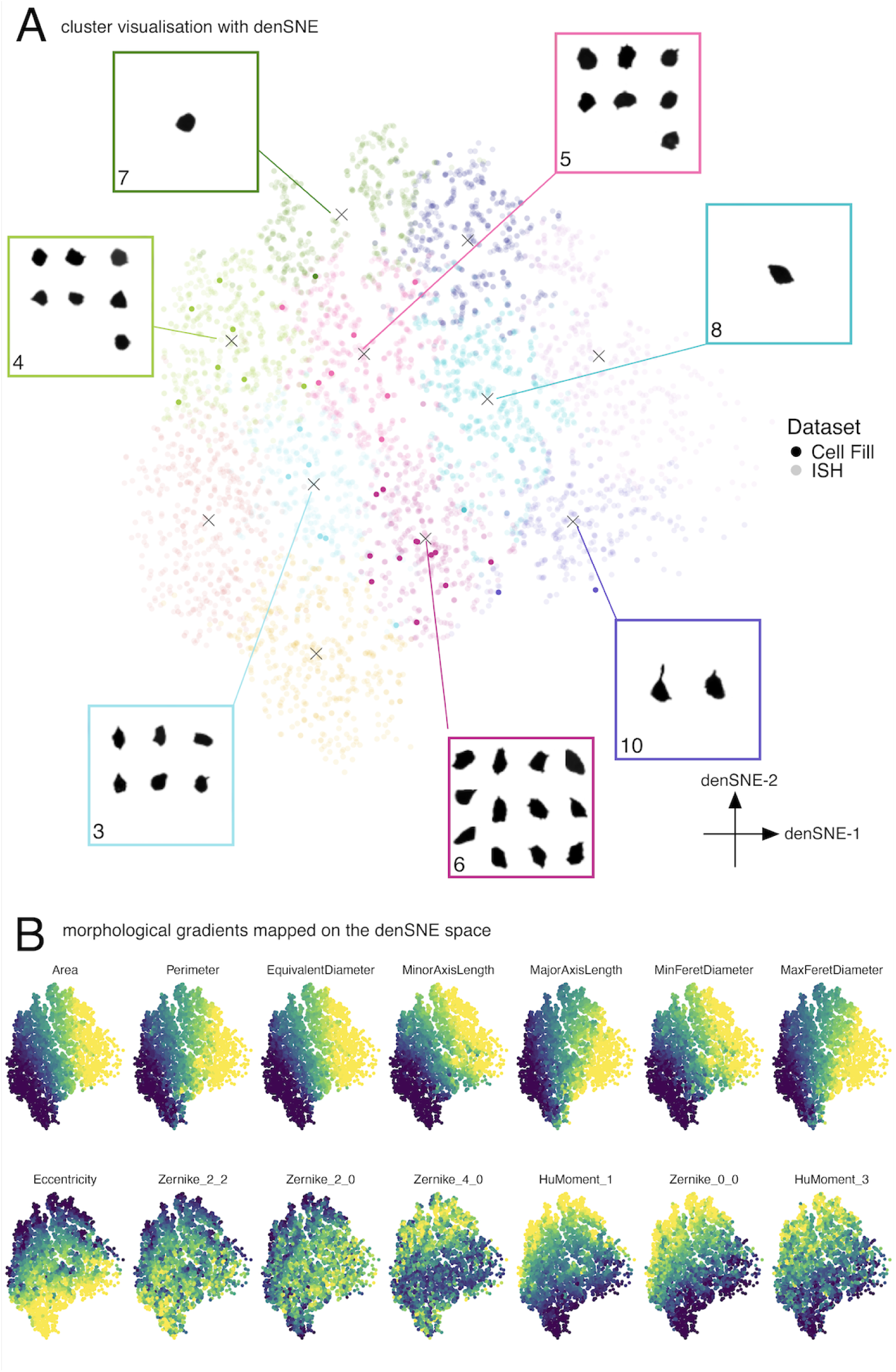
Morphological properties of filled cells integrate along the morphology gradients established by ISH-labelled cells. (A) - denSNE plot of the PV+ cell body morphology space annotated by cluster assignment. Cluster numbers indicated the progression of median cell size (Cluster 1 = lowest median cell size Cluster 11 = highest median cell size). Single points are single cells. Fully opaque points are whole-filled cells, and translucent points are ISH-labelled cells. For morphology clusters with filled cells, the cell bodies of those filled cells are shown (B) denSNE plot from panel A, but annotated for six top-weighted size features (top row) and six top-weighted shape features (bottom row). The colour progression from dark blue to yellow indicates increasing feature values for each morphology feature.

The 36 filled cells were found in 7 clusters along PV+ ISH-labelled cells (Fig. 26). Similar to the analysis of PV+ ISH cells, the denSNE plot represented roughly orthogonal gradients of size and shape. The filled cells were small to medium size and distributed across the range of shape patterns (Fig. 26B). For example, filled cells found in clusters 3, 4, and 6 represent a band of medium-sized cells in the denSNE plot but cut across a range of cell shapes from round in cluster 4 (dark blue region in the denSNE for eccentricity) to more and more oval-shaped cells in clusters 3 and 6 (Fig. 26A).

The heatmap of the top-weighted features for the filled and ISH PV+ cells from mouse V1 provides a clear visualization of the morphological features that separate cells into clusters and define morphological phenotypes. For example, clusters 3 and 4 had similar cells (greenish colour-coding of the size features), but the shape features had opposite patterns. The eccentricity of cluster 3 was higher (reddish colour-code), indicating round cells, while cluster 4 had lower eccentricity (bluish colour-code), indicative of not round cell shapes (Fig. 27).

**Figure 27.**
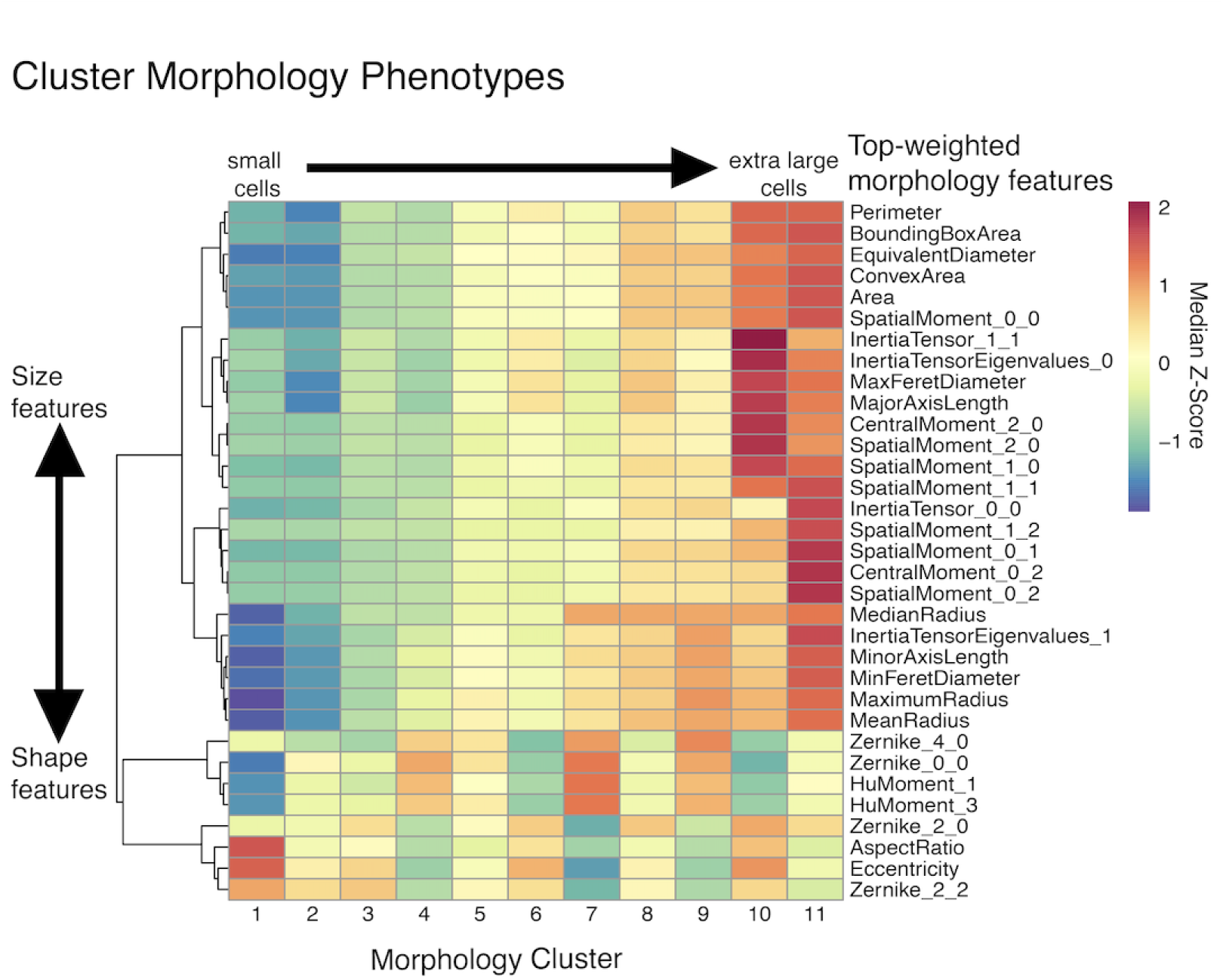
Morphological phenotypes for clusters of PV+ ISH-labelled cells and Whole-Filled cells. Heatmap of median z-scored feature values for top-weighted features for each morphology cluster. Cluster numbers 1-11 reflect size progression, and are related to the median cell size in each cluster (Cluster 1 = smallest cells, Cluster 11 = largest cells). The rows in the phenotype heatmap were organized by hierarchical clustering (Ward.D2 algorithm) to make the separation of size and shape parameters evident. The separation is shown by the row-wise dendrogram. The first branch point of the dendrogram indicates the separation of size parameters towards the top of the heatmap, and shape parameters towards the bottom of the heatmap. Furthermore across the size gradient, repeated shape motifs can be seen.

We used a river plot to map the correspondence between the clustering of PV+ filled cells based on dendritic versus soma morphologies (Fig. 28). Dendritic morphology clusters did not restrict themselves to exclusive combinations of cell body morphology clusters. Dendritic morphology A is primarily seen across the smaller, round PV+ cells from deep layers, but smaller fusiform PV+ cells in the supragranular and granular layers. This dendritic morphology pattern was not identified in the cells from layer 5. Dendritic morphology B is morphologically broad across cell body morphology clusters 3, 4, 5, and 6. These cells encompass a range of sizes and shapes (small to medium sizes and all three main shapes - fusiform, pyriform and round). Despite various cell body morphologies, this dendritic morphology is concentrated across layer 5 cells. Dendritic morphology C is also very morphologically and laminarly broad, found across cell body morphology clusters 4, 5, 6, 7, and 10, with the greatest proportions of cells falling into cell body morphology clusters 5 and 6 (pyriform and fusiform cells respectively). This dendritic morphology pattern is more representative of cells with less rounded cell bodies and is found across the layers in larger proportions than other morphology clusters.

**Figure 28.**
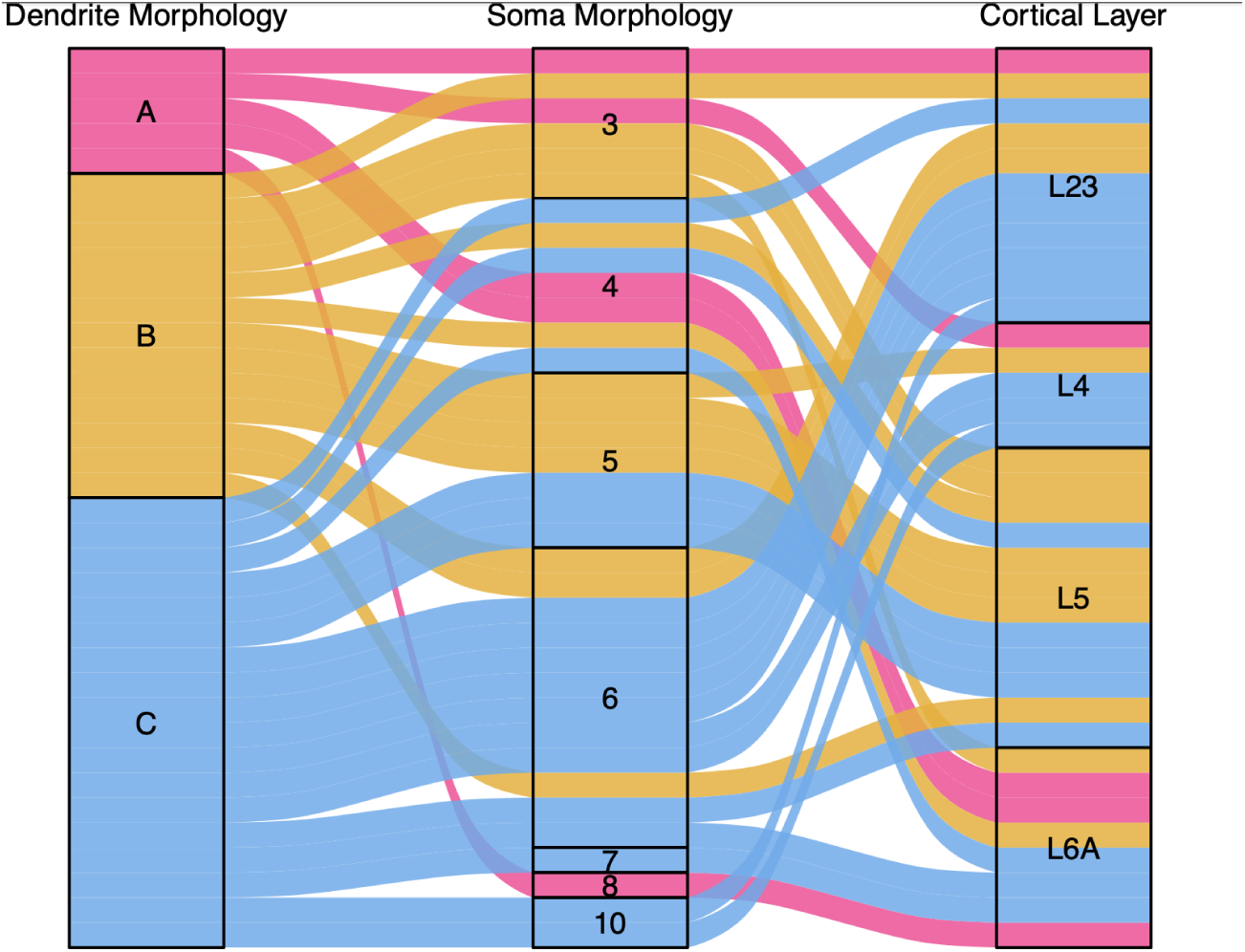
Correspondence among cell dendritic and cell body morphologies. River plot showing the correspondence among dendritic morphology clusters (left), cell body morphology cluster (centre) and cortical layer (right) for the 36 filled PV+ cells. Cell body morphology cluster labels reflect their position along the size gradient defined across all morphology clusters.

### Comparing the Morphology of PV+ and PV+ Associated Gene Cells

We compared the morphologies of PV+ cells with cells labelled for PV-associated genes from different T-types to determine if cell morphology could help identify PV+ T-types. The 13 PV-associated genes were selected from the list of marker genes for PV+ T-types defined in Tasic et al. 2016 and represent the transcriptomic types defined in Gouwens et al., 2020. The ISH for these selected genes had good-quality labelling in both S1 and V1. Although we could not perform the spatial transcriptomics needed to validate PV+ cells across the T-types, it is likely that these cells are from different PV+ T-types. The PV-associated cells overlap with the PV+ ISH cells in their laminar locations, however, some genes (Calb1, Etv1) display characteristic laminar restrictions, whereas PV+ cells were found across all cortical layers. (Fig. 29).

**Figure 29.**
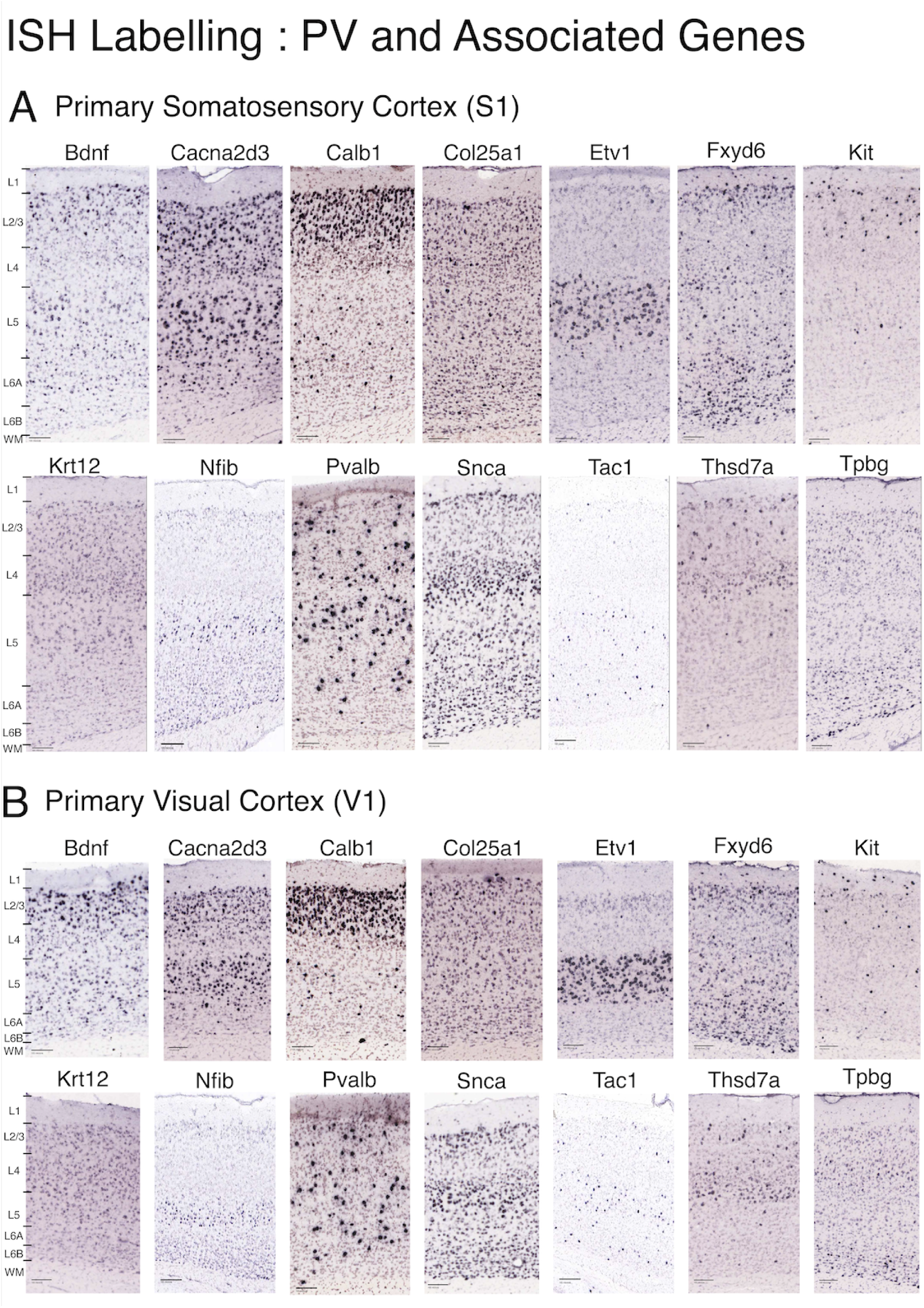
ISH samples for PV+ and Associated Genes. Samples of ISH extracted from (A) S1 and (B) V1 labelling a panel of 14 genes associated with PV+ transcriptomic types. Each gene’s ISH was extracted from a different animal, with S1 and V1 samples coming from a single section for each gene. The samples were cropped to the pial surface and leave a portion of the subcortical white matter visible beneath the deep limit of L6B. In all samples, only cortical cells were used in subsequent analyses. Pixel size : 1 pixel = 1 microns. Scale bar = 100 microns.

The morphology measurements from cells in S1 and V1 were clustered separately. In S1, 13 clusters were identified, and in V1, there were 10 clusters (Figs. 30A & 31A). The top-weighted features included size (area, perimeter, diameters, radii, axis lengths) and shape parameters (circularity – Zernike 0_0, Zernike 4_0, Hu Moments 1 and 3; elongation – eccentricity, Zernike 2_0, Zernike 2_2). Features with low weights in both cortical areas described the topology of the cells and higher-order shape moments indicated the presence of processes extending from the cell body (S1: Figure 30B, V1: Figure 31B). The larger number of clusters in S1 represented a greater range of cell sizes than V1.

**Figure 30.**
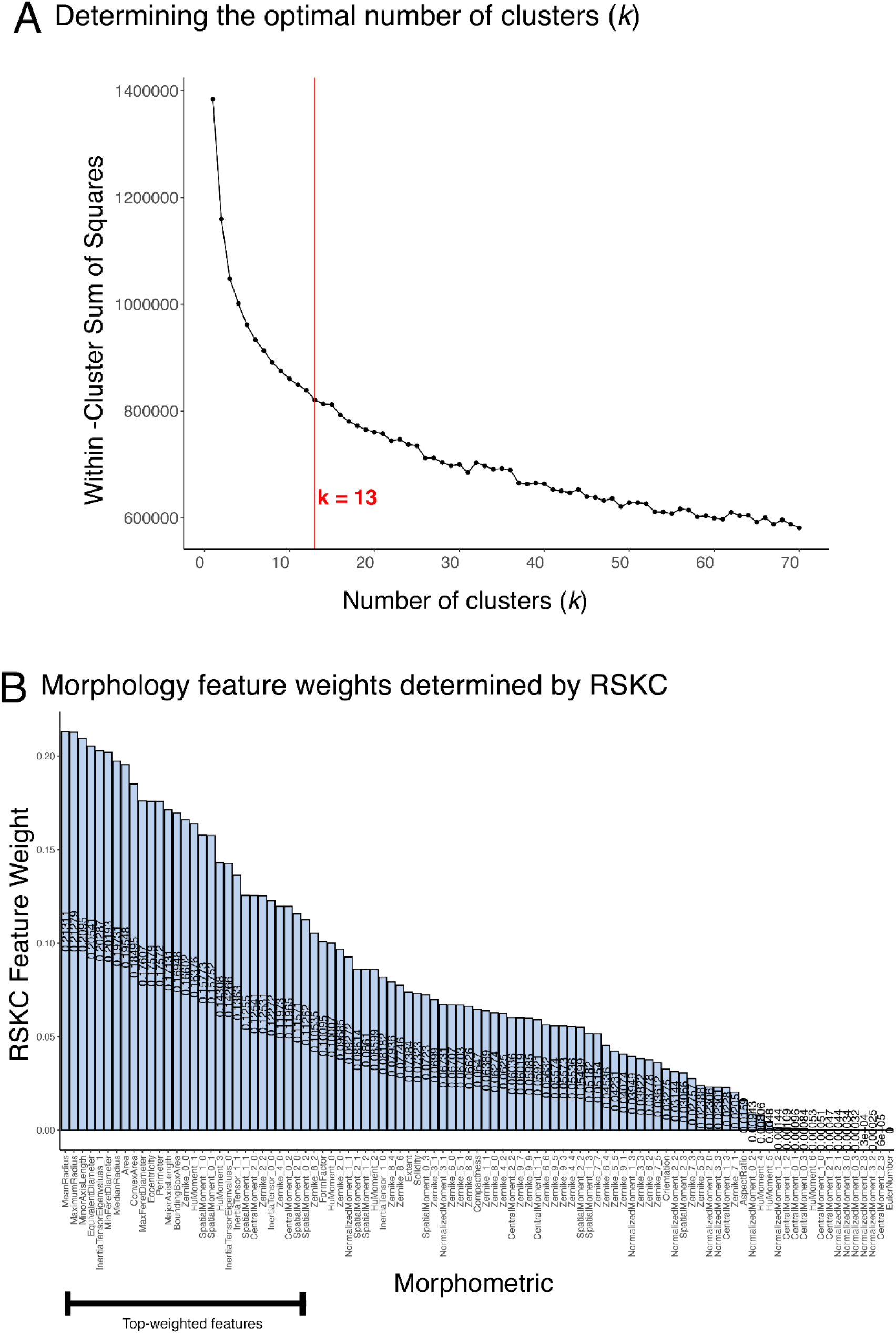
Clustering PV+ gene morphologies in primary somatosensory cortex S1. Weights plot showing the distribution of weight across the 97 morphology features as determined by the RSKC algorithm. Features are arranged so weights appear in decreasing order in the weights plot. Top weighted features (>50% max weight) are indicated by the black bar.

**Figure 31.**
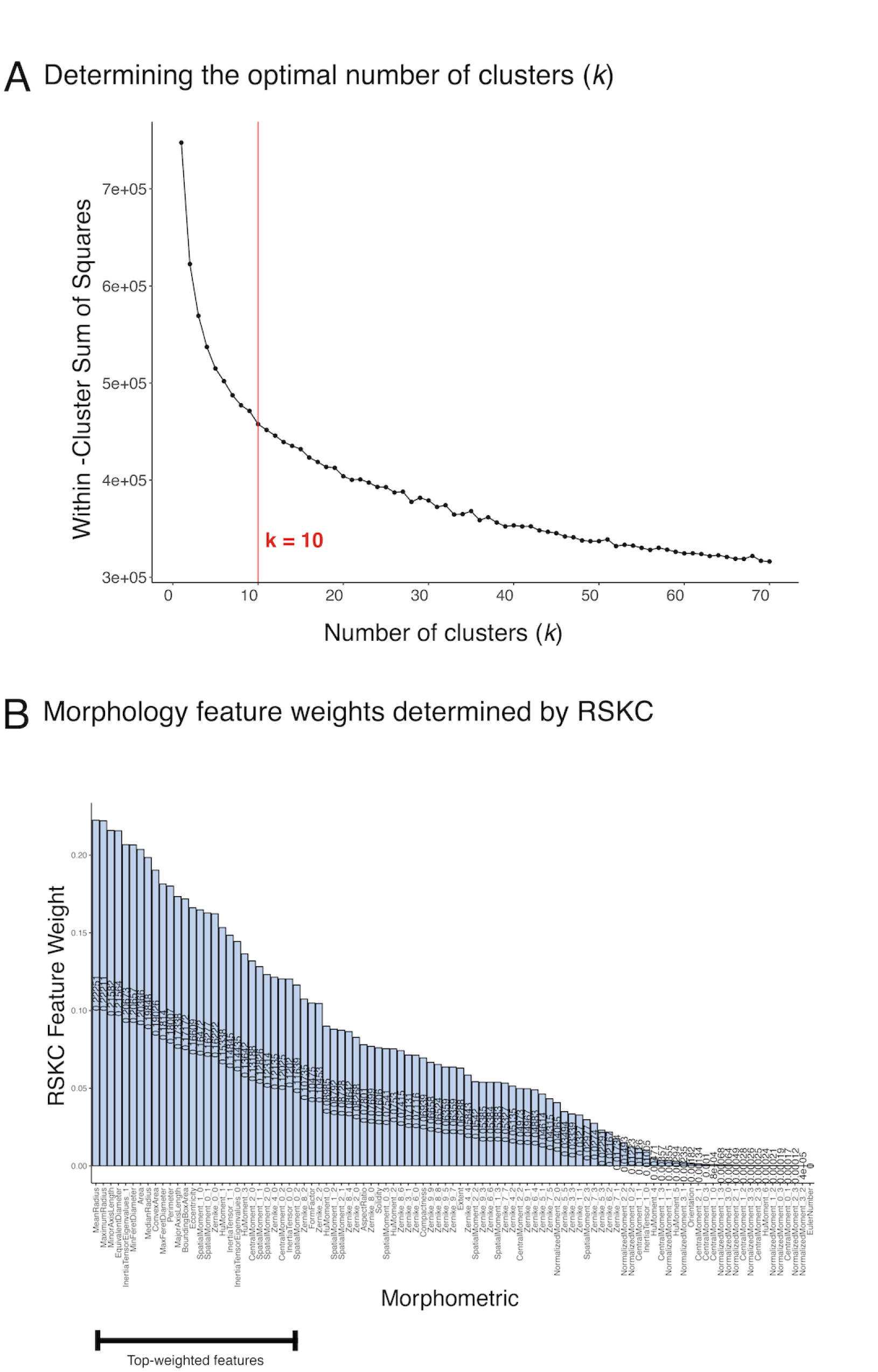
Clustering PV+ gene morphologies in primary visual cortex V1. (A) Elbow plot for the within cluster sum of squares as a function of the number of clusters (k). The optimal k = 10 is shown with the red line. (B) Weights plot showing the distribution of weight across the 97 morphology features as determined by the RSKC algorithm. Features are arranged so weights appear in decreasing order in the weights plot. Top weighted features (>50% max weight) are indicated by the black bar.

The denSNE plots of morphology clusters for the PV-associated genes in S1 (Fig. 32) and V1 (Fig. 33) had similar organizations to the plot of just PV+ cells labelled by ISH(Fig. 7). Both denSNE plots had roughly orthogonal gradients for size and shape features (Figs. 32B & 33B). Examples of the PV-associated cells in the clusters illustrate the range of cell sizes and shapes (Figs. 32A & 33A).

**Figure 32.**
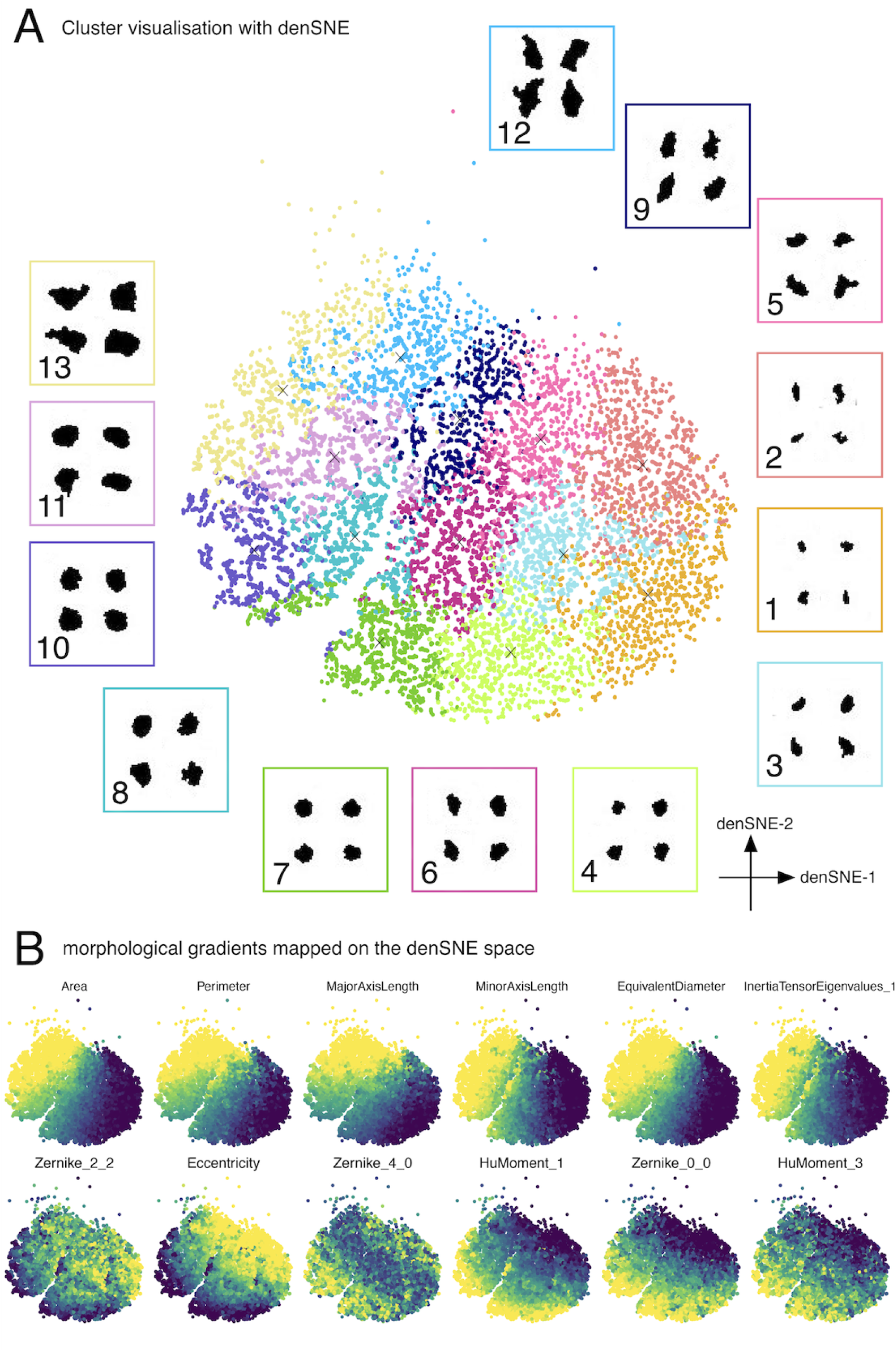
Visualising PV+ gene morphologies for S1-. (A) Morphology space for S1 cells labelled for PV and associated genes visualized using density preserving t-SNE (denSNE). The denSNE algorithm was applied to a transformed dataset where the weights obtained from RSKC were multiplied to their corresponding feature z-scores. Single points are single cells and points are coloured by their cluster assignments. Cluster numbers reflect the progression of median cell sizes (Cluster 1 - lowest median cell size, Cluster 13 - highest median cell size). 4 cells were randomly sampled from each cluster and are shown in the inset squares surrounding the denSNE plot. (B) The denSNE plot in panel A was annotated for a series of six top-weighted size parameters (top row), and six top-weighted shape parameters (bottom row). The colour progression from dark blue to yellow indicates increasing feature values for each morphology feature.

**Figure 33.**
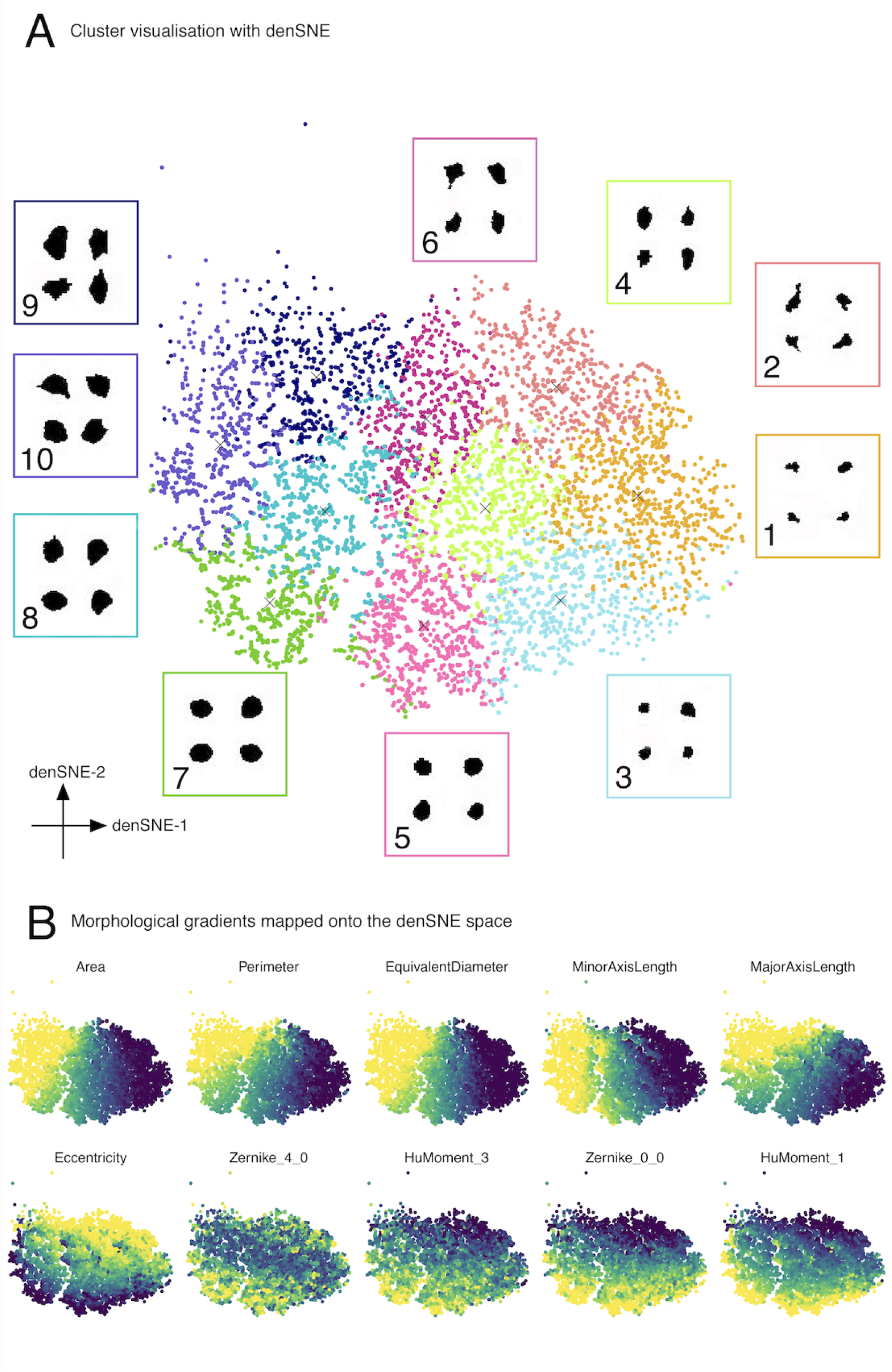
Visualising PV+ gene morphologies for V1-. (A) Morphology space for V1 cells labelled for PV and associated genes visualized using density preserving t-SNE (denSNE). The denSNE algorithm was applied to a transformed dataset where the weights obtained from RSKC were multiplied to their corresponding feature z-scores. Single points are single cells and points are coloured by their cluster assignments. Cluster numbers reflect the progression of median cell sizes (Cluster 1 - lowest median cell size, Cluster 10 - highest median cell size). 4 cells were randomly sampled from each cluster and are shown in the inset squares surrounding the denSNE plot. (B) denSNE plot from panel A, but annotated for a series of six top-weighted size parameters (top row), and six top-weighted shape parameters (bottom row). The colour progression from dark blue to yellow indicates increasing feature values for each morphology feature.

Heatmaps of the morphology clusters and top-weighted features were plotted for each cortical area to identify the morphological features that differentiated among the clusters. The heatmaps had similar organizations in S1 and V1 and were consistent with the previous analyses. For example, clusters were ordered by cell size features and clusters of similar size were separated by the shape features (Figs. 34A & 35A). In both cortical areas, there were three shape motifs: fusiform, pyriform and round, similar to the PV+ cell dataset (Figs. 37B & 38B). In V1, 3 size divisions were identified: small, medium and large (Fig. 38A), whereas S1 had 4 sizes: small, medium, large and extra large (Fig. 37A).

**Figure 34.**
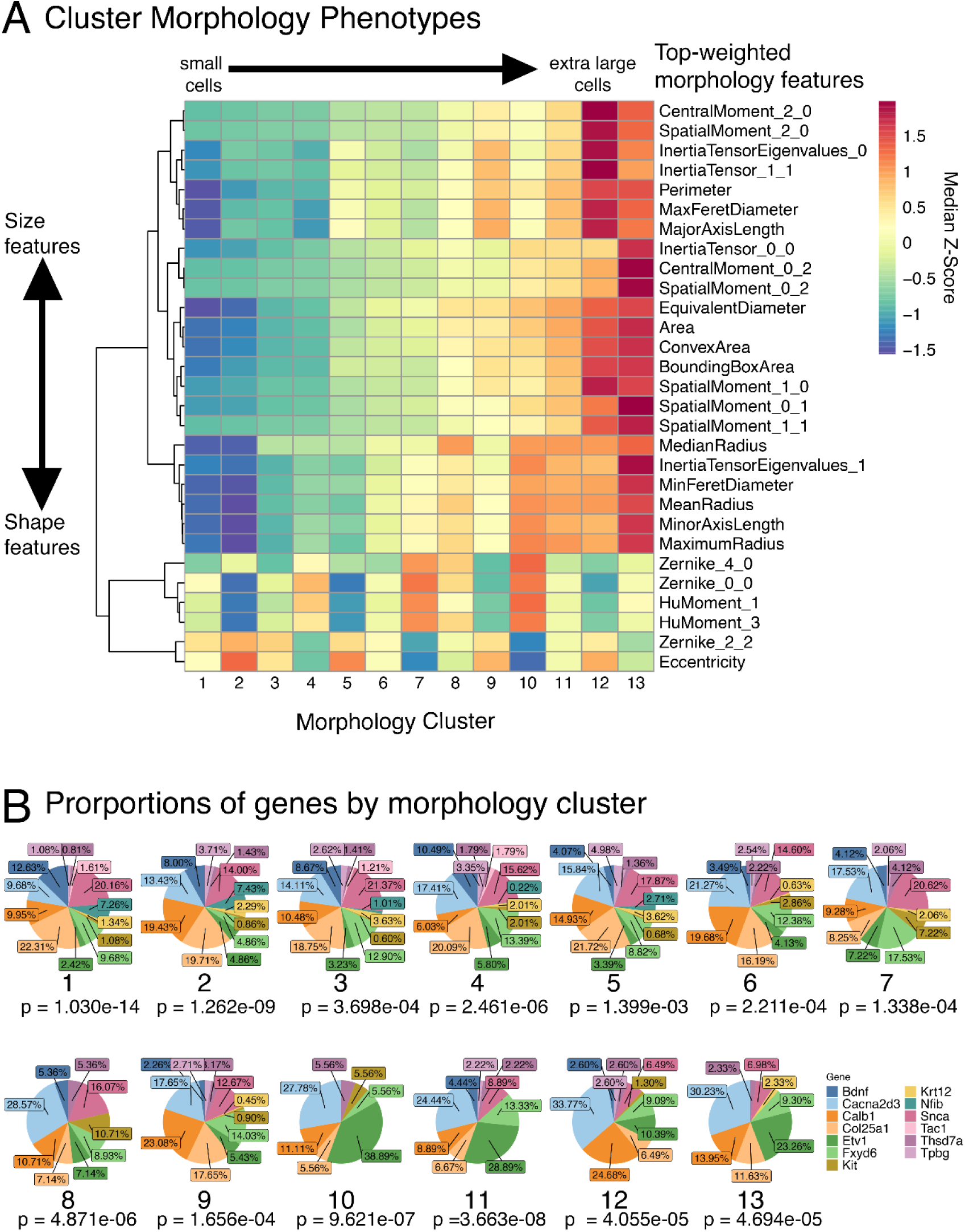
Characterizing morphology clusters for PV genes in S1 -. (A) Morphology phenotype heatmap for the morphology clusters determined for S1. Phenotypes are constructed using the median z-score values for the top-weighted morphology features for each cluster, and organized with a row-wise dendrogram. Cluster numbers reflect the progression of median cell sizes (Cluster 1 : lowest median cell size, Cluster 13 : highest median cell size). (B) Proportions of cells labelled for the 13 genes in each morphology cluster. The proportions were compared to the global proportion of cells labelled for each gene in the dataset. P-values were determined using the chi-square test to assess goodness of fit.

We calculated the proportions of cells labelled for each of the PV-associated genes in S1 and to assess whether specific morphologies are specific to different groups of genes. To compare these to the expected distribution of cells labelled for PV-associated genes, we used the chi-square test for goodness of fit. Across all clusters, the proportions of genes are different from the expected proportions shown in figure 36. In both S1 and V1, the associated genes more readily identify morphologies of PV+ interneurons with smaller median cell sizes (Figs. 34B, 35B). PV-associated genes are more distributed across these clusters, whereas among clusters with larger median cell size, specific genes account for large proportions of the cells labelled for PV-associated genes. Among the large and extra-large size divisions, cluster 9 was the only cluster without a largely different distribution of genes than expected. In V1, similar to S1, most of the PV-associated genes concentrate across the smaller size divisions. The medium and large size divisions display greater restriction to a smaller subset of genes, with the largest proportion of non-PV+ cells in V1 being labelled for Calb1 (p = 5.207e-09). Despite the greater representation of our 13-gene panel across the smaller size divisions, our panel of 13 genes does identify some larger cells among fewer of the genes, with the greatest proportions of cells being labelled for Cacna2d3 in S1 or Calb1 in V1.

**Figure 35.**
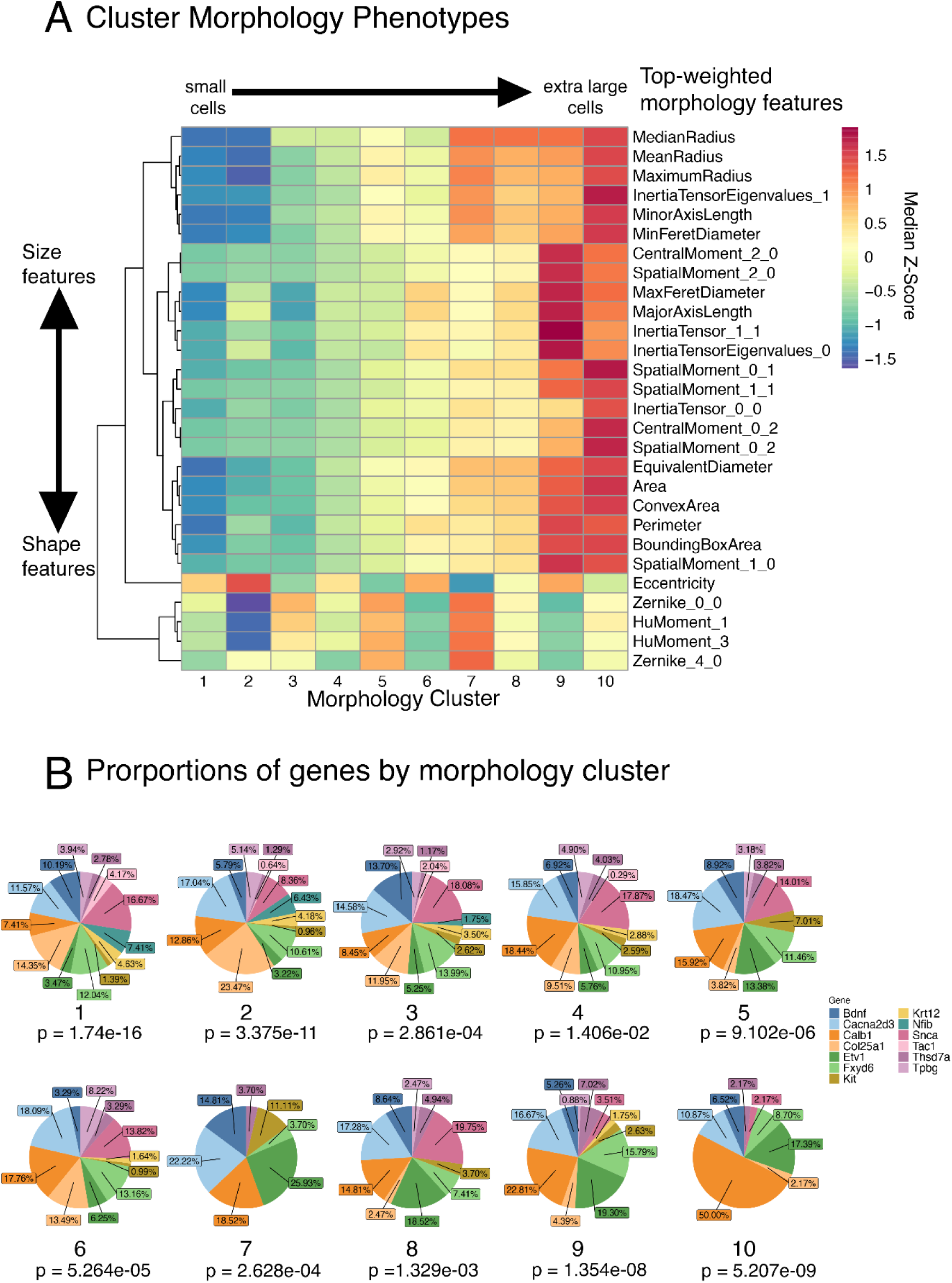
Characterizing morphology clusters for PV+ genes in V1. (A) Using the top-weighted features, a morphology phenotype was constructed for each cluster and organized by row-wise dendrograms. The clusters are ordered and numbered according to the progressions of median sizes (Cluster 1 : lowest median cell size, Cluster 10: highest median cell size). The first branch point on the row-wise dendrogram separates features into size and shape parameters. (B)The proportions of PV-associated genes are shown in pie charts for each morphology cluster. Proportions are compared against the global proportion of cells for each gene, p-values were determined using the chi-square test for goodness of fit.

**Figure 36.**
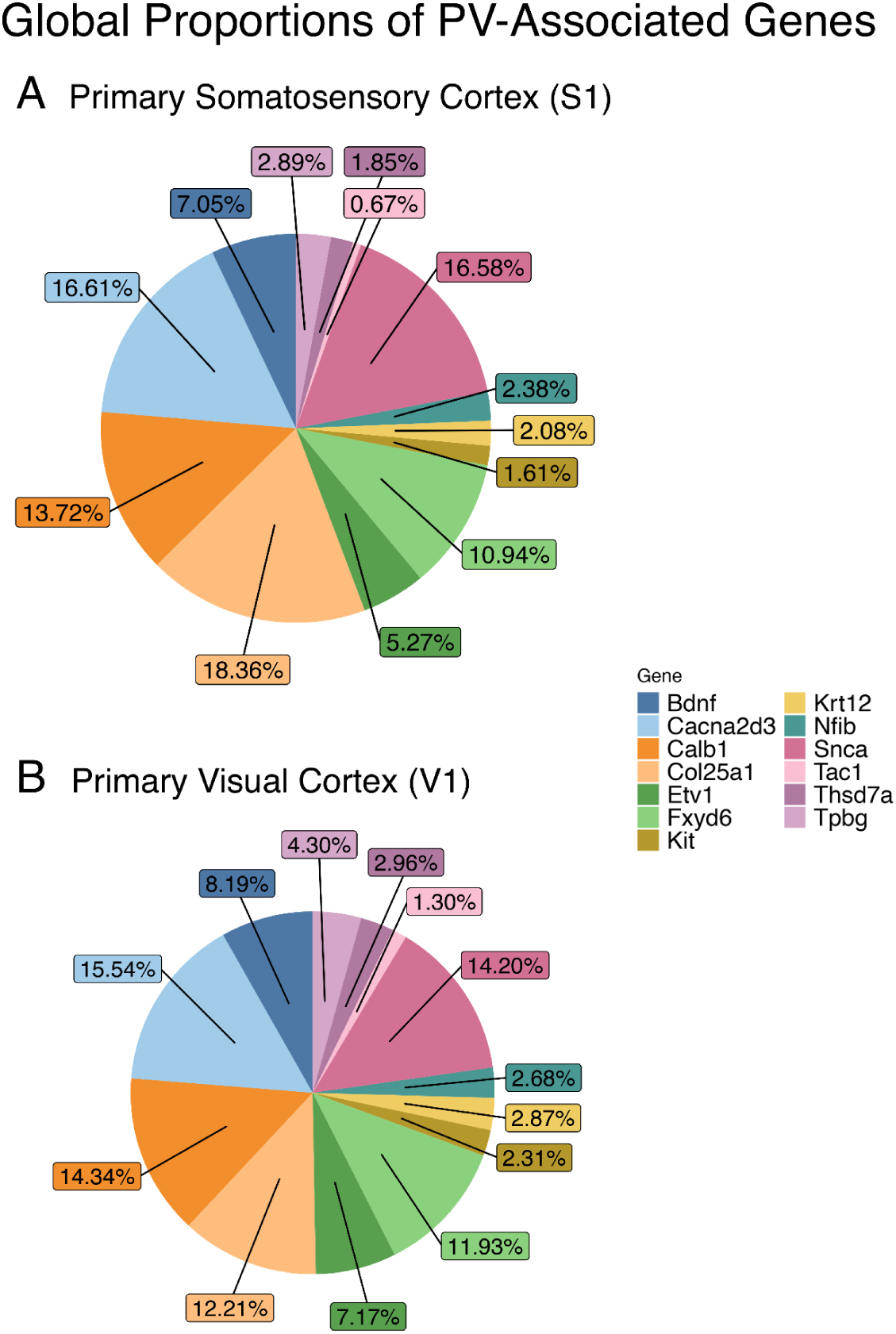
Expected proportions of PV-associated genes for S1 and V1. Pie charts showing the proportion of cells labelled for each PV-associated gene in (A) S1 and (B) V1. These are the proportions of cells taken across the entire dataset of PV-associated genes for S1, and V1 respectively. These are the comparators used in the chi-square tests presented in figure 35.

**Figure 37.**
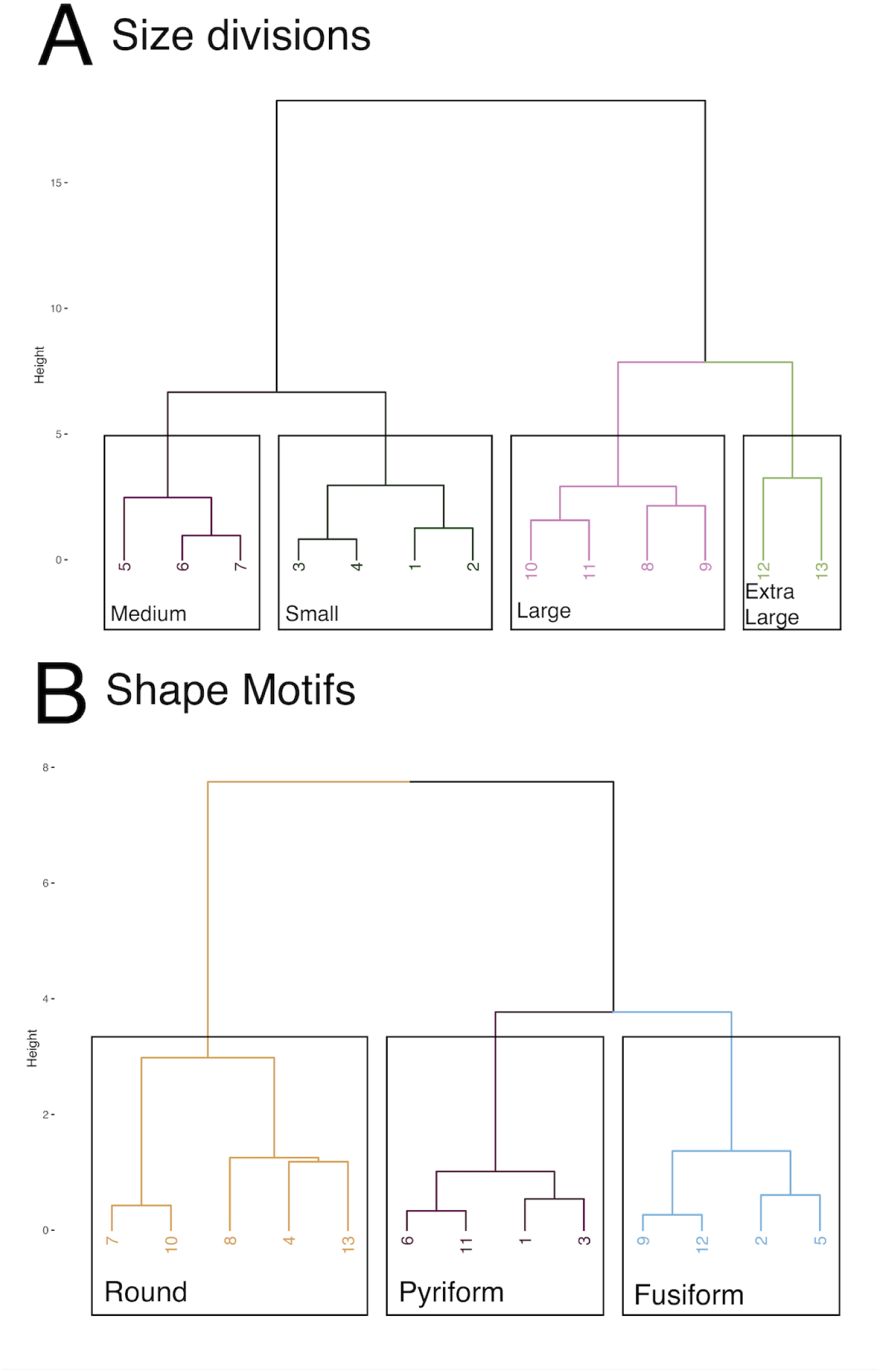
Size divisions and shape motifs for PV+ gene morphology clusters in S1. The median z-scored values for the top-weighted size and shape parameters were clustered separately using unsupervised hierarchical clustering (Ward.D2 method) to identify the groups of clusters in S1 with similar sizes, and similar shape characteristics. Dendrograms showing the grouping of morphology clusters into (A) four size divisions, and (B) three shape motifs. The number of size divisions and shape motifs were determined using the elbow method.

**Figure 38.**
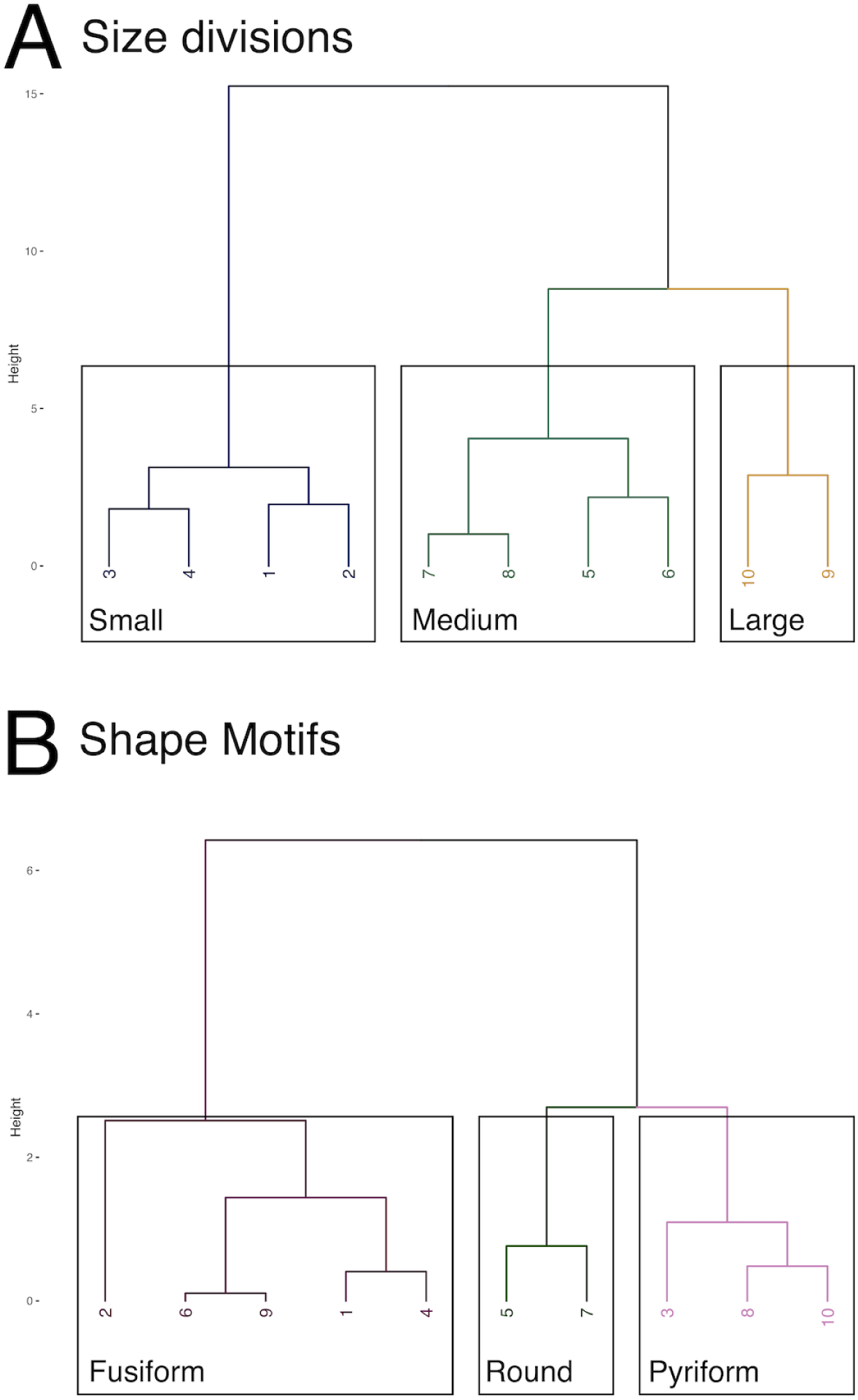
Size divisions and shape motifs for PV+ gene morphology clusters in V1. The median z-scored values for the top-weighted size and shape parameters were clustered separately using unsupervised hierarchical clustering (Ward.D2 method) to identify the groups of clusters in V1 with similar sizes, and similar shape characteristics. Dendrograms showing the grouping of morphology clusters into (A) three size divisions, and (B) three shape motifs. The number of size divisions and shape motifs were determined using the elbow method.

To compare the laminar location of the different PV-associated gene in each morphology cluster, we mapped the profiles for each gene in the morphology clusters (Figs. 39 & 40). We only plotted data for genes where there were at least 10 cells in the morphology cluster labelled for that gene. Among the small size divisions, in S1, PV+ cells display a unimodal, infragranular peak across layers 5 and 6A. The medium and larger size divisions display a different distribution with two peaks. A smaller peak across the supragranular and granular layers (L2/3-4), and a larger peak across infragranular data. In V1, PV+ cells follow a similar distribution across layers among small and medium size divisions, the large size division displays a sustained, more uniform density of cells across the layers. These findings indicate that different PV+ morphologies are found across different layers, which alludes to transcriptomic differences among PV+ cells (Gouwens et al., 2019).

**Figure 39.**
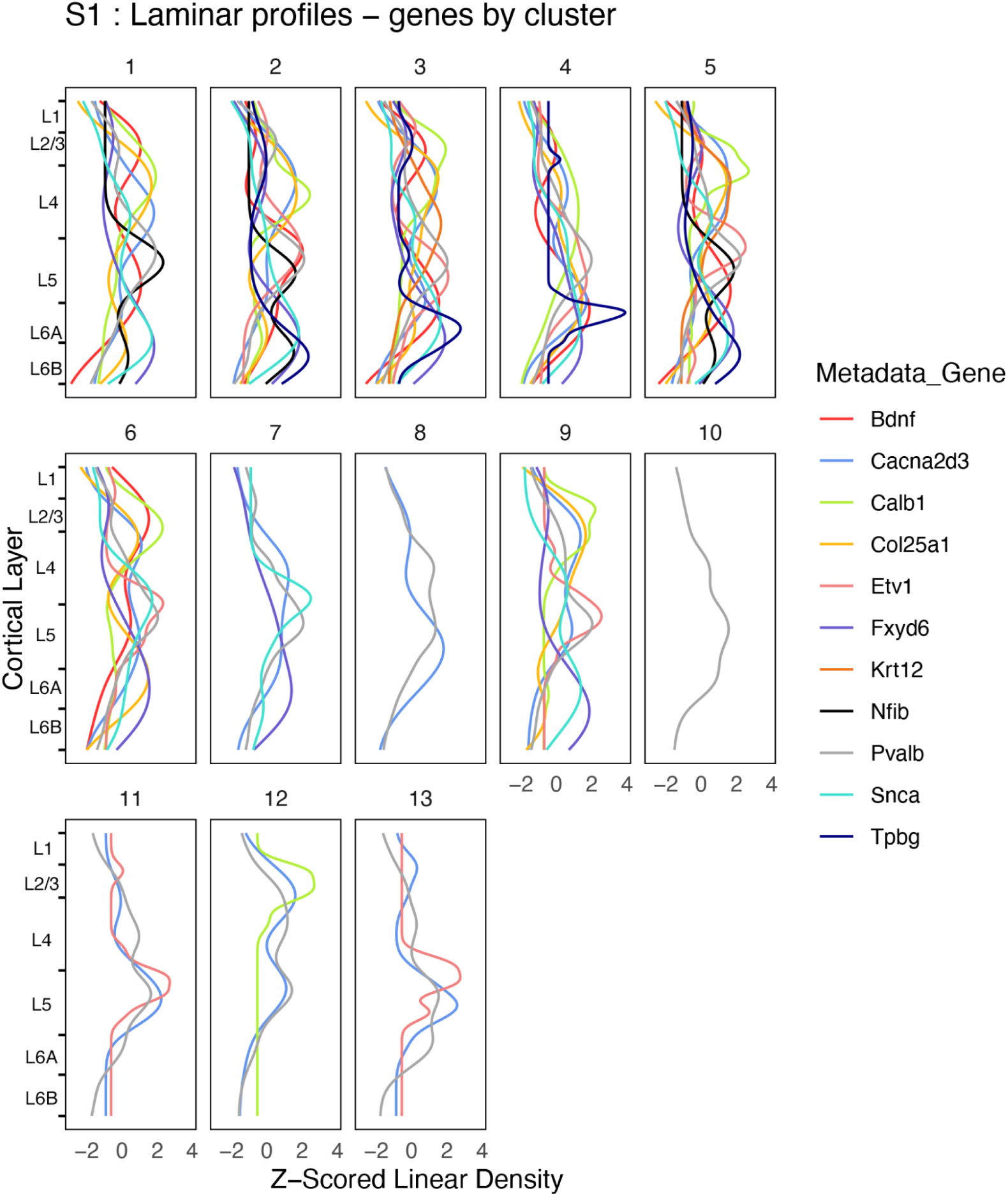
Laminar profiles for PV-associated genes : genes by morphology cluster for S1. In each morphology cluster, the Z-scored linear density profile was plotted along the standard cortical depth for each gene. In each morphology cluster, curves were plotted for a gene if at least 10 cells were labelled within a morphology cluster.

**Figure 40.**
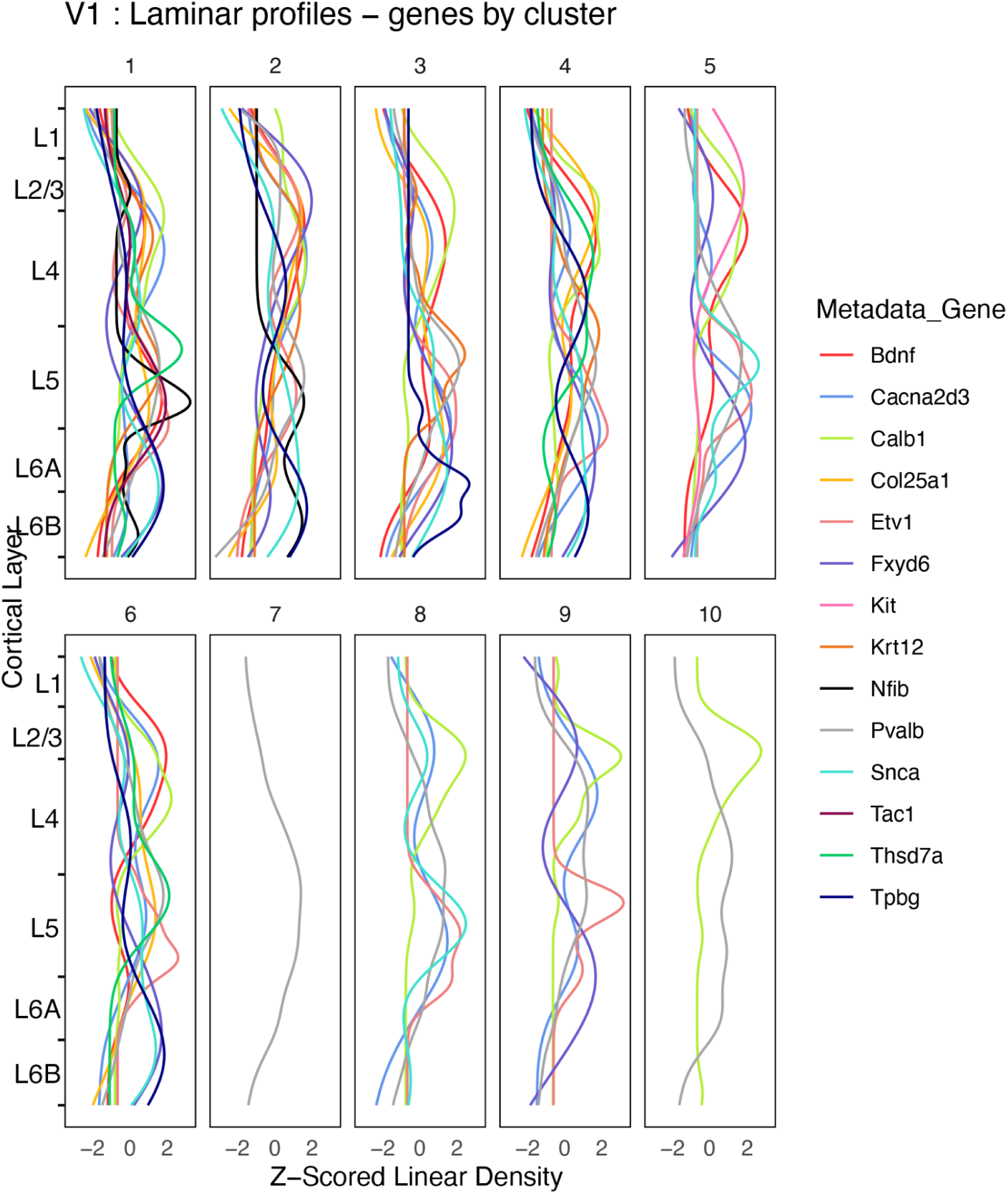
Laminar profiles for PV-associated genes : genes by morphology cluster for V1. In each morphology cluster, the Z-scored linear density was plotted along the standard cortical depth for each gene. In each morphology cluster, singe curves were plotted for a gene if at least 10 cells were present within the morphology cluster.

In both cortical areas, there is a greater representation of our panel of 13 genes across the small size divisions along with the smaller end of the medium size division (clusters 1-6 in S1 and V1). These distributions indicate a broad laminar distribution for these morphologies, however within each of these clusters, the laminar distributions of individual genes are separating across different groups of layers. For cluster 12 in S1, the laminar distribution of Cacna2d3 overlaps with the infragranular peak of PV+ cells (Fig. 39). Whereas in cluster 8, the distribution of PV+ cells and Cacna2d3+ cells appear out of phase with one another. In cluster 12, a subset of PV+ cells and Cacna2d3+ cells share a common morphology with comparable laminar distributions. This would allude to Cacna2d3+ cells in cluster 12 being of a specific PV+ T-type that is not shared with the Cacna2d3+ cells in cluster 8.

In V1, among the large size divisions, there are fewer genes represented across individual morphology clusters compared to small size divisions. This is consistent with what is seen in S1. However, among the large size division, we can see the lack of alignment between PV and several of the associated genes. For example,in cluster 10, there is a low PV+ cell density across the supragranular layers, where Calb1+ cell density is high (Fig. 40). This combination of common morphology but lack of alignment with th PV+ laminar profile could suggest that the Calb1+ cells identified in cluster 10 do not share the transcriptomic correlates of the PV+ cells identified in the same cluster.

Notably, Etv1 was found across several morphology clusters, despite its characteristic restriction to layer 5 (Figs. 39 & 40). This finding establishes that several cell morphologies can exist among cells labelled for a single gene, and that for a single cortical layer, there is still a distribution of cell morphologies that are present.

Together, these analyses of PV-associated cells highlight that cell morphology can add another dimension to studying the diversity of PV+ cells. Our approach uncovers robust morphological complexity, and establishes that subtle morphological differences for a sufficiently large number of morphology parameters, for a sufficiently large number of cells, can yield pronounced differences in size and shape that distinguish among classically homogeneous cell classes.

## Discussion

Our approach is a novel data-driven framework to analyze cell morphologies. By obtaining high-dimensional cell morphology data, our approach aligns analyses of neuroanatomical data with approaches applied in single-cell transcriptomics studies. Our image analysis procedure constructs a high-dimensional dataset that quantifies 97 morphometric parameters per cell. We leverage unsupervised clustering and dimensionality reduction to cluster cells based on morphologic patterns and visualize these clusters in their high-dimensional morphology space. Our morphology data extend beyond classic morphological data in neuroanatomical studies, as we also quantify the shapes of single cells and incorporate shape information together with size in defining our morphology clusters. Our inclusion of Zernike moments serves as an important shape parameter, as these moments capture complex shape features, including aspects of cell geometry and symmetry (Pincus and Theriot, 2007). Because of their invariance to scale and rotation, Zernike moments provide a basis to consistently quantify the morphologies of PV+ interneuron cell bodies, capturing salient shape features that differentiate among different groups of cells. Our use of the RSKC and denSNE algorithms served as powerful ways to both define morphology clusters and interrogate the relative importance and contribution of single morphometric features. RSKC is particularly powerful for our study as feature weights identify which morphometric features most strongly separate PV+ cells(Kondo et al., 2016). The weights were also important in characterizing the aspects of cell shape that further distinguish among cells of similar size. We incorporated the weights from RSKC into the denSNE algorithm to visualize the clustering result and how these morphometric parameters organize cells in their high-dimensional space (Balsor et al., 2021). By combining both high-dimensional data and an unsupervised cluster analysis approach, we provide a more nuanced, reproducible characterization of PV+ interneuron morphologies by quantifying aspects of cell morphology that have remained beyond the reach of classic morphological analyses (Lingley et al., 2018; Bjerke et al., 2021). Our approach bridges classical neuroanatomical techniques with modern data-driven analyses. This novel and accessible approach allows us to construct comparable data structures and apply a highly sensitive approach to characterize single PV+ interneuron morphologies in mouse S1 and V1.

However, despite these strengths, important limitations must be considered. All of the Allen Mouse ISH Database mice are male, so we cannot comment on potential sex differences in PV+ cell morphology. Moreover, our study compares S1 and V1 and does not account for the presence of additional PV+ cell morphologies that could be present in other cortical or subcortical areas. These additional morphologies, which were not profiled in this analysis, may differ from those we identified here. As a result, our analysis does not necessarily represent a complete repertoire of brain-wide PV+ interneuron morphologies. In our integrated data analysis, we combined ISH and whole-cell fill data. Despite the overlap of size distributions for cells obtained from either labelling technique, the filled cells we analyzed did not represent the full range of sizes identified in the ISH data, as they mostly fell into clusters with larger median cell sizes. This is due in part to the low number of filled cells (n=36) and the fact that small cells are more difficult to fill and record from (Ritzau-Jost et al., 2021). Consequently, it is unknown from the Allen Cell Types Database data whether smaller cells are underrepresented due to technical challenges with the whole-cell fills or whether tissue processing steps affect the apparent sizes of cells across the full-size range.

Investigating cell morphology has garnered attention, especially in other cell types where dynamic morphological change is associated with changes in functional state (Nimmerjahn et al., 2005). For example, in microglia, brain-resident macrophages, morphological changes associated with Alzheimer’s disease have been shown to follow sexually dimorphic patterns, with different disease-related changes occurring in females than males (Guillot-Sestier et al., 2021). Additionally, in animal models of Alzheimer’s disease, microglial morphological change has been shown to occur earlier in females (Colombo et al., 2022). Regarding our cluster analysis, our results provide intriguing insights into the subtle morphological differences among PV+ cells; however, it remains unknown whether direct functional differences are associated with the morphologies we have uncovered.

We also explored the relationship between gene expression and cell morphology to interrogate how single-cell morphologies recapitulate the transcriptomic patterns seen among PV+ interneurons. It is important to recognize that our gene panel is limited as it does not represent the entire panel of marker genes associated with PV+ transcriptomic types (Tasic et al., 2016). These were the marker genes of good quality that were available in the Allen Institute mouse ISH database. As we found our 13-gene panel was more represented among smaller cell morphologies, an expanded gene panel for more marker genes stated across major cell type taxonomy studies would be needed to canvas the range of morphological types we have discovered. We found similar gene distributions in S1 and V1, which suggests alignment with our first analysis of cell morphologies of PV+ interneurons from S1 and V1 together. This has identified no specific morphologies distinguishing V1 from S1, but common morphologies occur in different proportions. As V1 has been identified as transcriptomically distinct from other cortical areas (Jorstad et al., 2023), our initial results suggest that morphological differences may distinguish V1 from other cortical areas. Although we did not observe distinct morphologies between S1 and V1, it is possible that the distribution of PV+ morphological types in V1 differs from other cortical areas or that specific morphologies outside of V1 are also present. This is an important avenue for our approach to assess directly by analyzing data from several cortical areas and comparing the distributions of different cell morphologies identified across other cortical areas. Capturing the distributions of cell morphologies across several cortical areas would also further align our approach with the brain-wide approaches used to profile the transcriptomic landscapes of several brain areas (Langlieb et al., 2023; Zhang et al., 2023).

Quantifying and analyzing cell morphologies has been established as a powerful quantitative tool for understanding how cell structures change in response to complex biological processes like aging, disease and dysfunction. Cell morphology studies have revealed novel morphological biomarkers that serve as the structural correlates of age, sex, and disease-specific morphology patterns for different cell types (Prasad and Alizadeh, 2019; Hagemann et al., 2021; Colombo et al., 2022; Kamat et al., 2024). Assessments of morphological change of PV+ cells in disease need to be further elucidated as current approaches have emphasized the dendritic arbour together with the spatial networks of axonal fibres (DeFelipe, 1999; Pierri et al., 1999; Song et al., 2023), with less attention given to the size and shape of the cell body. Given the cell body’s role in coordinating cellular energy and the high metabolic demand of fast-spiking PV+ interneurons, altered cell body morphologies could underscore the heightened vulnerabilities of PV+ interneurons in neuropsychiatric disorders where cell damage due to metabolic stress impacting PV+ interneurons (Whittaker et al., 2011; Steullet et al., 2017; Ruden et al., 2021). Our approach responds to the need for a more comprehensive approach to capturing the morphological diversity of PV+ cells. Our ability to resolve and quantify the morphological patterns among the cell body is a powerful additional insight into PV+ interneuron structure. It provides novel structural dimensions to explore in disease states. Our approach provides novel insight into the structural changes of PV+ cells across all major neuronal compartments, and has the potential to enrich our understanding of how morphological changes impact PV+ cell dysfunction in disease.

We have developed and applied a novel approach to quantitative neuroanatomy that readily obtains high-dimensional cell morphology data for large numbers of individual cells. The datasets we can obtain are comparable in structure and dimension to those generated in single-cell RNA sequencing studies and offer a powerful new platform to explore cell morphology at a scale previously unattainable from classic neuroanatomical image data. Classic approaches to defining cell morphologies rely heavily on expert annotations, which are largely descriptive and often labor-intensive. Consequently, this approach becomes time-intensive and impractical if applied to larger datasets, in addition to issues with reproducibility across datasets. Our consistent image processing steps, automated morphology quantification, and unsupervised clustering techniques overcome the subjectivity and time intensity previously associated with analyses of cell morphology. Our use of open software platforms also helps make large-scale, high-throughput workflows accessible to a wider range of researchers. In sum, our analytical framework is a powerful tool to unlock cell morphologies at an unprecedented scale, with the potential to afford novel insight into the dynamic changes in cell morphology during development, disease progression, or in response to novel treatments.

